# Rapid Antimicrobial Resistance Decline Coupled with Microbial Community Shifts in Sewage Polluted River Mesocosms

**DOI:** 10.64898/2026.09.08.750159

**Authors:** Arun Kashyap, Vikas Sonkar, David W. Graham, Jan-Ulrich Kreft, Shashidhar Thatikonda

**Author notes:** Corresponding authors: Jan-Ulrich Kreft, and Shashidhar Thatikonda.

## Abstract

Antimicrobial resistance poses a global health threat, yet the fate of antimicrobial resistance genes (ARGs) and microbial communities from wastewaters in receiving rivers is rarely quantified. This study investigated the degradation kinetics of ARGs in sewage-polluted river water mesocosms and under sulfamethoxazole and copper stress, separately and in combination. Seven clinically relevant ARGs (*bla*_CTX-M_, *bla*_NDM_, *qnrS*, *sul2*, *tetW*, *ermF*, and *aph(3’’)-Ib*), a mobile genetic element marker (*intI1*) and bacterial marker genes (*uidA*, *16S rDNA*) were quantified by qPCR. Bacterial community changes were monitored using 16S rDNA amplicon sequencing. All target genes decayed with approximately first-order kinetics in all conditions, with *qnrS* (half-life: 8.0 h) and *bla*_CTX-M_ (8.4 h) declining most rapidly and *intI1* (32.5 h) and *sul2* (46.5 h) declining most slowly. Ordination showed that ARG composition shifted primarily over time rather than by antibiotic/metal treatment. Bacterial communities shifted from initially *Campylobacterota* dominated (≥ 90%) to *Pseudomonadota*, *Bacillota*, and *Actinomycetota* dominated communities over 168 h. Quantitative microbiome profiling showed a 95% reduction in ASV richness. PICRUSt2 analysis suggested progressive enrichment of aerobic and several metabolic pathways, as the mesocosms transitioned from highly polluted anoxic conditions to oxygenated conditions. Shifts in microbial community composition were coupled with changes in the resistome as identified by Procrustes analyses (r = 0.841, p = 0.001) and other analyses. Together, results show that river self-purification can rapidly reduce ARG loads, at least in warmer climates. This study provides approximate kinetic parameters for mathematical models to predict the fate of ARGs in rivers.

## 1. Introduction

Antimicrobial resistance (AMR) is a growing global One Health crisis, with aquatic ecosystems recognized as key reservoirs and transmission routes of antimicrobial resistance genes (ARGs) and antimicrobial-resistant bacteria (Graham et al., 2011; Singh et al., 2024; Liu et al., 2025). Raw or treated wastewater discharges, from communities, hospitals and agriculture, containing antibiotic residues, ARBs and ARGs, are frequently reported to pollute urban rivers (Berendonk et al., 2015; Kashyap et al., 2023). These contaminated waters contain persistent antibiotics, including sulfonamides, fluoroquinolones and macrolides, at concentrations that may exert selective pressures for resistance (Johnson et al., 2015; Larsson and Flach, 2021; Zhang et al., 2024; Yu et al., 2025). Sulfamethoxazole (SMX) is one of the most widely used sulfonamide antibiotics and among the most frequently detected antibiotics in surface waters. Untreated wastewaters typically contain SMX at concentrations of approximately 1–200 µg/L (Le et al., 2016; Quoc Tuc et al., 2017; Phonsiri et al., 2019; Borsetto et al., 2021), whereas treated effluents generally contain residual concentrations of approximately 0.01–1 µg/L (Michael et al., 2013; Tran et al., 2018). Nevertheless, both concentration ranges remain capable of imposing selective pressure for AMR, creating localized “hotspots” within river systems (Michael et al., 2013; Sabri et al., 2020; Yan et al., 2025).

Urban wastewater also introduces non-antibiotic contaminants, particularly heavy metals, which can facilitate AMR emergence. Copper (Cu) is among the most ubiquitous metal polluting urban rivers owing to industrial, urban, and agricultural activities (Jamwal et al., 2022; Gupta et al., 2023; Tiwari et al., 2025). Unlike organic antibiotics, metals do not degrade, although their bioavailability may decrease through sorption and redox reactions. Consequently, they persist in environmental compartments and exert continuous selective pressure on microbial communities (Xavier et al., 2019; Gupta et al., 2022). Co-exposure to Cu and SMX can significantly perturb microbial community composition, reduce diversity, alter carbon and nitrogen cycling pathways (Liu et al., 2016; Li et al., 2023; Narciso et al., 2024) and synergistically enhance the relative abundance of *sul2* and *intI1* (Wu et al., 2022; Wang et al., 2024). Quantifying these co-selection processes is essential for parameterizing mathematical models that predict ARG persistence and dissemination in aquatic environments (Arya et al., 2021).

Most experimental studies have focused on the effects of long-term exposure (weeks to months) to antibiotics and metals, demonstrating shifts in microbial community composition, reductions in diversity, and enrichment of resistance determinants such as *sul2* and *intI1* (Radke et al., 2009; Patrolecco et al., 2018; Deng et al., 2020; Li et al., 2023; Narciso et al., 2024; Schuijt et al., 2024; Zhang et al., 2024; Jia et al., 2025). Additionally, there is limited understanding of how environmentally relevant, sub-inhibitory concentrations of Cu and SMX influence the early kinetics of ARG proliferation and decay, horizontal gene transfer, and microbial community restructuring during the initial stages of exposure (Engemann et al., 2008; Li et al., 2015; Borsetto et al., 2021).

Environmental conditions also affect the persistence and dissemination of AMR. Physicochemical parameters, including pH, temperature, dissolved oxygen (DO), chemical oxygen demand (COD), total organic carbon and nutrient availability, influence microbial community assembly and ARG persistence in aquatic ecosystems (Chen et al., 2019; Yang et al., 2020; Jampani et al., 2024). For example, TOC can sustain heterotrophic bacterial activity and indirectly enhance ARG stability (Jia et al., 2017; Yang et al., 2020), while lower DO and acidic pH have been associated with increased ARG abundance in sediments and biofilms (Ju et al., 2019; Zhu et al., 2025). Time-series investigations further suggest that these responses are often characterized by an initial increase in ARG abundance and microbial community restructuring, followed by progressive attenuation and stabilization over time. (Y. Y. Liu et al., 2024). Understanding these early responses is therefore essential because they may influence whether AMR becomes established or diminishes within aquatic ecosystems, potentially marking critical transition periods in ecological response trajectories (Lee et al., 2022; Leão et al., 2023). Addressing this knowledge gap requires experimental systems that simulate realistic river conditions.

Mesocosms provide an effective bridge between simplified laboratory experiments and natural river systems by maintaining ecological complexity while allowing controlled manipulation of environmental variables (Wimpenny, 1988; Battin et al., 2003; Knapp et al., 2008, 2010; Walters et al., 2010; Lehmann et al., 2016; Borsetto et al., 2021). Batch mesocosms are particularly well-suited for quantifying ARG decay kinetics because no additional contaminant inputs occur during the experiment, allowing temporal changes in ARG abundance to be attributed to internal ecological and physicochemical processes rather than to continuous external loading (Baker et al., 2022). Their larger volume also captures greater spatial heterogeneity and microbial diversity than conventional laboratory microcosms, generating experimentally derived parameters that can improve predictive models of environmental AMR dynamics (Baker et al., 2022).

This study used batch mesocosms to investigate the fate of ARGs in wastewater-polluted river water under environmentally realistic conditions and in the absence of additional pollution inputs to quantify ARG decay kinetics. Specifically, we examined whether temporal changes in ARG abundance were associated with microbial community succession, microbial diversity, and physicochemical conditions, including dissolved oxygen. We further evaluated the effects of environmentally relevant additions of SMX and Cu, with and without river sediment, on these processes. The resulting kinetic parameters provide a basis for improving predictive models of ARG fate in river systems and strengthening environmental AMR risk assessment.

## 2. Materials and methods

### 2.1 Sampling polluted river water and sediment

Water samples were pumped from the riverbank into a tanker of 5,000 L capacity and sediment samples were grabbed in duplicate at 0-15 cm depth for the mesocosm experiments during the dry season (19 February 2024) from a sewage-impacted site of the Musi River, located within Hyderabad city (Nagole Bridge: 17°22′56.43″ N, 78°33′27.51″ E) in Telangana, Southern India (**Supplementary Figure S1**).

### 2.2 Mesocosm experiment

The temporal dynamics of water quality, microbial communities and genes were studied in six 300 L cylindrical mesocosm reactors (Diameter: 80 cm; Height: 60 cm; and Operating volume: 250 L), each representing the following simulated conditions: Polluted Musi river water only (MW); MW with SMX only (MW.SMX, 6.4 mg/L, a subinhibitory concentration set to 0.1× the experimentally determined MIC of 64 mg/L for the M24 *E. coli* strain (GenBank accession number MZ63432) (Palkovicova et al., 2022); MW with Cu only (MW.Cu, 0.05 mg/L, environmental concentration), MW with SMX (6.4 mg/L) and Cu (0.05 mg/L) as combined stressors (MW.SMX.Cu), MW with SMX (6.4 mg/L) and Cu (0.05 mg/L) in the presence of sediment (MW.Sed.SMX.Cu) and a control with only tap water (TW). The reactor set-up with aeration (4L/min per tank) and LED lights (∼6,000 lumens per tank) and other operating parameters is described in **Supplementary Figure S2**. These tanks were placed in an open space under the roof of the Institute’s Civil Engineering Department building with an average temperature of 26.6±2 °C and humidity around 30±10%.

Water quality parameters (WQPs) including temperature, pH, DO, turbidity, and conductivity were measured under *in situ* conditions using a portable multimeter with smart probes (Orion Star A329, ThermoFisher Scientific, USA). Mesocosms were sampled at 0, 4, 16, 24, 72 and 168 h for determining WQPs including COD, TOC, ammonia, and ortho-phosphate using the *APH*A standard laboratory procedures (**Supplementary Table S1**). In addition to water samples, sediment samples from the MW.Sed.SMX.Cu mesocosm were collected at 0, 24, 72, and 168 h for further testing of physicochemical parameters **(Supplementary Table S2)**. Unlike most previous studies, the mesocosm reactors were deliberately operated without a stabilization period to capture fast dynamics of WQPs, ARGs and bacterial community structure under the treatment conditions.

### 2.3 Bacterial enumeration

*Escherichia coli* was selected as an indicator bacterium for assessing sewage contamination because of its well-documented ability to survive, acclimate, and persist in environmental matrices over extended periods. Moreover, it is a widely accepted marker of fecal pollution originating from both human and animal sources and is capable of undergoing HGT, making it a relevant sentinel organism for tracking antibiotic resistance in contaminated aquatic environments (Baker et al., 2022).

The bacterial enumeration in water and sediment samples was performed as described in Sonkar et al. (2025): *E. coli* was enumerated on modified OECD Synthetic Sewage (OSS) medium supplemented with 24 mg/L X-Gluc (5-Bromo-4-chloro-3-indolyl-β-D-glucuronide) (HiMedia, India) (Frampton et al., 1988) to mimic the nutrient conditions of polluted rivers. The OSS medium (OECD, 2001; O’Flaherty and Gray, 2013; Tanaka et al., 2014) was supplemented with 100 mg/L sodium dodecyl sulphate (Birkeland and Steinhaus, 1939) to inhibit Gram-positive bacteria. The appearance of blue colonies on the modified agar plates showed β-Glucuronidase activity of *E. coli* colonies. To quantify SMX-resistant *E.* coli, the bacteria were enumerated on OSS with X-Gluc supplemented media containing 6.4 mg/L SMX (OSS.SMX, TCI, Japan). All plates were incubated at 44 to select for human gut-associated bacteria, and triplicate colony counts were recorded after 24 h.

### 2.4 DNA extraction and gene quantification

DNA was extracted from water and sediment samples collected at 0, 24, 72 and 168 h from mesocosms using the DNeasy® PowerWater® Kit or the DNeasy® PowerSoil® Pro Kit (Qiagen, Germany), respectively, following the manufacturer’s guidelines. Water samples were filtered through a sterile 0.2 µm polyether-sulfone membrane (Diameter: 47 mm; Pall Corporation, USA) until it clogged in order to obtain sufficient biomass, and the filtrate volume was recorded. Filter membranes were stored at –20 before DNA extraction. Duplicates of extracted DNA were pooled in sterile Eppendorf tubes to minimize extraction-level variability prior to downstream molecular analyses.

The purity (A260/A280 ratio: 1.8–2.1) of the extracted DNA was assessed using a NanoDrop™ 2000/2000c spectrophotometer (ThermoFisher, USA) and the concentration was measured using a Qubit™ dsDNA HS assay kit (Invitrogen, USA) on a Qubit™ 4 fluorometer (ThermoFisher, USA), following the manufacturer’s guidelines.

The absolute abundance of total bacteria (*16S rDNA*), the *E. coli* marker gene *uidA*, the MGE class I integron (*intI1*) and seven ARGs conferring resistance to aminoglycosides (*aph(3’’)-Ib*), carbapenems and third-generation cephalosporins (*bla*_NDM_ and *bla*_CTX-M_), fluoroquinolones (*qnrS*), macrolides (*ermF*), sulfonamides (*sul2*) and tetracycline (*tetW*) were quantified by the CFX Opus 96 quantitative PCR (qPCR) system (Bio-Rad, USA) using primers described previously (Sonkar et al., 2025) and listed in **Supplementary Table S3**. These ARGs were selected based on Berendonk et al. (2015) and to represent resistance against antibiotics commonly sold and consumed in Hyderabad, India (Sonkar et al., 2026). The DNA samples were diluted 1:10 prior to qPCR analysis to minimize inhibition. All reactions were performed following the protocol described in (Sonkar et al., 2025). The amplification efficiencies and R^2^ values for the target genes were 97.99 ± 2.88% and 0.997 ± 0.002, respectively.

### 2.5 Microbial community analysis by 16S rRNA amplicon sequencing

The DNA extracted from water and sediment samples collected at 0, 24, 72, and 168 h from each mesocosm was also analyzed for the microbial community structure. The hypervariable V3–V4 region (341F-806R) (Kozich et al., 2013) of the *16S rRNA* gene was sequenced by Gujarat Biotechnology Research Centre, Gujarat, India (Gajjar et al., 2026) using the Illumina NovaSeq 6000 platform. The raw 16S amplicon sequencing data generated in this study have been deposited in the NCBI Sequence Read Archive (SRA) database under BioProject accession number PRJNA1512206.

Raw sequences were processed following Ott et al. (2021). Briefly, the read quality assessment and trimming were performed using FastQC v0.12.1 and fastp v0.23.4, respectively. Quality-filtered sequences were analyzed with QIIME2 v.2024.10.1 (Bolyen et al., 2019), with reads denoised into amplicon sequence variants (ASVs) using DADA2 (Callahan et al., 2016) and assigned up to genus level using the MiDAS reference database v5.3 (Dueholm et al., 2024). All taxonomic-level assignments were reported according to the MiDAS taxonomy throughout all downstream analyses. The resulting taxonomy data was exported for downstream analysis in R (v4.5.1) using the *phyloseq* (v1.52.0) and *vegan* (v2.5–7) packages.

### 2.6 Data analysis

All data were analyzed in R v4.5.1 (R Core Team, 2023) using the *vegan* (v2.5–7), *phyloseq* (v1.52.0), ggplot2 (v4.0.2) and tidyverse (v2.0.0) packages. Due to the unreplicated experimental design, statistical analyses were focused on decay kinetics rather than hypothesis testing. Technical duplicates were averaged to a single mean value per Condition × Time point prior to model fitting. The reported SE and 95% CI for rate constant are derived directly from the fitted linear model’s coefficient standard errors, not from the variability among technical replicates. Time series were fitted to first-order kinetics to calculate the rate constant (k), with its standard error (SE) and the related half-life (t_1/2_) (Brown et al., 2020) using the following equations:

First order decay kinetics

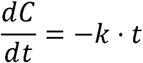

Solution:

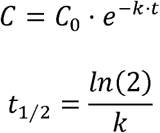

ARG (log -transformed) trajectories across all five treatments at four major time points (0, 24, 72, and 168 h; n = 20 average samples) were analyzed by principal component analysis (PCA) based on Aitchison distances. Distance based Redundancy analysis (db-RDA) was performed on the centered log-ratio (CLR)-transformed ARG abundances.

Quantitative Microbiome Profiling (QMP) was applied following Ott et al. (2021) to generate absolute abundance estimates independent of sequencing depth variation. For each sample, the rarefaction depth was scaled by the ratio of sequencing reads to absolute cell density (cells mL^-1^) derived from *16S rDNA* gene qPCR copy numbers corrected by an average 16S gene copy number of 4.1 copies per genome. The resulting absolute abundance tables, expressed in units of cells mL^-1^ per amplicon sequence variant (ASV), were used for all downstream community composition analyses and diversity estimations. Alpha diversity was determined using the unified Hill number framework. Prior to Hill number calculation, ASV count tables were rarefied to an even sequencing depth of 11,427 reads per sample. Beta diversity was assessed via principal coordinates analysis (PCoA) of Bray-Curtis’s dissimilarities. Predicted functional profiles of bacterial communities were generated using PICRUSt2 based on 16S rDNA gene amplicon data to assess temporal changes in microbial functions across mesocosm treatments. Procrustes analysis was used to compare the similarity between bacterial community composition and ARG ordination patterns across different taxonomic levels using Aitchison distance-based ordinations. Mantel tests were performed using Aitchison distance matrices from CLR-transformed bacterial community and ARG datasets to evaluate the relationship between microbial community and resistome dissimilarity. Ecological vector fitting was conducted to project microbial taxa as vectors onto the ARG ordination space. For each taxon, the fitted vector indicates the direction of increasing abundance, while the associated squared correlation coefficient (r²; vector length) quantifies the strength of its relationship with the ordination configuration. Taxa with higher r² values were interpreted as being more strongly associated with the dominant gradients of ARG community structure, without implying that they are causal drivers of resistome variation.

## 3. Results and discussion

### 3.1 Changes in water quality parameters

All mesocosms started with the same conditions to enable cross-treatment comparisons (**Figure 1**): pH (7.81–7.84), turbidity (120–126.5 NTU), dissolved oxygen (DO; 0 mg/L, fully anoxic), and chemical oxygen demand (COD; 83–96 mg/L). These initial conditions are characteristic of severely polluted urban river water typical of sewage-receiving water bodies. Unlike most mesocosm-based studies, which incorporate a stabilization or equilibration phase prior to monitoring, mesocosms in the present study were deliberately operated without such a pre-equilibration period to capture the fast dynamics of WQPs, ARGs, and bacterial community structure. Pronounced and similar temporal shifts in WQPs were observed across all treatment conditions in the first 24–72 h, with little change towards 168 h (**Figure 1**). Among all WQPs monitored (**Supplementary Table S4**), pH, turbidity, DO, and COD changed most consistently over time, as discussed below.

**Figure 1.**
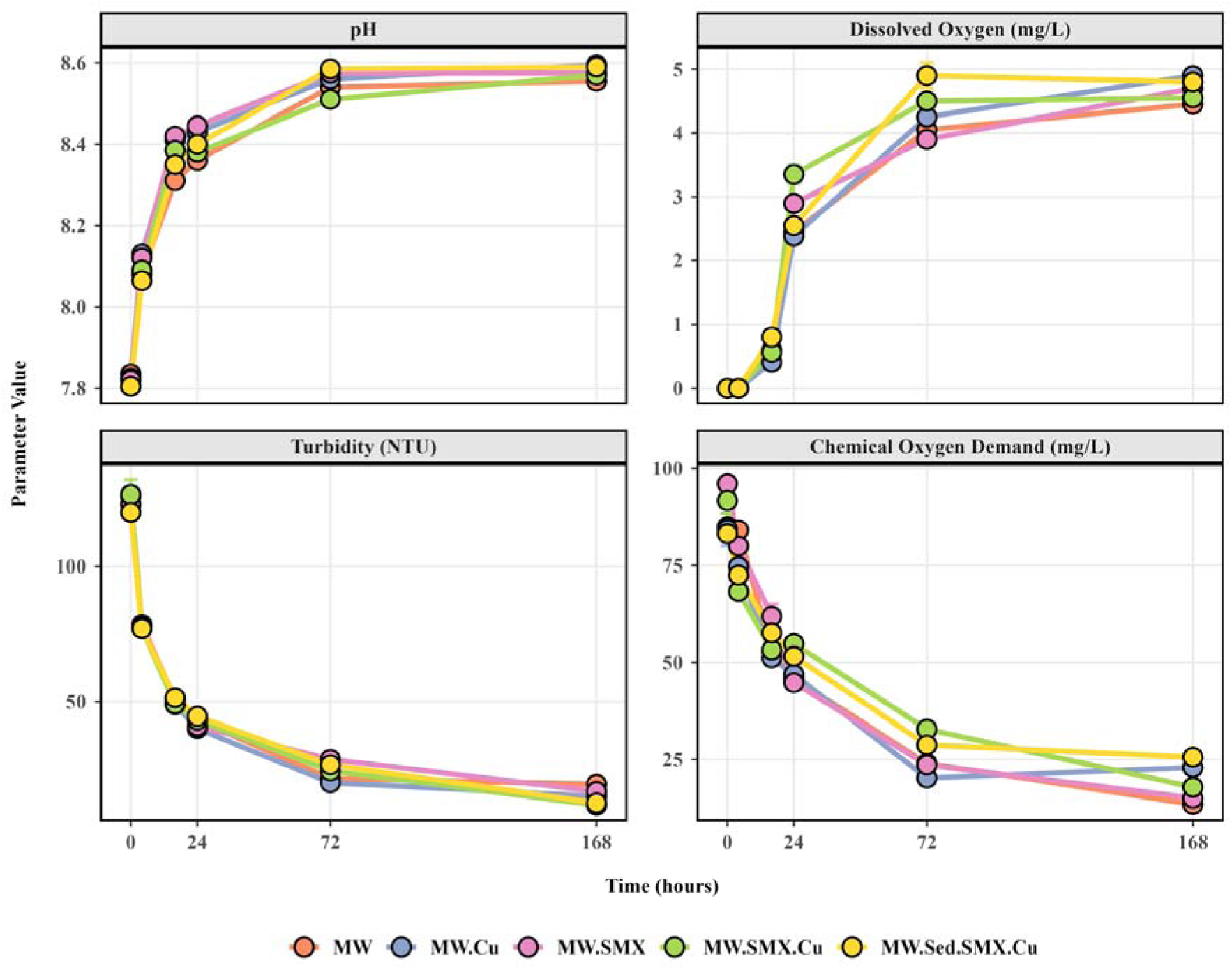
WQP dynamics for all treatment conditions (MW, MW.Cu, MW.SMX, MW.SMX.Cu and MW.Sed.SMX.Cu) over 168 h. Each data point represents the mean ± SE (n=2).

The pH increased rapidly within the first 24 h, with the rise slowing down gradually to reach a stable level of ∼8.6 across all treatments. This alkalinization was likely driven by continuous aeration (4 L/min per tank), which strips dissolved CO_2_ from the water column, thereby shifting the carbonate equilibrium toward higher pH (Kundral et al., 2015; He et al., 2017). Photosynthetic CO_2_ assimilation by indigenous algae and cyanobacteria due to the illumination (∼6,000 lumens) can further elevate pH. Comparable pH rises under aerated, light-exposed conditions have been documented in reclaimed water intake areas (He et al., 2017) and aerated pond systems (Yang et al., 2016), corroborating the mechanistic basis of our observations. The parallel development of pH values across all six treatments further indicates that this alkalinization was governed primarily by physicochemical stabilization processes (Kang et al., 2017) rather than the specific chemical stressors (SMX, Cu) applied.

Turbidity declined sharply within the first 24 h with the decline slowing down and reaching stable levels by 168 h. The rapid decrease probably reflects gravitational settling of suspended particulate matter as the mesocosms were not stirred.

DO levels remained negligible (≈ 0 mg/L) during the initial 4 h, consistent with the anoxic character of the raw sewage-polluted Musi River water used to fill the mesocosms (Sonkar et al., 2024). By 24 h of continuous aeration, DO had already reached ∼3 mg/L to stabilize at 4.4–4.9 mg/L by 168 h following the same timescales as COD decline. Thus, the applied aeration levels achieved aerobic conditions, effectively replicating the natural aeration mechanisms of flowing river systems (Jane et al., 2021; Zhang et al., 2022).

COD dropped markedly by 38–53% within the first 24 h across all treatments, driven by rapid aerobic biodegradation of dissolved organic matter (Li et al., 2008) and the even faster settling of particulate organic carbon through sedimentation (Barana et al., 2013; Peng et al., 2022). The COD reduction mirrored the rising DO profile, demonstrating a shift from anaerobic to aerobic decomposition. This is consistent with observations in aerated vertical-flow wetland systems treating polluted river water, where continuous aeration achieved the highest COD removal at a hydraulic retention time of two days (Dong et al., 2012).

Collectively, the temporal dynamics of WQPs reveal that continuous aeration caused a rapid shift of the mesocosms from initially anoxic, turbid and high-COD conditions toward aerobic, clear and low COD conditions within 24–72 h, broadly analogous to the degradation-to-recovery transition zones described in classical river self-purification models (H. W. Streeter and B. Phelps, 1925; Cox, 2003) and our observations from the Musi downstream of Hyderabad (Sonkar et al., 2024). All six treatment conditions (MW, MW.SMX, MW.Cu, MW.SMX.Cu, and MW.Sed.SMX.Cu) showed nearly identical WQP trajectories, suggesting that the mesocosm dynamics were governed by aeration-driven biodegradation of organic matter rather than the specific chemical stressors.

### 3.2 Decay kinetics of ARGs and other target genes

The decay of all target genes was modeled over the 0–72 h experimental window using first-order kinetics (**Figure 2**). The last time point (168 h) was excluded from analysis as the regression value was less (**Supplementary Table S5A and Supplementary Figure 3**). The corresponding kinetic parameters and statistics are presented in **Table 1**. To examine whether treatment conditions influenced decay rates, treatment-specific slopes were estimated for each target gene (**Supplementary Table S5B and S5C**). Given the limited statistical power at the individual treatment level (n = 5 time points per regression, df = 3), treatment-specific slope estimates carried large standard errors and showed no significant pairwise differences (**Supplementary Text 1**). Consequently, the ANCOVA model for a given gene (ln[gene i copies/ml] ∼ Time + Treatment Condition) was fitted simultaneously across all five mesocosm conditions (n = 25).

**Figure 2.**
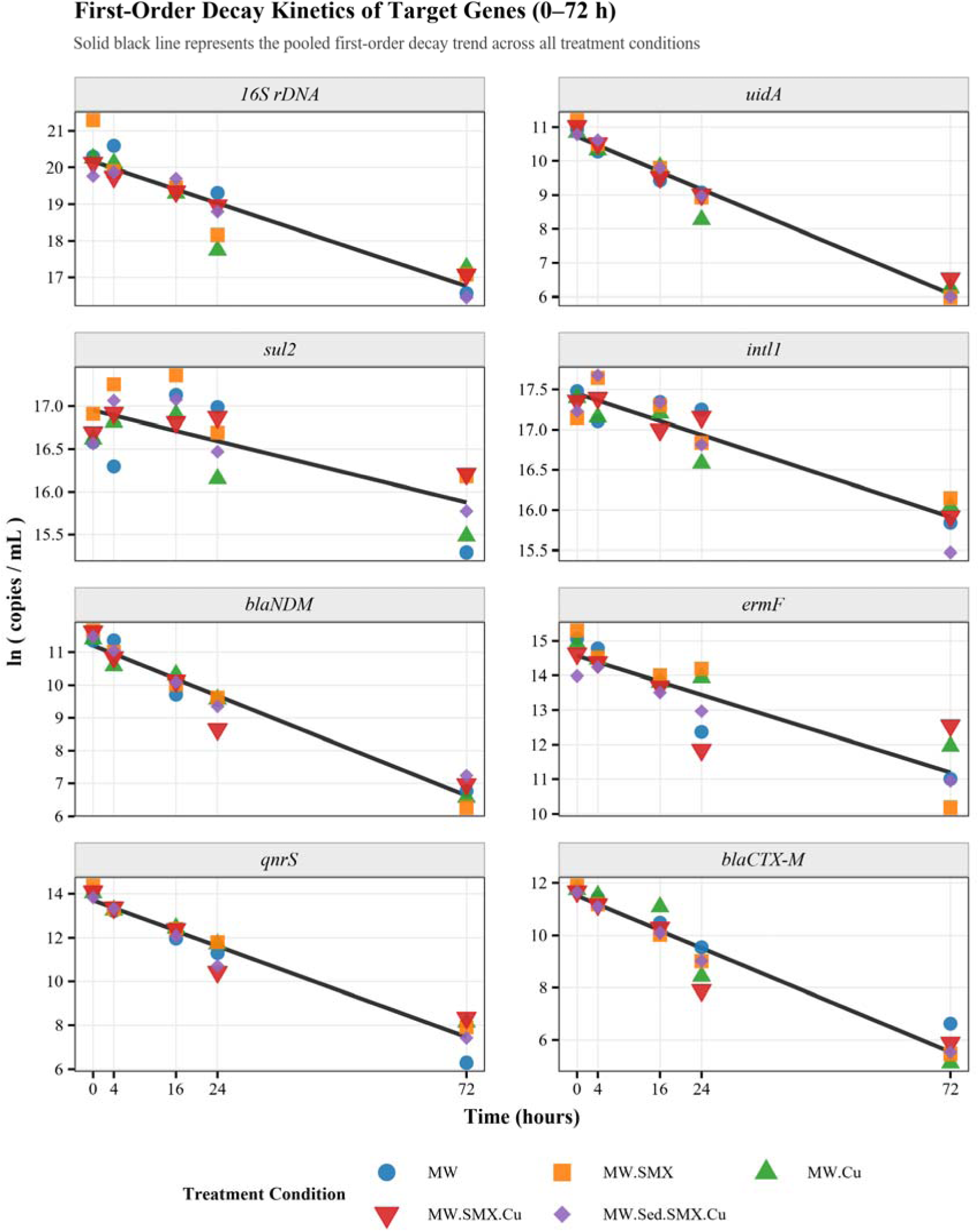
Decay kinetics of target genes (0-72 h) under different treatments using a pooled common-slope model. Points represented observed values across treatment conditions and colored lines denoted fitted first-order decay models with a shared slope per gene across treatments.

**Table 1.** Decay kinetics for each target gene (0-72 h).

| Gene | Category | $k \pm SE$ (h<sup>-1</sup>) | $t_{1/2}$ (h) | $R^2$ | Adj. $R^2$ |
| --- | --- | --- | --- | --- | --- |
| **16S rDNA** | Bacterial marker | $-0.047 \pm 0.003$ | 14.7 | 0.88 | 0.85 |
| ***uidA*** | Fecal marker | $-0.064 \pm 0.002$ | 10.8 | 0.97 | 0.97 |
| ***sul2*** | Sulfonamide | $-0.014 \pm 0.002$ | 46.5 | 0.67 | 0.58 |
| ***intl1*** | MGE | $-0.021 \pm 0.002$ | 32.5 | 0.87 | 0.84 |
| ***tetW*\*** | Tetracycline | – | – | – | – |
| ***bla*<sub>NDM</sub>** | Carbapenem | $-0.063 \pm 0.003$ | 10.9 | 0.95 | 0.94 |
| ***ermF*** | Macrolide | $-0.046 \pm 0.005$ | 14.8 | 0.81 | 0.76 |
| ***qnrS*** | Fluoroquinolone | $-0.086 \pm 0.004$ | 8.0 | 0.96 | 0.95 |
| ***aph(3'')-Ib*\*** | Aminoglycoside | – | – | – | – |
| ***bla*<sub>CTX-M</sub>** | Extended-spectrum beta-lactamase | $-0.083 \pm 0.004$ | 8.4 | 0.94 | 0.93 |

\*ANCOVA indicated a significant Time × Condition interaction for *tetW* ( $p = 0.007$ ) and *aph(3'')-Ib* ( $p = 0.009$ ), violating the common-slope assumption. Refer to the **Supplementary Table S5C**.

Bacterial Marker Genes: The *E. coli*-specific *uidA* gene declined faster (t_1/2_ = 10.8 h) than the bacterial marker 16S rDNA (t_1/2_ = 14.7 h), reflecting the compositional distinction between the sewage-associated fecal population and the total bacterial community. The drop in the 16S rDNA gene signifies the exponential decline of the overall prokaryotic community across all conditions as the mesocosms transitioned from anoxic to aerobic states with less organic carbon (Section 3.1). Similarly, Zhang et al. (2015) reported decreasing 16S rRNA gene abundance in raw-wastewater-amended batch microcosms incubated in darkness at 22–24°C, with biphasic decline in seawater and slower reduction in beach sand. Complementary freshwater experiments have also shown temporal decay of wastewater-associated bacterial 16S rDNA markers, including HF183, under dark, aerobic, continuously mixed conditions (Jeanneau et al., 2012). Although the environment differed from the present study, all observations support the occurrence of declining total bacterial 16S rRNA gene abundance during batch incubation. A faster attenuation of fecal-indicator organisms relative to the total bacterial community has been consistently observed in aquatic microcosm and mesocosm experiments and confirms that the *uidA* decline here is attributable to specific *E. coli* population loss rather than a general community-wide effect (Tsai et al., 1993; Oliver et al., 2016).

ARGs: The decay of *qnrS* was fastest (half-life 8.0 h), followed closely by *bla*_CTX-M_ (t_1/2_ = 8.4 h). The decay of *bla*_NDM_ (t_1/2_ = 10.9 h) and *tetW* (t_1/2_ = 11.5 h) was slower than *qnrS* and *bla*_CTX-M_ but faster than the *16S rDNA* while *ermF* (t_1/2_ = 14.8 h) and *aph(3’’)-Ib* (t_1/2_ = 15.4 h) decayed at a similar rate to the 16S rDNA. The only ARG decaying more slowly than the 16S rDNA was *sul2* (t_1/2_ = 46.5 h), but note the poor model fit, which may be due to an effect of the combined SMX and Cu treatment. While our data does not permit a statistically conclusive attribution to treatment-specific effects, it hints that combined presence of SMX and Cu selected *sul2*-carrying bacterial populations. This interpretation is supported by evidence from soil microcosms demonstrating that SMX and Cu co-presence significantly increases the relative abundance of *sul2*, with Cu exerting a particularly strong co-selective pressure for sulfonamide resistance (Narciso-da-Rocha et al., 2018). Co-selection of sulfonamide resistance by metal stressors has also been reported across aquatic environments, where Cu, Zn, and Hg select for *sul*-gene carrying populations through shared resistance mechanisms (Gillieatt and Coleman, 2024). In rivers, *sul2* is one of the most persistently elevated ARGs following WWTP effluent discharge into pristine rivers, detectable at significantly elevated abundances several kilometers downstream, beyond all other co-monitored ARGs in the same system. The class 1 integron-integrase gene *intI1* also decayed considerably more slowly than the 16S rDNA and ARGs other than *sul2* (t_1/2_ = 32.5 h) (Chaturvedi et al., 2021). This co-persistence of *sul2* and *intI1* could be explained by their frequent co-localization on the same mobile genetic elements, e.g., *sul*-gene cassettes are commonly embedded with class 1 integrons (de los Santos et al., 2021).

### 3.3 Ordination shows temporal shifts in ARG composition

To determine whether the seven ARG profiles underwent systematic compositional shifts over the 168 h experiment, ordination analyses were performed on log-transformed absolute gene abundance data. The first two principal components together captured 93.1% of the total ARG compositional variance (PC1 = 86.3%; PC2 = 6.8%), providing an informative low-dimensional representation of resistome dynamics across all treatments and time points (**Figure 3**).

**Figure 3.**
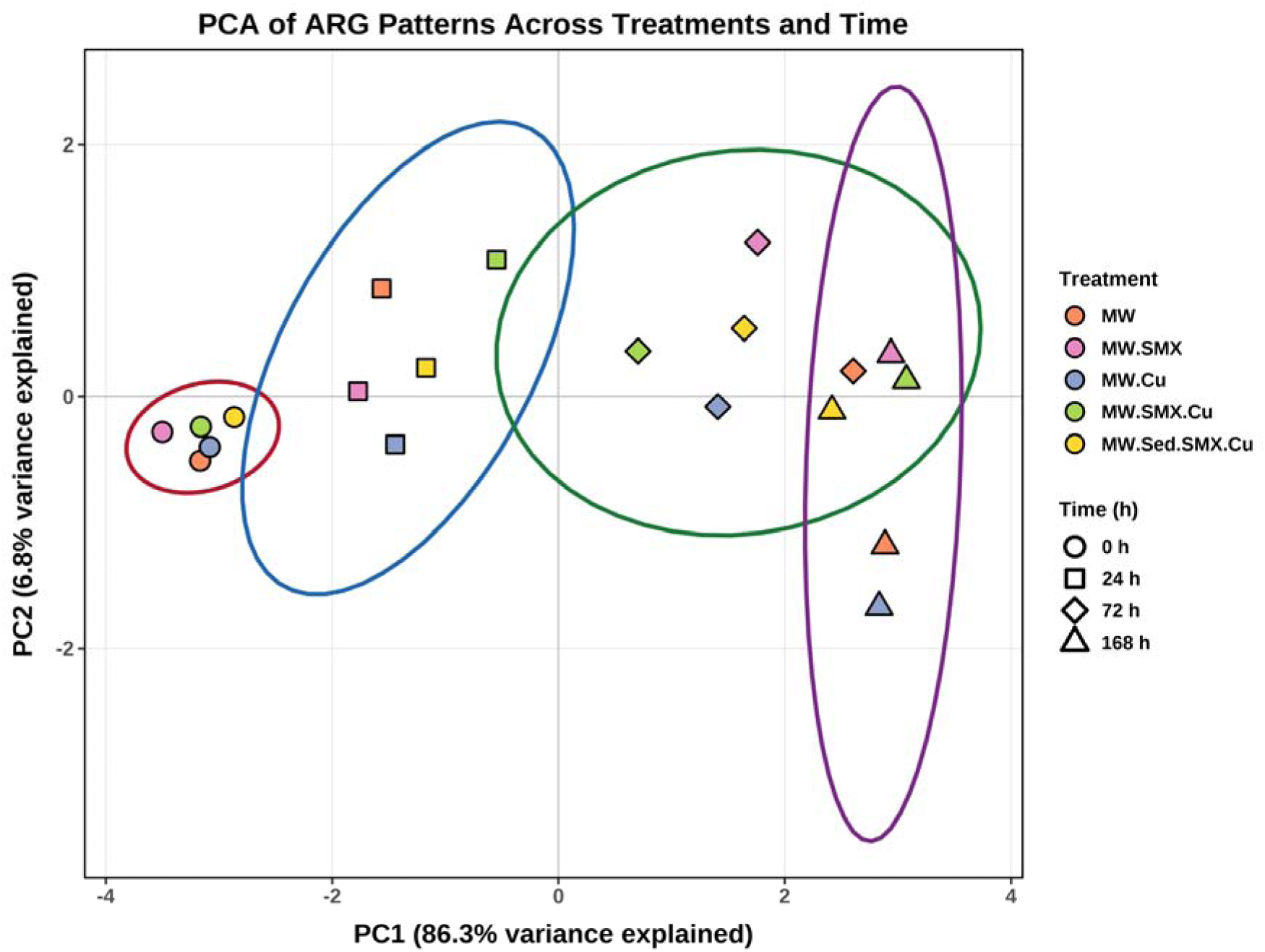
PCA of ARG profiles across treatments (color-coded) and time (symbol-coded). Ellipses show clustering by time. The first two principal components explained 93.1% of the total variance.

All seven resistance genes loaded strongly and uniformly in the same negative direction along PC1, with loadings ranging from −0.321 (*sul2*) to −0.401 (*qnrS*), indicating that the decline of the entire resistome rather than gene-specific responses was the main feature (**Supplementary Table S6**). Samples clustered tightly at the beginning and shifted consistently along PC1 over time, while dispersing along PC2. This indicates that all five mesocosm treatments started from statistically similar ARG compositions (as intended) and reached comparable final states, with more dispersion at intermediate time points. Dispersion is likely due to treatments, but as we lack replication, we cannot draw conclusions.

### 3.4 Bacterial community compositional shifts

The composition of prokaryotic communities underwent drastic changes over the 168 h treatment period (relative abundances in **Figure 4** and **Supplementary Figure S6**, absolute abundances in **Supplementary Figure S7**). At the start, all mesocosms were dominated by *Campylobacterota* (90–93%), followed by *Pseudomonadota* (*Proteobacteria)* (5.3–6.4%), *Bacillota* (*Firmicutes)* (1.4–2.7%), *Halobacteriota* (0.3–4.3%), and other phyla (< 1%). This strong *Campylobacterota* dominance is consistent with wastewater-impacted rivers (Fisher et al., 2014; Cui et al., 2019). The prevalence of the *Campylobacterota* genus *Arcobacter* likely reflects the low and fluctuating dissolved oxygen levels of polluted stretches of the Musi River, where high organic inputs are coupled with intermittent oxygenation *via* mixing and photosynthesis, that favor the aerotolerant *Arcobacter*.

**Figure 2.**
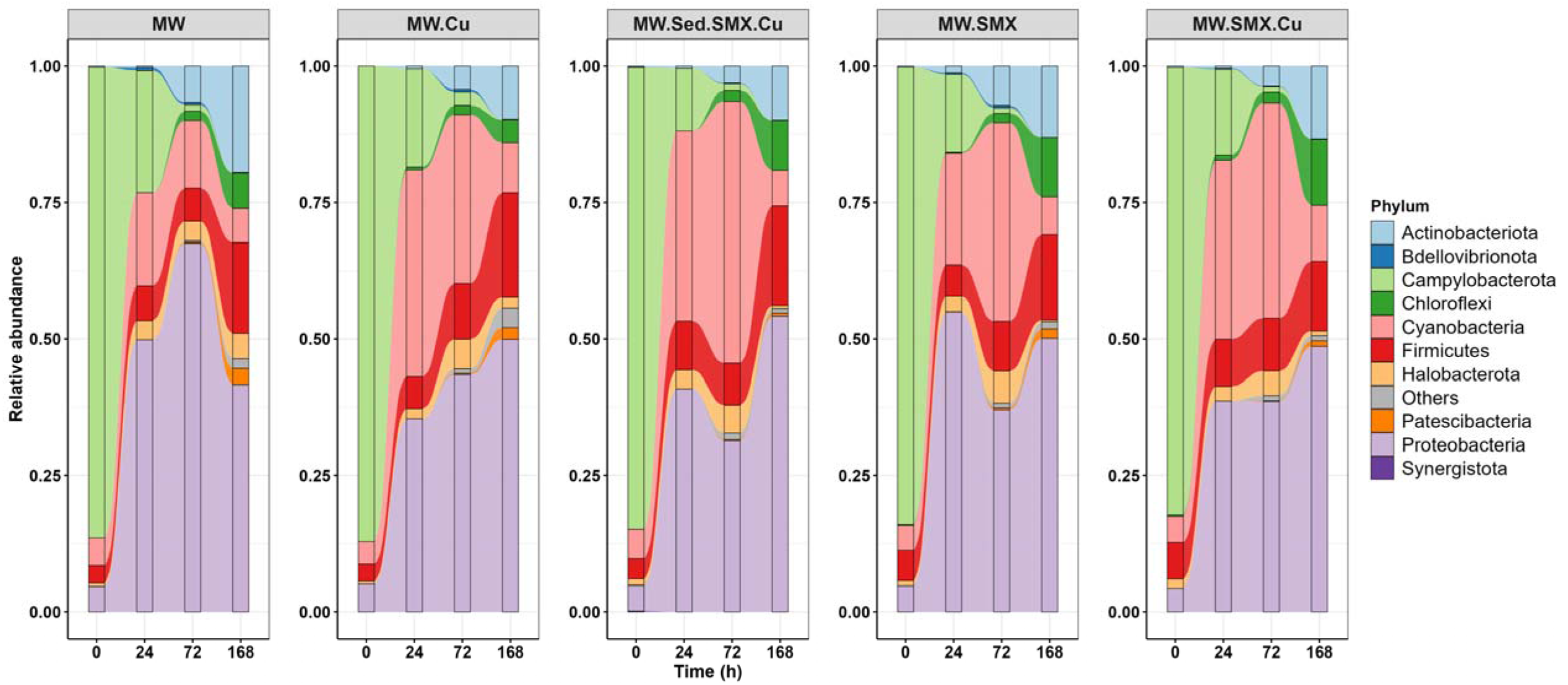
Changes in relative abundance (%) of the major prokaryotic phyla in mesocosm reactors across 0, 24, 72, and 168 h based on 16S rDNA amplicon sequencing. “Others” indicates all low-abundance phyla. For the family and genus levels, see Supplementary Figure S6 and for absolute abundance **see Supplementary Figure S7.**

By 24 h, *Campylobacterota* had almost disappeared across all treatments, the drop was especially drastic in absolute abundance (**Supplementary Figure S7**). There was a concurrent drop of *Pseudomonadota* in absolute abundance but rise to dominance in relative abundance. *Pseudomonadota* relative abundance declined somewhat in the unamended MW mesocosm. The relative abundance of several other phyla also increased substantially, such as *Bacillota*, *Actinomycetota* (*Actinobacteria)*, *Cyanobacteriota* (Cyanobacteria) and *Chloroflexota* (*Chloroflexi)* but note that the absolute abundances of these taxa declined markedly.

Overall, the transition from high density *Campylobacterota*-dominated communities toward low density *Pseudomonadota*, *Bacillota*, and *Actinomycetota* -enriched assemblages is notable, as these latter phyla are frequently associated with higher ARG carriage, mobile genetic elements, and stress-adaptive traits in wastewater-impacted environments (Leroy-Freitas et al., 2022; K. Liu et al., 2024; Li et al., 2024). The direction of community change was consistent across treatments, indicating that sulfamethoxazole, copper or sediment amendments did not fundamentally alter short-term successional pathways, but may have modulated the relative abundance and persistence of specific bacterial groups. Collectively, these results indicate that short-term microbial community restructuring in sewage-polluted river water is driven primarily by oxygenation and decreasing organic matter.

### 3.5 Quantitative microbiome profiling using Hill diversity

*Alpha-Diversity*: Using quantitative microbiome profiling (QMP) and a range of Hill numbers, the overall trend was a drastic decrease in diversity over time, with some transient increases under certain conditions (**Figure 5(A)**). Initial richness ranged from 888 (MW.Cu) to 1,418 ASVs (MW.SMX) and collapsed to between 36 (MW.Cu) and 66 (MW.SMX) ASVs by 168 h, representing a ∼95% decline in absolute taxon richness across all conditions (**Supplementary Figure S8 and Supplementary Table S7**). The exponential of Shannon diversity (q = 1) followed a similar trend to richness but with a tendency to increase (or decrease less) transiently. Initial values ranged from 494 (MW.Cu) to 673 (MW.SMX), reflecting communities with hundreds of effectively contributing taxa. By 168 h, values had collapsed to between 35 (MW.Cu) and 64 (MW.SMX), confirming that both rare and dominant taxa were lost in absolute terms. The inverse Simpson (q = 2, sensitive only to dominant taxa) showed a similar decline from 297.6–381.9 at the start to 33.0–62.2 at the end across conditions, indicating that even the dominant core community underwent substantial absolute reduction. Transient increases in diversity measures were found in Cu and SMX containing treatments, particularly the combined treatment and more pronounced in the higher order Hill numbers (larger q). This transient increase in diversity under Cu and/or SMX treatment likely reflects a temporary rise in various taxa that were expanding due to reduced competition caused by the reduction of dominant taxa, before substrate depletion set in.

**Figure 5.**
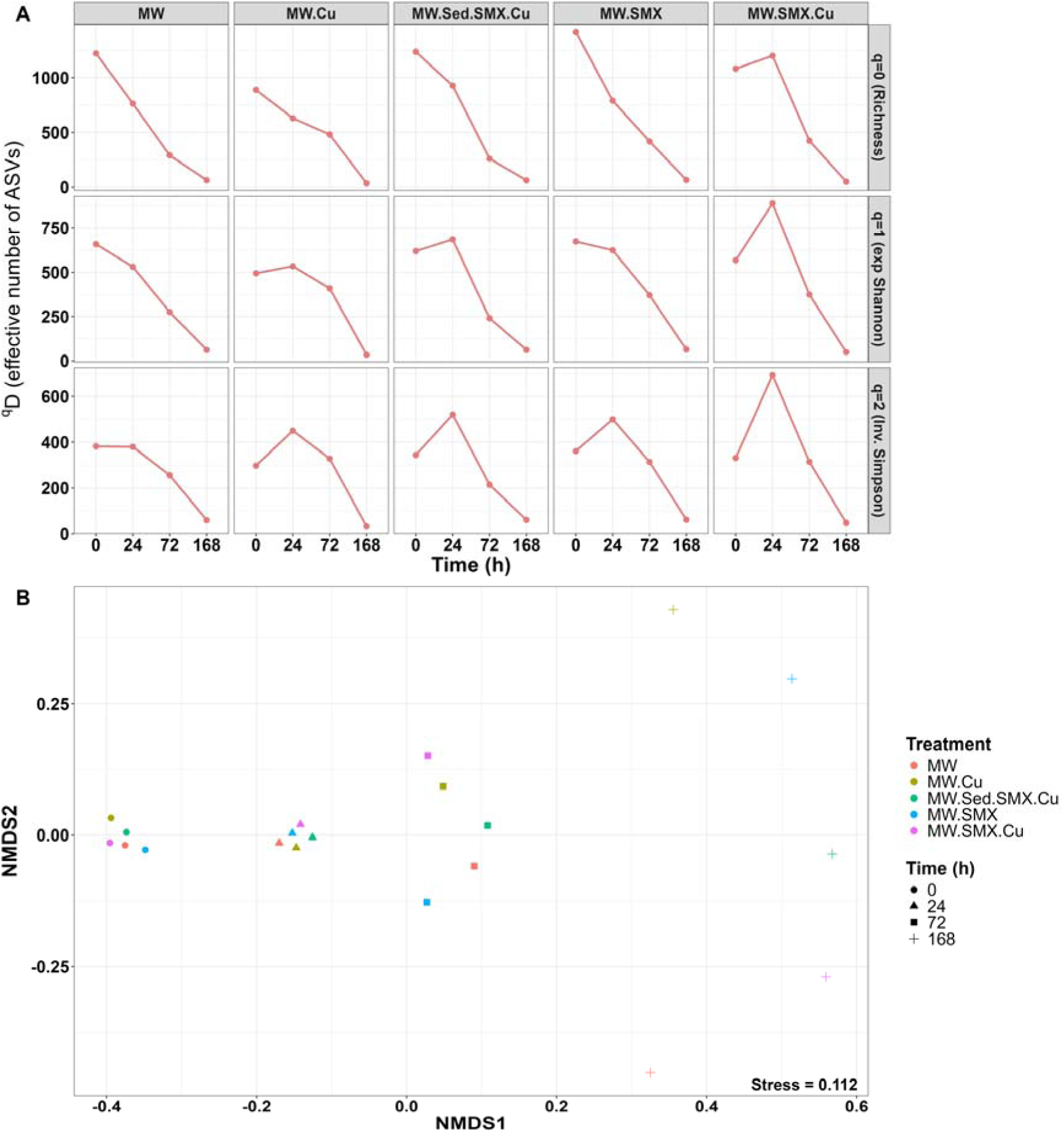
**(A)** Temporal changes in quantitative alpha diversity (Hill numbers: q = 0 (Richness), q = 1 (Exponential of Shannon index), and q = 2 (Inverse Simpson index)) across mesocosm treatments showing a strong decline in effective bacterial diversity over the 168 h incubation period, with transient increases in diversity under copper-containing conditions at 24 h. **Supplementary Figure S8** shows the trends of the Hill numbers ^q^D with increasing q. **(B)** NMDS ordination (stress = 0.111) of Sørensen-type beta diversity (q = 1) based on QMP absolute abundance data showing strong restructuring of bacterial communities over time, with increasing community divergence across mesocosms during the 168 h incubation period.

#### Beta Diversity

Sørensen-type beta diversity at q = 1 was ordinated *via* NMDS (**Figure 5(B)**) using QMP absolute abundance tables following the approach of Ott et al. (2021). The NMDS ordination revealed a clear trend over time with increasing divergence between mesocosms/treatments (although we cannot exclude that this divergence is by chance). As expected, mesocosms clustered tightly at the beginning, followed by a temporal shift primarily along NDMS1 while the divergence between mesocosms was both along NDMS1 and NDMS2, indicating progressive community restructuring over time that substantially exceeded any within-timepoint treatment-level separation. These temporal shifts in community composition were similar to the shifts of the resistome (**Figure 3**) with divergence more pronounced.

### 3.6 PICRUSt2 analysis

PICRUSt2-based functional prediction indicated distinct temporal shifts in putative microbial metabolic guilds across the mesocosm treatments (**Figure 6**). Overall, the functional profiles suggested progressive restructuring from anaerobic to aerobic and oxygenic phototrophic communities during incubation, broadly consistent with the observed physicochemical shifts and bacterial succession patterns. The aerobic functional marker (*cytochrome c oxidase*) increased over time in all mesocosms indicating increasing aerobic metabolic potential during progressive oxygenation of the mesocosms. Surprisingly, the functional marker for fermentative metabolism (*alcohol dehydrogenase*) also increased in all mesocosms, maybe because the same enzyme is used to oxidize ethanol by aerobic microorganisms. Predicted nitrifier-associated markers (*ammonia monooxygenase*) generally increased, either steadily or peaking at intermediate stages in some mesosms, particularly in MW, whereas denitrifier-associated functions (*nitrite reductase*) were most abundant initially and then gradually declining. These shifts are consistent with the transition from initially reduced conditions (denitrification as anaerobic respiration) toward increasingly oxygenated environments (nitrification as aerobic respiration). Chlorophyll synthesis as a functional marker for photosynthesis inceased in all mesocosms either gradually or peaking at intermediate stages as expected from the increasing availability of oxygen and the illumination of the mesocosms. Intriguingly, the Calvin cycle as a supposed marker for photosynthesis showed opposing trends. The contrast was strongest between negligible chlorophyll synthesis and highly abundant Calvin cycle in the sediment fraction of the sediment amended mesocosm. This may be due to the Calvin cycle being used in some anoxygenic phototrophs (purple bacteria using bacteriochlorophyll) and chemolithoautotrophs (nitrifiers, sulfide, iron or hydrogen oxidizers) and not only in oxygenic phototrophs. The methanogenesis-associated marker (*Methyl-CoM reductase*) followed the same pattern as the Calvin cycle, remaining low across all treatments but high in the sediment fraction of the sediment-containing mesocosm (**Supplementary Figure S18**).

**Figure 6.**
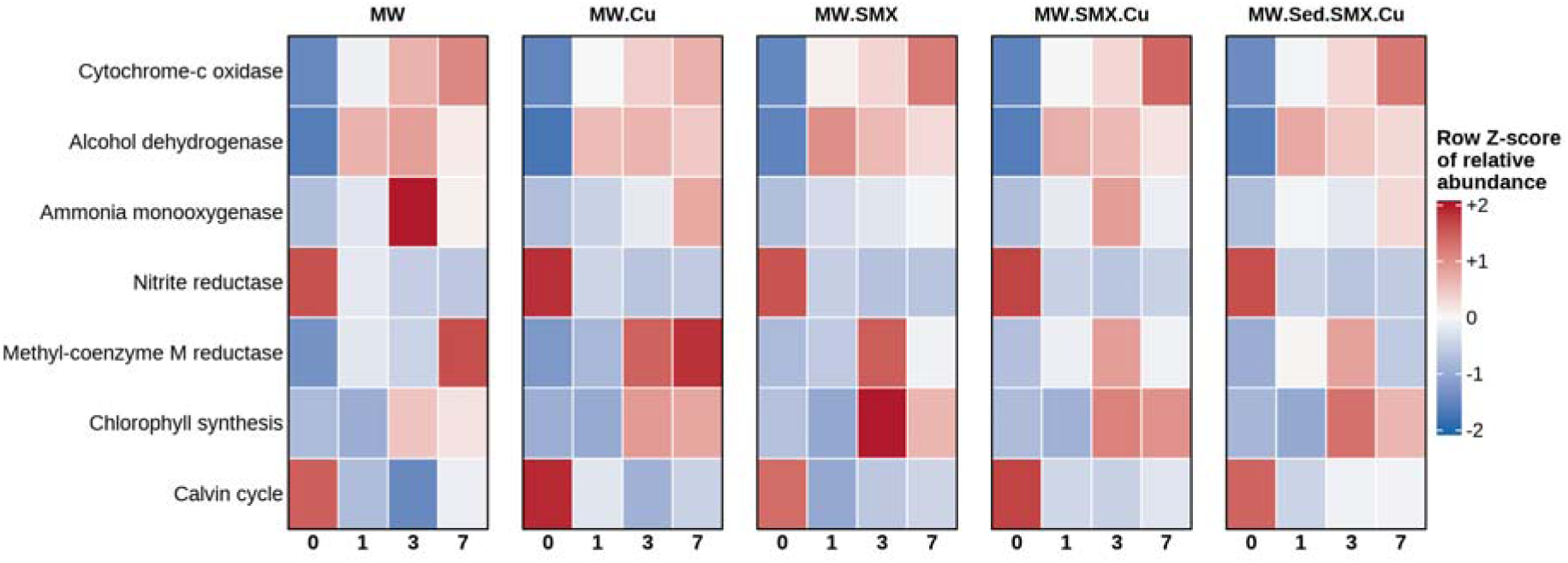
Temporal changes in PICRUSt2-predicted microbial functional groups across mesocosm treatments during the 168 h experiment.

### 3.7 Constrained ordination (db-RDA) of resistome and microbial community changes

To evaluate the relative influence of temporal progression, treatment exposure, and physicochemical changes on antimicrobial resistance and microbial community dynamics, db-RDA coupled with variation partitioning analysis was performed on ARG and bacterial community composition datasets. These multivariate analyses were used to disentangle the independent and shared contributions of time, treatment, and water quality gradients to resistome and community restructuring during mesocosm evolution.

#### Temporal restructuring governed both resistome and bacterial community composition

db-RDA demonstrated that temporal progression was the primary determinant of changes in both ARG composition and bacterial community structure throughout the mesocosm incubation period. In contrast, effects of SMX, Cu, or their combined treatment were negligible.

For ARG composition, the combined model including time and treatment explained a substantial proportion of the total variation (R^2^ = 0.668; adjusted R^2^ = 0.549, **Figure 7(A)** and **Supplementary Table S8**). This was due to the time-only model that yielded comparable explanatory power (Adj.R^2^ = 0.576), as the treatment-only model did not explain the variation at all (Adj. R^2^ = -0.179). Permutation analysis confirmed that time significantly structured ARG composition (F = 25.24, p = 0.001), while treatment effects were not significant (F = 0.73, p = 0.640). Partial db-RDA further showed that treatment remained non-significant after conditioning on time (F = 0.73, p = 0.354), whereas time retained strong explanatory power independent of treatment (Adj.R^2^ = 0.728).

**Figure 7.**
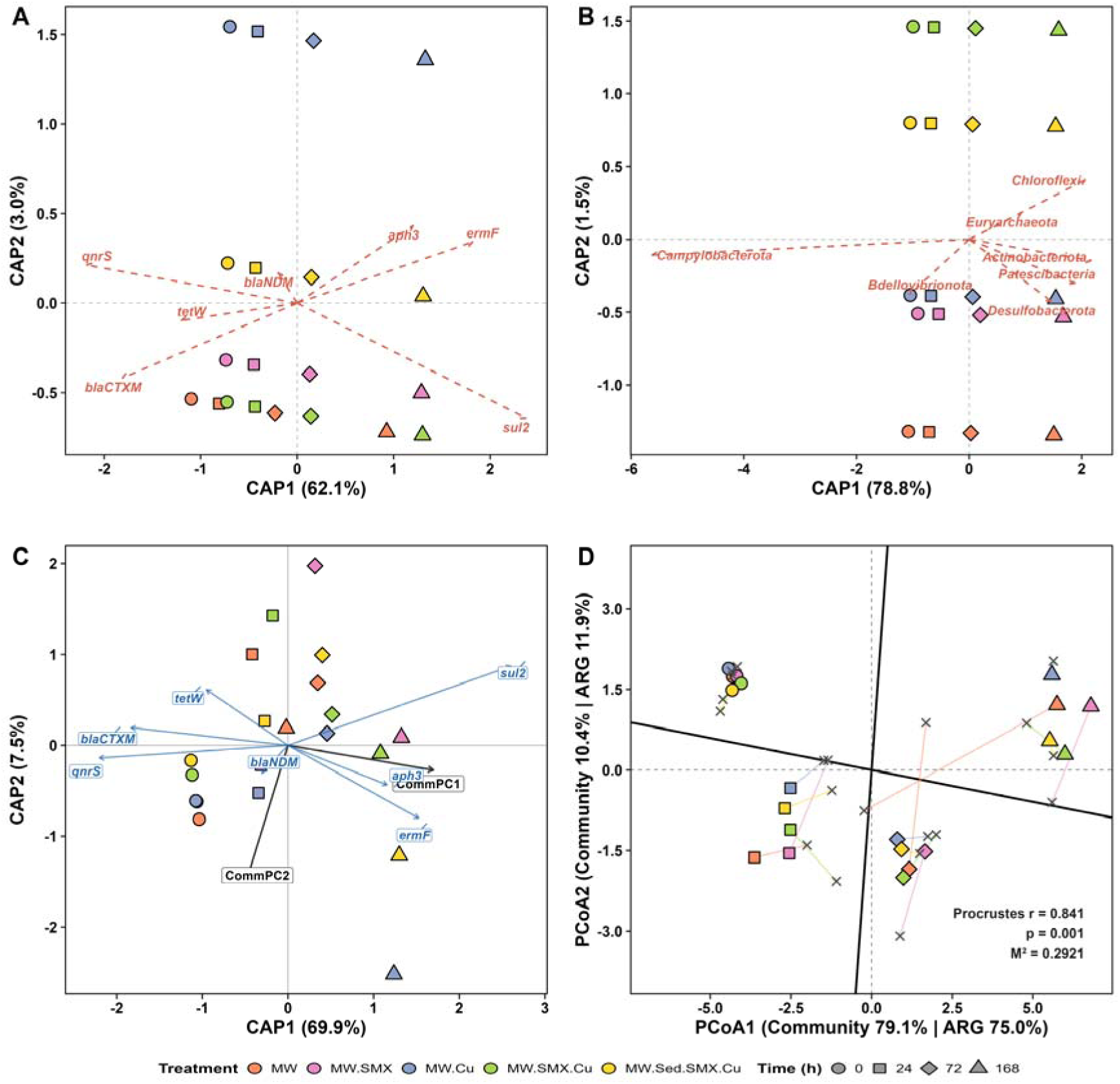
db-RDA ordination of **(A)** ARG composition showing strong temporal structuring of resistome profiles across mesocosm treatments, with time explaining all of the variation and treatment none. **(B)** Phylum-level bacterial community composition showing strong temporal succession across mesocosm treatments, with time explaining substantially greater community variation than treatment effects. **(C)** showing the association between phylum-level bacterial community and ARG composition across mesocosm treatments. **(D)** Procrustes analysis showing strong concordance (M^2^ = 0.292, p = 0.001) between phylum-level bacterial community composition and ARG ordination patterns across mesocosm treatments.

A similar pattern was observed for phylum-level bacterial community composition (**Figure 7(B)**, and **Supplementary Table S8**). The full model explained a large proportion of community variation (R^2^ = 0.814; adjusted R² = 0.748), with time as the dominant factor (F = 59.17, p = 0.001). In contrast, treatment effects were not significant (F = 0.55, p = 0.764) and the treatment-only model had no explanatory power (Adj. R^2^ = -0.230). After accounting for temporal effects, treatment remained non-significant (F = 0.55, p = 0.07; Adj. R^2^ = -0.025).

Phylum-level scores further indicated that the temporal restructuring was driven primarily by dominant taxa (**Supplementary Table S9**). *Campylobacterota* exhibited the strongest loading along CAP1 (−3.13), whereas *Pseudomonadota* showed moderate positive association with the constrained axis (0.22). *Bacillota*, *Cyanobacteriota*, and *Halobacteriota* contributed little to constrained variation. These shifts are consistent with community restructuring (**Supplementary Figure S9 and Supplementary Figure S10**) occurring under increasingly oxygenated conditions.

#### Physicochemical changes were coupled with resistome and bacterial community dynamics

The influence of WQPs on ARG and bacterial community composition was evaluated using db-RDA coupled with variation partitioning against temporal effects. Both resistome and phylum-level community composition was clearly associated with WQP changes; however, a substantial proportion of this relationship was shared with temporal progression, suggesting that the changes in WQP over time (arising from aeration and irradiation interacting with microbial activities) were the underlying reason for temporal restructuring (**Supplementary Text 2**).

Phylum score analysis (**Supplementary Table S11**) indicated that major phyla responded differentially to the dominant environmental changes. *Campylobacterota* exhibited strong negative loading with CAP1 (−3.05), whereas *Proteobacteria* displayed positive loading (0.25). These opposing loadings are consistent with restructuring under increasing oxygenation and decreasing organic matter during incubation.

#### Coupling between bacterial community succession and resistome restructuring

To evaluate whether bacterial community succession contributed to resistome restructuring during mesocosm incubation, coupling between microbial community and ARG composition was assessed using db-RDA, Procrustes superimposition analysis and Mantel correlation. These multivariate approaches consistently demonstrated strong concordance between microbial community succession and resistome restructuring throughout the experimental period. The db-RDA model incorporating community composition, time, and treatment explained a substantial proportion of ARG variation at the phylum level (R^2^ = 0.813; Adj. R^2^ = 0.704; p = 0.002, **Supplementary Table S12**). Community composition retained measurable explanatory contribution beyond temporal and treatment effects (R^2^ = 0.145), indicating that bacterial succession accounted for additional resistome variation independent of mesocosm treatments.

At phylum level, permutation analysis (**Figure 7(C)**, **Supplementary Table S13**) identified the primary community ordination axis (CommPC1) as the major factor associated with ARG composition (F = 40.15, p = 0.001), while CommPC2 also showed a significant but smaller contribution (F = 6.79, p = 0.028). After incorporation of community structure into the constrained model, the independent effect of time lost significance (F = 1.18, p = 0.202), suggesting that much of the temporal variation in ARG composition was due to the microbial community restructuring itself. Although treatment remained statistically significant (F = 1.01, p = 0.002), its contribution was weaker.

Ecological vector fitting identified several phyla strongly associated with the dominant resistome restructuring gradient (**Supplementary Table S14** and **Supplementary Figure S13**). *Campylobacterota* showed the strongest negative association along CAP1 (loading = −0.944; R^2^ = 0.891; adjusted p = 0.0028), whereas *Actinobacteriota* (loading = 0.874; R^2^ = 0.763; adjusted p = 0.0028), *Chloroflexota* (loading = 0.839; R^2^ = 0.707; adjusted p = 0.0028), and *Euryarchaeota* (loading = 0.728; R^2^ = 0.594; adjusted p = 0.0070) showed strong positive associations. These contrasting ordination loadings suggest that shifts in ARG composition were driven by changes in bacterial community structure during mesocosm stabilization.

The strong association of *Campylobacterota* with one end of the constrained ordination space agrees with the earlier community analysis (Section 3.3) showing this phylum was associated with the initial sewage-derived conditions characterized by low dissolved oxygen and high organic loading. As the mesocosm environment changed to aerobic and lower in carbon, the other phyla became more closely associated with later-stage resistome patterns. The link between bacterial succession and resistome restructuring was further supported by Procrustes analysis of Aitchison distance matrices (**Figure 7(D)**). Significant positive correlations were observed across all taxonomic levels, indicating that samples with higher microbial community dissimilarity also showed higher resistome dissimilarity.

Notably, db-RDA, Procrustes and Mantel analyses were performed using time-blocked permutation schemes to reduce the influence of temporal autocorrelation on statistical significance. Therefore, the observed relationships are unlikely to be solely driven by incubation time and instead support a consistent association between bacterial community succession and resistome restructuring. Overall, these findings suggest that physicochemical changes, microbial succession, and ARG restructuring were interlinked during mesocosm incubation. Such linkage between community shifts and ARG composition and abundance is common but often overlooked when interpreting changes in ARGs naively (Uluseker et al., 2025).

## 4. Study limitations

The main limitation of the current study is the unreplicated experimental design, which can be used to explore potential effects of the treatments but not to estimate effect sizes and test hypotheses. Nevertheless, our results suggest that treatment effects are small relative to temporal changes. Due to the unreplicated design, we focused on the changes within each mesocosm. The mesocosms were meant to mimic river conditions, including aeration and irradiation, but more realistic simulation of these conditions would have required measuring river conditions and controlling these experimentally. Rivers also continuously receive pollutant inflows, which the mesocosms did not on purpose, because this batch operation enabled the measurement of decay kinetics.

## 5. Conclusions

We filled aerated and illuminated mesocosms with wastewater polluted river water from the Musi in Hyderabad, India, to measure temporal dynamics of physicochemical parameters, microbial community composition via 16S amplicon sequencing and ARGs and taxonomic markers via qPCR. Mesocosms became aerobic while COD halved within a day. Most genes decayed rapidly, approximately following first order kinetics with half-lives from 8 to 15 hours, demonstrating that rivers are ‘self-cleaning’. However, self-cleaning likely takes longer in colder climates. Concomitantly, the absolute abundance of microbes as well as diversity dropped markedly while the community composition shifted from sewage-associated anaerobic *Campylobacterota* to aerobic *Pseudomonadota* dominance. db-RDA, Procrustes and Mantel analysis showed that the changes in the community composition were coupled with the changes in the resistome. Surprisingly, we found no effects of subinhibitory concentrations of SMX or Cu, single or combined. Overall, this study provides approximate kinetic parameters under environmentally relevant conditions for modelling the fate of microbial groups and ARGs entering aquatic water bodies with wastewater point sources.

## CRediT authorship contribution statement

**Arun Kashyap:** Conceptualization, Methodology, Formal analysis, Validation, Visualization, Investigation, Data Curation, Writing -Original Draft, Writing - Review & Editing. **Vikas Sonkar:** Methodology, Data Curation, Validation, Investigation, Writing – review & editing. **David W. Graham:** Conceptualization, Formal analysis, Funding acquisition, Writing – original draft, review & editing. **Jan-Ulrich Kreft:** Conceptualization, Methodology, Formal Analysis, Supervision, Funding acquisition, Validation, Project administration, Writing – original draft, review & editing. **Shashidhar Thatikonda:** Conceptualization, Supervision, Funding acquisition, Project administration, Methodology, Validation, Investigation, Writing – review & editing.

## Supporting information for publication

All data, including the additional figures and necessary tables are made available as part of supplementary information.

## Acknowledgements

We cheerfully acknowledge all laboratory associates for their invaluable assistance in this work. We sincerely thank Dr. Sangeetha C. Jambu for reviewing this manuscript and assisting with manuscript refinement.

## References

1. Gajjar, K., Patel, S., Chaudhary, M., Agrawal, D., Maniyar, R., Chaudhary, D., Patel, C.K., Joshi, C., Joshi, M., Dharajiya, D., 2026. Metagenomic insights reveal the impact of natural farming on soil nutrients, enzyme activities, microbial communities, and yield in turmeric cultivation. BMC Plant Biol. 26, 28. 10.1186/s12870-025-07781-3

2. Sonkar, V., Kashyap, A., Pallares-Vega, R., Sasidharan, S.S., Modi, A., Uluseker, C., Jambu, S.C., Mohapatra, P.K., Larsen, J., Graham, D.W., Thatikonda, S., Kreft, J.-U., consortium, Amr., 2026. Hydraulic modelling reveals untreated sewage, not pharmaceutical waste, drives antimicrobial resistance in a small river running through a big city. bioRxiv 2024.12.21.629897. 10.1101/2024.12.21.629897

3. Jia, J., Liu, Q., Wang, T., Zou, B., Xiong, X., Xu, J., Wu, C., 2025. Multiple stressors enhance Microcystis dominance and modulate phycospheric antibiotic resistome in aquatic mesocosm. J. Hazard. Mater. 497, 139633. 10.1016/j.jhazmat.2025.139633

4. Liu, N., Zhang, L., Xue, H., Yang, Z., Meng, F., 2025. Sources, dissemination, and risk assessment of antibiotic resistance in surface waters: A review. Emerg. Contam. 11, 100455. 10.1016/j.emcon.2024.100455

5. Sonkar, V., Kashyap, A., Pallares-Vega, R., Sasidharan, S.S., Chandrakalabai Jambu, S., Naorem, N., Graham, D.W., Kreft, J.U., Thatikonda, S., 2025. Removal of antimicrobial resistance determinants from wastewater: Role of capacity overloading and treatment technology in wastewater treatment plants. J. Environ. Manage. 395, 127897. 10.1016/j.jenvman.2025.127897

6. Tiwari, D., Kumar, R., Yadav, M., Gupta, G.K., Singh, S. kumar, Dhapekar, N.K., Alotaibi, M.A., Sharma, R., 2025. Holistic analysis of Ganga basin water quality: a statistical approach with WQI, HMCI, HMQI and HRI indices. RSC Adv. 15, 3290–3316. 10.1039/d4ra06144f

7. Uluseker, C., Raguideau, S., Quince, C., Kreft, J.-U., 2025. Inferring antibiotic resistance selection in the environment can be confounded by correlations between resistance genes and unrelated functional traits. bioRxiv 2025.10.12.681873. 10.1101/2025.10.12.681873

8. Yan, B., Huang, F., Ying, J., Zhou, D., Norouzi, S., Zhang, X., Wang, B., Liu, F., 2025. Global antibiotic hotspots and risks: A One Health assessment. Environmental Science and Ecotechnology 25, 100564. 10.1016/j.ese.2025.100564

9. Yu, Z., Gray, D.A., Fick, J., Waters, N., Lindberg, R., Grabic, R., Tysklind, M., Ekwanzala, M.D., Martiny, H.M., Flach, C.F., Aarestrup, F.M., Larsson, D.G.J., 2025. Antibiotic resistance selection and deselection in municipal wastewater from 47 countries. Nature Communications 2025 16:1 16, 9698-. 10.1038/s41467-025-65670-7

10. Zhu, C., Wu, Linwei, Ning, D., Tian, R., Gao, S., Zhang, B., Zhao, J., Zhang, Ya, Xiao, N., Wang, Y., Brown, M.R., Tu, Q., Zhuang, W., Zhou, H., Zheng, W., Zhang, W., Zhang, Q., Zhang, C., Young, M., Yang, M., Yan, T., Xi, C., Wu, Liyou, Wu, J.H., Woo, S.G., West, S., Weaver, J.E., Wang, B., Wakelin, S., Van Nostrand, J.D., Tooker, N.B., Sun, C., Stephens, K., de Sousa, O.V., Smith, K., Sidhu, J., Rossetti, S., de los Reyes, F.L., Reginatto, V., Parameswaran, P., Palmer, A., Meyers, A.J., de Menezes, F.G.R., Mendonça-Hagler, L.C., Liu, Y., Li, M., Li, Z., Li, Y., Lee, Z.M.P., Leal, C.D., Kumari, S., Keucken, A., Marcantini, D.G., Johnson, D.R., Hu, Z., Horn, H., Harmon, M., Hale, L., Habagil, M., Gu, A.Z., Griffin, J.S., Gómez, J.S., Ford, A., Etchebehere, C., Chen, S., Cabrol, L., Cabezas, A., Bux, F., Brewster, R.K., Bovio-Winkler, P., Bott, C.B., Bond, P., Boehnke, K., de Abreu Mac Conell, É.F., de Araujo, J.C., Agullo-Barcelo, M., Acevedo, D., Ju, F., Wells, G.F., Guo, J., He, Z., Nielsen, P.H., Wang, A., Zhang, Yu, Chen, T., He, Q., Criddle, C.S., Wagner, M., Tiedje, J.M., Curtis, T.P., Wen, X., Yang, Y., Alvarez-Cohen, L., Stahl, D.A., Alvarez, P.J.J., Rittmann, B.E., Zhou, J., 2025. Global diversity and distribution of antibiotic resistance genes in human wastewater treatment systems. Nature Communications 2025 16:1 16, 4006-. 10.1038/s41467-025-59019-3

11. Dueholm, M.K.D., Andersen, K.S., Korntved, A.K.C., Rudkjøbing, V., Alves, M., Bajón-Fernández, Y., Batstone, D., Butler, C., Cruz, M.C., Davidsson, Å., Erijman, L., Holliger, C., Koch, K., Kreuzinger, N., Lee, C., Lyberatos, G., Mutnuri, S., O’Flaherty, V., Oleskowicz-Popiel, P., Pokorna, D., Rajal, V., Recktenwald, M., Rodríguez, J., Saikaly, P.E., Tooker, N., Vierheilig, J., De Vrieze, J., Wurzbacher, C., Nielsen, P.H., 2024. MiDAS 5: Global diversity of bacteria and archaea in anaerobic digesters. Nat. Commun. 15, 5361. 10.1038/s41467-024-49641-y

12. Gillieatt, B.F., Coleman, N. V, 2024. Unravelling the mechanisms of antibiotic and heavy metal resistance co-selection in environmental bacteria. FEMS Microbiol. Rev. 48, fuae017. 10.1093/femsre/fuae017

13. Jampani, M., Mateo-Sagasta, J., Chandrasekar, A., Fatta-Kassinos, D., Graham, D.W., Gothwal, R., Moodley, A., Chadag, V.M., Wiberg, D., Langan, S., 2024. Fate and transport modelling for evaluating antibiotic resistance in aquatic environments: Current knowledge and research priorities. J. Hazard. Mater. 461, 132527. 10.1016/J.JHAZMAT.2023.132527

14. Li, K., Zhu, Y., Shi, X., Yan, M., Li, J., Zhang, W., Shao, Yingying, Shao, Yanqiu, 2024. Effects of Zn and oxytetracycline on mobile genetic elements, antibiotic resistance genes, and microbial community evolution in soil. Environmental Pollution 341, 122609. 10.1016/j.envpol.2023.122609

15. Liu, K., Li, Y., Ge, Z., Huang, D., Zhang, J., 2024. Microbial communities and mobile genetic elements determine the variations of antibiotic resistance genes for a continuous year in the urban river deciphered by metagenome assembly. Environmental Pollution 362, 125018. 10.1016/j.envpol.2024.125018

16. Liu, Y.Y., Chen, H.Y., Zhang, Y.X., Liu, C., Song, L.T., 2024. Metagenomics-resolved genomics provide novel ecological insights into resistome community coalescence of wastewater in river environment. Water Res. 267, 122473. 10.1016/j.watres.2024.122473

17. Narciso, A., Grenni, P., Spataro, F., De Carolis, C., Rauseo, J., Patrolecco, L., Garbini, G.L., Rolando, L., Iannelli, M.A., Bustamante, M.A., Alvarez-Alonso, C., Barra Caracciolo, A., 2024. Effects of sulfamethoxazole and copper on the natural microbial community from a fertilized soil. Appl. Microbiol. Biotechnol. 108, 516. 10.1007/s00253-024-13324-x

18. Schuijt, L.M., van Drimmelen, C.K.E., Buijse, L.L., van Smeden, J., Wu, D., Boerwinkel, M.C., Belgers, D.J.M., Matser, A.M., Roessink, I., Beentjes, K.K., Trimbos, K.B., Smidt, H., Van den Brink, P.J., 2024. Assessing ecological responses to exposure to the antibiotic sulfamethoxazole in freshwater mesocosms. Environmental Pollution 343, 123199. 10.1016/j.envpol.2023.123199

19. Singh, A., Pratap, S.G., Raj, A., 2024. Occurrence and dissemination of antibiotics and antibiotic resistance in aquatic environment and its ecological implications: a review. Environmental Science and Pollution Research 2024 31:35 31, 47505–47529. 10.1007/S11356-024-34355-X

20. Sonkar, V., Kashyap, A., Pallarés-Vega, R., Sasidharan, S.S., Modi, A., Uluseker, C., Jambu, S.C., Mohapatra, P.K., Larsen, J., Graham, D., Thatikonda, S., Kreft, J.-U., 2024. Field monitoring and hydraulic modelling quantify untreated wastewater as dominant source of AMR in a small river running through a big city. 10.1101/2024.12.21.629897

21. Wang, Z., Gu, Z., Wang, X., Sun, Z., Yan, C., Du, J., Yang, D., Xia, S., 2024. Impact of copper and sulfamethazine stress on microbial community and antibiotic resistance genes in denitrification systems: Comparison between heterotrophic and sulfur autotrophic denitrification processes. Journal of Water Process Engineering 66, 106078. 10.1016/j.jwpe.2024.106078

22. Zhang, Y., Liu, L., Liu, Y., Chen, L., Wang, J., Li, Y., Wang, K., Wang, W., 2024. Deciphering the natural and anthropogenic drivers on the fate and risk of antibiotics and antibiotic resistance genes (ARGs) in a typical river-estuary system, China. J. Hazard. Mater. 480, 136006. 10.1016/J.JHAZMAT.2024.136006

23. Gupta, V., Kumar, D., Dwivedi, A., Vishwakarma, U., Malik, D.S., Paroha, S., Mohan, N., Gupta, N., 2023. Heavy metal contamination in river water, sediment, groundwater and human blood, from Kanpur, Uttar Pradesh, India. Environ. Geochem. Health 45, 1807– 1818. 10.1007/s10653-022-01290-0

24. Kashyap, A., Nishil, B., Thatikonda, S., 2023. Experimental and numerical elucidation of the fate and transport of antibiotics in aquatic environment: A review. Environmental Monitoring and Assessment 2023 195:8 195, 942-. 10.1007/S10661-023-11482-5

25. Leão, I., Khalifa, L., Gallois, N., Vaz-Moreira, I., Klümper, U., Youdkes, D., Palmony, S., Dagai, L., Berendonk, T.U., Merlin, C., Manaia, C.M., Cytryn, E., 2023. Microbiome and Resistome Profiles along a Sewage-Effluent-Reservoir Trajectory Underline the Role of Natural Attenuation in Wastewater Stabilization Reservoirs. Appl. Environ. Microbiol. 89, e00170–23. 10.1128/aem.00170-23

26. Li, S., Chen, J., Zhao, J., Qi, W., Liu, H., 2023. The response of microbial compositions and functions to chronic single and multiple antibiotic exposure by batch experiment. Environ. Int. 179, 108181. 10.1016/j.envint.2023.108181

27. Baker, M., Williams, A.D., Hooton, S.P.T., Helliwell, R., King, E., Dodsworth, T., María Baena-Nogueras, R., Warry, A., Ortori, C.A., Todman, H., Gray-Hammerton, C.J., Pritchard, A.C.W., Iles, E., Cook, R., Emes, R.D., Jones, M.A., Kypraios, T., West, H., Barrett, D.A., Ramsden, S.J., Gomes, R.L., Hudson, C., Millard, A.D., Raman, S., Morris, C., Dodd, C.E.R., Kreft, J.U., Hobman, J.L., Stekel, D.J., 2022. Antimicrobial resistance in dairy slurry tanks: A critical point for measurement and control. Environ. Int. 169, 107516. 10.1016/J.ENVINT.2022.107516

28. Gupta, S., Graham, D.W., Sreekrishnan, T.R., Ahammad, S.Z., 2022. Effects of heavy metals pollution on the co-selection of metal and antibiotic resistance in urban rivers in UK and India. Environmental Pollution 306, 119326. 10.1016/j.envpol.2022.119326

29. Jamwal, P., Nayak, D., Urs, P.R., Thatey, M.Z., Gopinath, M., Idris, M., Lele, S., 2022. A multi-pronged approach to source attribution and apportionment of heavy metals in urban rivers. Ambio 51, 2182–2200. 10.1007/s13280-022-01734-y

30. Lee, J., Beck, K., Bürgmann, H., 2022. Wastewater bypass is a major temporary point-source of antibiotic resistance genes and multi-resistance risk factors in a Swiss river. Water Res. 208, 117827. 10.1016/j.watres.2021.117827

31. Leroy-Freitas, D., Machado, E.C., Torres-Franco, A.F., Dias, M.F., Leal, C.D., Araújo, J.C., 2022. Exploring the microbiome, antibiotic resistance genes, mobile genetic element, and potential resistant pathogens in municipal wastewater treatment plants in Brazil. Science of the Total Environment 842, 156773. 10.1016/j.scitotenv.2022.156773

32. Palkovicova, J., Sukkar, I., Delafuente, J., Valcek, A., Medvecky, M., Jamborova, I., Bitar, I., Phan, M.D., Millan, A.S., Dolejska, M., 2022. Fitness effects of blaCTX-M-15-harbouring F2:A1:B− plasmids on their native Escherichia coli ST131 H30Rx hosts. Journal of Antimicrobial Chemotherapy 77, 2960–2963. 10.1093/JAC/DKAC250

33. Peng, J., Zhao, L., Wang, Q., Song, W., Wang, Z., Li, J., Zhang, X., Yuan, F., 2022. Enhancing the Stability of Aerobic Granular Sludge Process Treating Municipal Wastewater by Adjusting Organic Loading Rate and Dissolved Oxygen Concentration. Separations 9, 228. 10.3390/separations9080228

34. Wu, Y., Wen, Q., Chen, Z., Fu, Q., Bao, H., 2022. Response of antibiotic resistance to the co-exposure of sulfamethoxazole and copper during swine manure composting. Science of the Total Environment 805, 150086. 10.1016/j.scitotenv.2021.150086

35. Zhang, W., Han, S., Zhang, D., Jin, X., Cao, E., Shan, B., Wei, D., 2022. Preliminary Study on the Dissolved Oxygen Recovery Process in Freshwater Ecosystems under the Coupling Effect of Oxygen-Consuming Pollutants and Temperature. ACS ES and T Water 2, 1639–1646. 10.1021/acsestwater.2c00150

36. Arya, S., Williams, A., Reina, S.V., Knapp, C.W., Kreft, J.U., Hobman, J.L., Stekel, D.J., 2021. Towards a general model for predicting minimal metal concentrations co-selecting for antibiotic resistance plasmids. Environmental Pollution 275, 116602. 10.1016/J.ENVPOL.2021.116602

37. Borsetto, C., Raguideau, S., Travis, E., Kim, D.W., Lee, D.H., Bottrill, A., Stark, R., Song, L., Cha, C.J., Pearson, J., Quince, C., Singer, A.C., Wellington, E.M.H., 2021. Impact of sulfamethoxazole on a riverine microbiome. Water Res. 201, 117382. 10.1016/j.watres.2021.117382

38. Chaturvedi, P., Singh, A., Chowdhary, P., Pandey, A., Gupta, P., 2021. Occurrence of emerging sulfonamide resistance (sul1 and sul2) associated with mobile integrons-integrase (intI1 and intI2) in riverine systems. Science of The Total Environment 751, 142217. 10.1016/J.SCITOTENV.2020.142217

39. de los Santos, E., Laviña, M., Poey, M.E., 2021. Strict relationship between class 1 integrons and resistance to sulfamethoxazole in Escherichia coli. Microb. Pathog. 161, 105206. 10.1016/j.micpath.2021.105206

40. Jane, S.F., Hansen, G.J.A., Kraemer, B.M., Leavitt, P.R., Mincer, J.L., North, R.L., Pilla, R.M., Stetler, J.T., Williamson, C.E., Woolway, R.I., Arvola, L., Chandra, S., DeGasperi, C.L., Diemer, L., Dunalska, J., Erina, O., Flaim, G., Grossart, H.P., Hambright, K.D., Hein, C., Hejzlar, J., Janus, L.L., Jenny, J.P., Jones, J.R., Knoll, L.B., Leoni, B., Mackay, E., Matsuzaki, S.I.S., McBride, C., Müller-Navarra, D.C., Paterson, A.M., Pierson, D., Rogora, M., Rusak, J.A., Sadro, S., Saulnier-Talbot, E., Schmid, M., Sommaruga, R., Thiery, W., Verburg, P., Weathers, K.C., Weyhenmeyer, G.A., Yokota, K., Rose, K.C., 2021. Widespread deoxygenation of temperate lakes. Nature 2021 594:7861 594, 66–70. 10.1038/s41586-021-03550-y

41. Larsson, D.G.J., Flach, C.F., 2021. Antibiotic resistance in the environment. Nature Reviews Microbiology 2021 20:5 20, 257–269. 10.1038/s41579-021-00649-x

42. Ott, A., Quintela-Baluja, M., Zealand, A.M., O’Donnell, G., Haniffah, M.R.M., Graham, D.W., 2021. Improved quantitative microbiome profiling for environmental antibiotic resistance surveillance. Environmental Microbiomes 16, 21. 10.1186/s40793-021-00391-0

43. Brown, P.C., Borowska, E., Peschke, R., Schwartz, T., Horn, H., 2020. Decay of elevated antibiotic resistance genes in natural river sediments after sedimentation of wastewater particles. Science of the Total Environment 705, 135861. 10.1016/j.scitotenv.2019.135861

44. Deng, C., Liu, X., Li, L., Shi, J., Guo, W., Xue, J., 2020. Temporal dynamics of antibiotic resistant genes and their association with the bacterial community in a water-sediment mesocosm under selection by 14 antibiotics. Environ. Int. 137, 105554. 10.1016/j.envint.2020.105554

45. Sabri, N.A., Schmitt, H., Van Der Zaan, B., Gerritsen, H.W., Zuidema, T., Rijnaarts, H.H.M., Langenhoff, A.A.M., 2020. Prevalence of antibiotics and antibiotic resistance genes in a wastewater effluent-receiving river in the Netherlands. J. Environ. Chem. Eng. 8, 102245. 10.1016/J.JECE.2018.03.004

46. Yang, J., Wang, H., Roberts, D.J., Du, H.N., Yu, X.F., Zhu, N.Z., Meng, X.Z., 2020. Persistence of antibiotic resistance genes from river water to tap water in the Yangtze River Delta. Science of The Total Environment 742, 140592. 10.1016/J.SCITOTENV.2020.140592

47. Bolyen, E., Rideout, J.R., Dillon, M.R., Bokulich, N.A., Abnet, C.C., Al-Ghalith, G.A., Alexander, H., Alm, E.J., Arumugam, M., Asnicar, F., Bai, Y., Bisanz, J.E., Bittinger, K., Brejnrod, A., Brislawn, C.J., Brown, C.T., Callahan, B.J., Caraballo-Rodríguez, A.M., Chase, J., Cope, E.K., Da Silva, R., Diener, C., Dorrestein, P.C., Douglas, G.M., Durall, D.M., Duvallet, C., Edwardson, C.F., Ernst, M., Estaki, M., Fouquier, J., Gauglitz, J.M., Gibbons, S.M., Gibson, D.L., Gonzalez, A., Gorlick, K., Guo, J., Hillmann, B., Holmes, S., Holste, H., Huttenhower, C., Huttley, G.A., Janssen, S., Jarmusch, A.K., Jiang, L., Kaehler, B.D., Kang, K. Bin, Keefe, C.R., Keim, P., Kelley, S.T., Knights, D., Koester, I., Kosciolek, T., Kreps, J., Langille, M.G.I., Lee, J., Ley, R., Liu, Y.X., Loftfield, E., Lozupone, C., Maher, M., Marotz, C., Martin, B.D., McDonald, D., McIver, L.J., Melnik, A. V., Metcalf, J.L., Morgan, S.C., Morton, J.T., Naimey, A.T., Navas-Molina, J.A., Nothias, L.F., Orchanian, S.B., Pearson, T., Peoples, S.L., Petras, D., Preuss, M.L., Pruesse, E., Rasmussen, L.B., Rivers, A., Robeson, M.S., Rosenthal, P., Segata, N., Shaffer, M., Shiffer, A., Sinha, R., Song, S.J., Spear, J.R., Swafford, A.D., Thompson, L.R., Torres, P.J., Trinh, P., Tripathi, A., Turnbaugh, P.J., Ul-Hasan, S., van der Hooft, J.J.J., Vargas, F., Vázquez-Baeza, Y., Vogtmann, E., von Hippel, M., Walters, W., Wan, Y., Wang, M., Warren, J., Weber, K.C., Williamson, C.H.D., Willis, A.D., Xu, Z.Z., Zaneveld, J.R., Zhang, Y., Zhu, Q., Knight, R., Caporaso, J.G., 2019. Reproducible, interactive, scalable and extensible microbiome data science using QIIME 2. Nature Biotechnology 2019 37:8 37, 852–857. 10.1038/s41587-019-0209-9

48. Chen, Y., Su, J.Q., Zhang, J., Li, P., Chen, H., Zhang, B., Gin, K.Y.H., He, Y., 2019. High-throughput profiling of antibiotic resistance gene dynamic in a drinking water river-reservoir system. Water Res. 149, 179–189. 10.1016/J.WATRES.2018.11.007

49. Cui, Q., Huang, Y., Wang, H., Fang, T., 2019. Diversity and abundance of bacterial pathogens in urban rivers impacted by domestic sewage. Environmental Pollution 249, 24–35. 10.1016/j.envpol.2019.02.094

50. Ju, F., Beck, K., Yin, X., Maccagnan, A., McArdell, C.S., Singer, H.P., Johnson, D.R., Zhang, T., Bürgmann, H., 2019. Wastewater treatment plant resistomes are shaped by bacterial composition, genetic exchange, and upregulated expression in the effluent microbiomes. ISME Journal 13, 346–360. 10.1038/s41396-018-0277-8

51. Phonsiri, V., Choi, S., Nguyen, C., Tsai, Y.L., Coss, R., Kurwadkar, S., 2019. Monitoring occurrence and removal of selected pharmaceuticals in two different wastewater treatment plants. SN Applied Sciences 2019 1:7 1, 798-. 10.1007/S42452-019-0774-Z

52. Xavier, J.C., Costa, P.E.S., Hissa, D.C., Melo, V.M.M., Falcão, R.M., Balbino, V.Q., Mendonça, L.A.R., Lima, M.G.S., Coutinho, H.D.M., Verde, L.C.L., 2019. Evaluation of the microbial diversity and heavy metal resistance genes of a microbial community on contaminated environment. Applied Geochemistry 105, 1–6. 10.1016/j.apgeochem.2019.04.012

53. Narciso-da-Rocha, C., Rocha, J., Vaz-Moreira, I., Lira, F., Tamames, J., Henriques, I., Martinez, J.L., Manaia, C.M., 2018. Bacterial lineages putatively associated with the dissemination of antibiotic resistance genes in a full-scale urban wastewater treatment plant. Environ. Int. 118, 179–188. 10.1016/j.envint.2018.05.040

54. Patrolecco, L., Rauseo, J., Ademollo, N., Grenni, P., Cardoni, M., Levantesi, C., Luprano, M.L., Caracciolo, A.B., 2018. Persistence of the antibiotic sulfamethoxazole in river water alone or in the co-presence of ciprofloxacin. Science of the Total Environment 640–641, 1438–1446. 10.1016/j.scitotenv.2018.06.025

55. Tran, N.H., Reinhard, M., Gin, K.Y.H., 2018. Occurrence and fate of emerging contaminants in municipal wastewater treatment plants from different geographical regions-a review. Water Res. 133, 182–207. 10.1016/J.WATRES.2017.12.029

56. He, B., He, J., Wang, J., Li, J., Wang, F., 2017. Abnormal pH elevation in the Chaobai River, a reclaimed water intake area. Environ. Sci. Process. Impacts 19, 111–122. 10.1039/C6EM00535G

57. Jia, S., Zhang, X.X., Miao, Y., Zhao, Y., Ye, L., Li, B., Zhang, T., 2017. Fate of antibiotic resistance genes and their associations with bacterial community in livestock breeding wastewater and its receiving river water. Water Res. 124, 259–268. 10.1016/j.watres.2017.07.061

58. Kang, J., Kwon, G., Nam, J.H., Kim, Y.O., Jahng, D., 2017. Carbon dioxide stripping from anaerobic digestate of food waste using two types of aerators. International Journal of Environmental Science and Technology 14, 1397–1408. 10.1007/s13762-017-1250-1

59. Quoc Tuc, D., Elodie, M.G., Pierre, L., Fabrice, A., Marie-Jeanne, T., Martine, B., Joelle, E., Marc, C., 2017. Fate of antibiotics from hospital and domestic sources in a sewage network. Science of The Total Environment 575, 758–766. 10.1016/J.SCITOTENV.2016.09.118

60. Callahan, B.J., McMurdie, P.J., Rosen, M.J., Han, A.W., Johnson, A.J.A., Holmes, S.P., 2016. DADA2: High-resolution sample inference from Illumina amplicon data. Nat. Methods 13, 581–583. 10.1038/nmeth.3869

61. Le, T.H., Ng, C., Chen, H., Yi, X.Z., Koh, T.H., Barkham, T.M.S., Zhou, Z., Gin, K.Y.H., 2016. Occurrences and characterization of antibiotic-resistant bacteria and genetic determinants of hospital wastewater in a tropical country. Antimicrob. Agents Chemother. 60, 7449–7456. 10.1128/AAC.01556-16;PAGE:STRING:ARTICLE/CHAPTER

62. Lehmann, K., Bell, T., Bowes, M.J., Amos, G.C.A., Gaze, W.H., Wellington, E.M.H., Singer, A.C., 2016. Trace levels of sewage effluent are sufficient to increase class 1 integron prevalence in freshwater biofilms without changing the core community. Water Res. 106, 163–170. 10.1016/J.WATRES.2016.09.035

63. Liu, A., Cao, H., Yang, Y., Ma, X., Liu, X., 2016. Combinational effects of sulfomethoxazole and copper on soil microbial community and function. Environmental Science and Pollution Research 23, 4235–4241. 10.1007/s11356-015-4892-x

64. Oliver, D.M., Bird, C., Burd, E., Wyman, M., 2016. Quantitative PCR profiling of Escherichia coli in livestock feces reveals increased population resilience relative to culturable counts under temperature extremes. Environ. Sci. Technol. 50, 9497–9505. 10.1021/acs.est.6b02657

65. Yang, L., He, J., Liu, Y., Wang, J., Jiang, L., Wang, G., 2016. Characteristics of change in water quality along reclaimed water intake area of the Chaobai River in Beijing, China. J. Environ. Sci. (China) 50, 93–102. 10.1016/j.jes.2016.05.023

66. Berendonk, T.U., Manaia, C.M., Merlin, C., Fatta-Kassinos, D., Cytryn, E., Walsh, F., Bürgmann, H., Sørum, H., Norström, M., Pons, M.N., Kreuzinger, N., Huovinen, P., Stefani, S., Schwartz, T., Kisand, V., Baquero, F., Martinez, J.L., 2015. Tackling antibiotic resistance: The environmental framework. Nat. Rev. Microbiol. 13, 310–317. 10.1038/nrmicro3439

67. Johnson, A.C., Keller, V., Dumont, E., Sumpter, J.P., 2015. Assessing the concentrations and risks of toxicity from the antibiotics ciprofloxacin, sulfamethoxazole, trimethoprim and erythromycin in European rivers. Science of the Total Environment 511, 747–755. 10.1016/j.scitotenv.2014.12.055

68. Kundral, S., Mudragada, R., Coro, E., Moncholi, M., Mora, N., Laha, S., Tansel, B., 2015. Improving Settling Characteristics of Pure Oxygen Activated Sludge by Stripping of Carbon Dioxide. Water Environment Research 87, 498–505. 10.2175/106143015x14212658614595

69. Li, Z., Sobek, A., Radke, M., 2015. Flume Experiments To Investigate the Environmental Fate of Pharmaceuticals and Their Transformation Products in Streams. Environ. Sci. Technol. 49, 6009–6017. 10.1021/ACS.EST.5B00273

70. Zhang, Q., He, X., Yan, T., 2015. Differential Decay of Wastewater Bacteria and Change of Microbial Communities in Beach Sand and Seawater Microcosms. Environ. Sci. Technol. 49, 8531–8540. 10.1021/acs.est.5b01879

71. Fisher, J.C., Levican, A., Figueras, M.J., McLellan, S.L., 2014. Population dynamics and ecology of Arcobacter in sewage. Front. Microbiol. 5. 10.3389/fmicb.2014.00525

72. Tanaka, T., Kawasaki, K., Daimon, S., Kitagawa, W., Yamamoto, K., Tamaki, H., Tanaka, M., Nakatsu, C.H., Kamagata, Y., 2014. A Hidden Pitfall in the Preparation of Agar Media Undermines Microorganism Cultivability. Appl. Environ. Microbiol. 80, 7659– 7666. 10.1128/AEM.02741-14

73. Barana, A.C., Lopes, D.D., Martins, T.H., Pozzi, E., Damianovic, M.H.R.Z., Del Nery, V., Foresti, E., 2013. Nitrogen and organic matter removal in an intermittently aerated fixed-bed reactor for post-treatment of anaerobic effluent from a slaughterhouse wastewater treatment plant. J. Environ. Chem. Eng. 1, 453–459. 10.1016/j.jece.2013.06.015

74. Kozich, J.J., Westcott, S.L., Baxter, N.T., Highlander, S.K., Schloss, P.D., 2013. Development of a dual-index sequencing strategy and curation pipeline for analyzing amplicon sequence data on the miseq illumina sequencing platform. Appl. Environ. Microbiol. 79, 5112–5120. 10.1128/AEM.01043-13

75. Michael, I., Rizzo, L., McArdell, C.S., Manaia, C.M., Merlin, C., Schwartz, T., Dagot, C., Fatta-Kassinos, D., 2013. Urban wastewater treatment plants as hotspots for the release of antibiotics in the environment: A review. Water Res. 47, 957–995. 10.1016/J.WATRES.2012.11.027

76. O’Flaherty, E., Gray, N.F., 2013. A comparative analysis of the characteristics of a range of real and synthetic wastewaters. Environmental Science and Pollution Research 2013 20:12 20, 8813–8830. 10.1007/S11356-013-1863-Y

77. Dong, H., Qiang, Z., Li, T., Jin, H., Chen, W., 2012. Effect of artificial aeration on the performance of vertical-flow constructed wetland treating heavily polluted river water. Journal of Environmental Sciences 24, 596–601. 10.1016/S1001-0742(11)60804-8

78. Jeanneau, L., Solecki, O., Wéry, N., Jardé, E., Gourmelon, M., Communal, P.Y., Jadas-Hécart, A., Caprais, M.P., Gruau, G., Pourcher, A.M., 2012. Relative decay of fecal indicator bacteria and human-associated markers: A microcosm study simulating wastewater input into seawater and freshwater. Environ. Sci. Technol. 46, 2375–2382. 10.1021/es203019y

79. Graham, D.W., Olivares-Rieumont, S., Knapp, C.W., Lima, L., Werner, D., Bowen, E., 2011. Antibiotic Resistance Gene Abundances Associated with Waste Discharges to the Almendares River near Havana, Cuba. Environ. Sci. Technol. 45, 418–424. 10.1021/ES102473Z

80. Knapp, C.W., Zhang, W., Sturm, B.S.M., Graham, D.W., 2010. Differential fate of erythromycin and beta-lactam resistance genes from swine lagoon waste under different aquatic conditions. Environmental Pollution 158, 1506–1512. 10.1016/J.ENVPOL.2009.12.020

81. Walters, E., McClellan, K., Halden, R.U., 2010. Occurrence and loss over three years of 72 pharmaceuticals and personal care products from biosolids–soil mixtures in outdoor mesocosms. Water Res. 44, 6011–6020. 10.1016/J.WATRES.2010.07.051

82. Radke, M., Lauwigi, C., Heinkele, G., Múrdter, T.E., Letzel, M., 2009. Fate of the Antibiotic Sulfamethoxazole and Its Two Major Human Metabolites in a Water Sediment Test. Environ. Sci. Technol. 43, 3135–3141. 10.1021/ES900300U

83. Engemann, C.A., Keen, P.L., Knapp, C.W., Hall, K.J., Graham, D.W., 2008. Fate of Tetracycline Resistance Genes in Aquatic Systems: Migration from the Water Column to Peripheral Biofilms. Environ. Sci. Technol. 42, 5131–5136. 10.1021/ES800238E

84. Knapp, C.W., Engemann, C.A., Hanson, M.L., Keen, P.L., Hall, K.J., Graham, D.W., 2008. Indirect Evidence of Transposon-Mediated Selection of Antibiotic Resistance Genes in Aquatic Systems at Low-Level Oxytetracycline Exposures. Environ. Sci. Technol. 42, 5348–5353. 10.1021/ES703199G

85. Li, J.P., Healy, M.G., Zhan, X.M., Rodgers, M., 2008. Nutrient removal from slaughterhouse wastewater in an intermittently aerated sequencing batch reactor. Bioresour. Technol. 99, 7644–7650. 10.1016/J.BIORTECH.2008.02.001

86. Battin, T.J., Kaplan, L.A., Newbold, J.D., Hansen, C.M.E., 2003. Contributions of microbial biofilms to ecosystem processes in stream mesocosms. Nature 2003 426:6965 426, 439–442. 10.1038/nature02152

87. Cox, B.A., 2003. A review of dissolved oxygen modelling techniques for lowland rivers. Science of the Total Environment 314–316, 303–334. 10.1016/S0048-9697(03)00062-7

88. OECD, 2001. Test No. 303: Simulation Test - Aerobic Sewage Treatment -- A: Activated Sludge Units; B: Biofilms. OECD Guidelines for the Testing of Chemicals, Section 3, OECD Guidelines for the Testing of Chemicals, Section 3. 10.1787/9789264070424-EN

89. Tsai, Y.L., Palmer, C.J., Sangermano, L.R., 1993. Detection of Escherichia coli in sewage and sludge by polymerase chain reaction. Appl. Environ. Microbiol. 59, 353–357. 10.1128/aem.59.2.353-357.1993

90. Frampton, E.W., Restaino, L., Blaszko, N., 1988. Evaluation of the β-Glucuronidase Substrate 5-Bromo-4-Chloro-3-Indolyl-β-D-Glucuronide (X-GLUC) in a 24-Hour Direct Plating Method for Escherichia coli. J. Food Prot. 51, 402–404. 10.4315/0362-028X-51.5.402

91. Wimpenny, J.W.T.., 1988. CRC handbook of laboratory model systems for microbial ecosystems.

92. Birkeland, J.M., Steinhaus, E.A., 1939. Selective Bacteriostatic Action of Sodium Lauryl Sulfate and of “Dreft.” Exp. Biol. Med. 40, 86–88. 10.3181/00379727-40-10314P

93. H. W. Streeter, B. Phelps, 1925. A study of the pollution and natural purification of the Illunois River. III: factors concerning the phenomena of oxidation and reaeration. Public Health Bulletin (Wash. D. C). 146, 75.

