## Supplementary Information for "Rapid Antimicrobial Resistance Decline Coupled with Microbial Community Shifts in Sewage Polluted River Mesocosms"

### Table of contents

Supplementary texts: 2

Supplementary tables: 18

Supplementary figures: 18

### S1 Sampling location for water and sediment samples

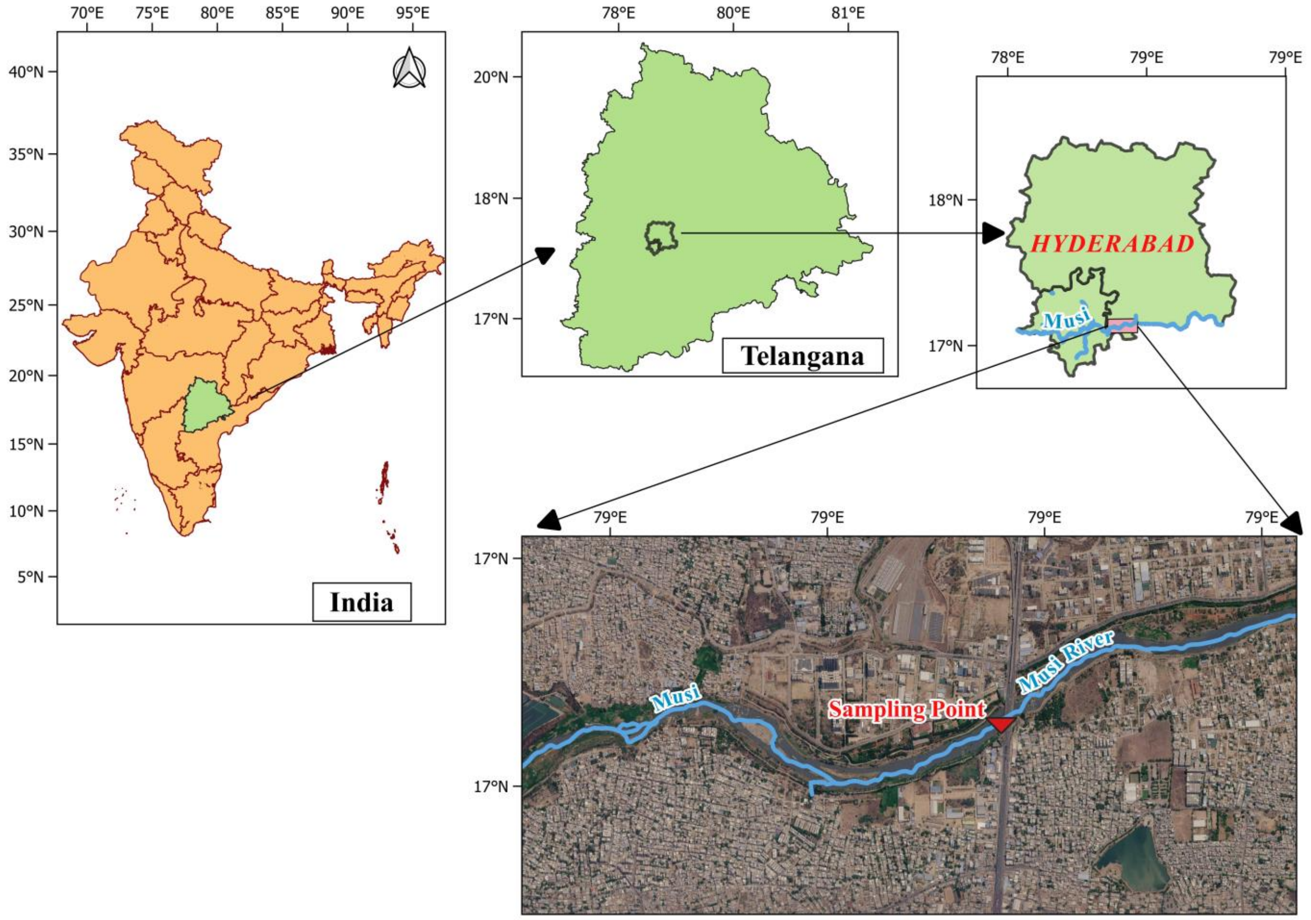


**Supplementary Figure S1.** Sampling point of polluted water and sediment for mesocosm set-up. The sampling location near Nagole Bridge within Hyderabad city was selected based on the previous Musi River study by Sonkar et al. (2024). The river at this location was heavily polluted with raw and treated wastewater.

### S2 Mesocosm experimental setup

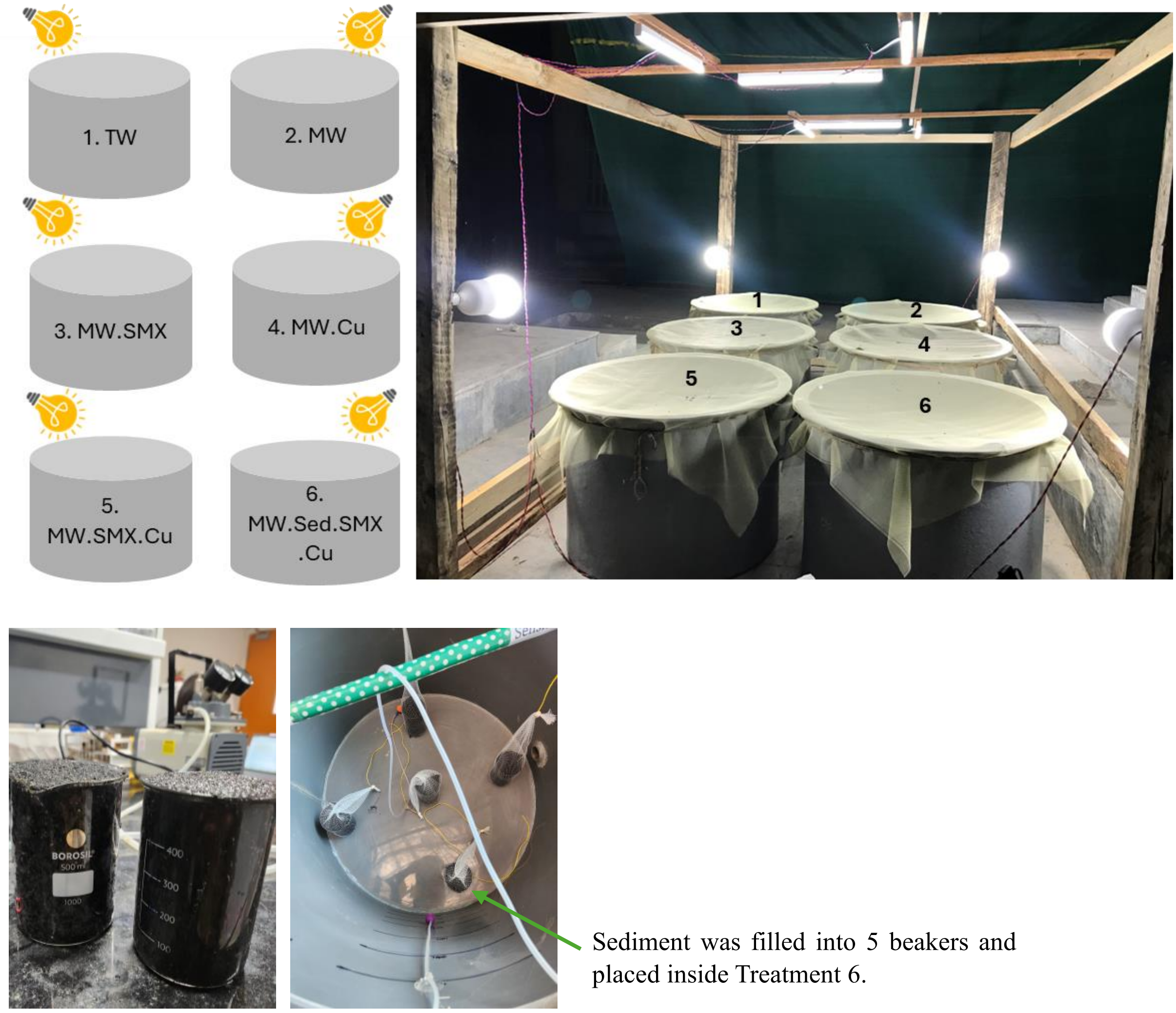


**Supplementary Figure S2.** Mesocosm experimental setup with schematic of the six different mesocosm treatment conditions as follows: Tap water (TW), Musi river water (MW), Musi river water with sulfamethoxazole (MW.SMX), Musi river water with copper (MW.Cu), Musi river water with sulfamethoxazole and copper (MW.SMX.Cu), and Musi river water with sediment, sulfamethoxazole and copper (MW.Sed.SMX.Cu). The operating conditions (per tank) were as follows: total volume 300 L, aeration 4 L/min, and illumination by about 6,000 lumens, with open-space temperature of 26.6 °C and 30 % humidity.

**Supplementary Table S1.** WQPs analyzed in water samples.

| S. No. | Parameter | Unit | Methods APHA (2017) |
|---|---|---|---|
| 1 | Temperature (Temp) | °C | |
| 2 | pH | | Potentiometry |
| 3 | Dissolved Oxygen (DO) | mg/L | Polarographic probe |
| 4 | Turbidity (TUR) | NTU | Nephelometry (2130-B) |
| 5 | Chemical Oxygen Demand (COD) | mg/L | Titrimetry (5220-C) |
| 6 | Total organic carbon (TOC) | mg/L | TOC analyzer |
| 7 | Ammonium ($NH_4^+ - N$) | mg/L | Ion Chromatography (Thomas et al., 2002) |
| 8 | Nitrite ($NO_2^- - N$) | mg/L | Ion Chromatography (Thomas et al., 2002) |
| 9 | Nitrate ($NO_3^- - N$) | mg/L | Ion Chromatography (Thomas et al., 2002) |
| 10 | Total Phosphate (TP) | mg/L | Vanadomolybdophosphoric Acid (4500-P C) |

**Supplementary Table S2.** Physicochemical parameters analyzed in sediment samples.

| S. No. | Parameter | Unit | Methods | References |
|---|---|---|---|---|
| 1 | pH | – | Potentiometry | (IS 2720 (Part26), 1987) |
| 2 | Ammonium ($NH_4^+ - N$ ) | mg/kg | Ion Chromatography | (Thomas et al., 2002) |
| 3 | Nitrate ($NO_3^- - N$ ) | mg/kg | Ion Chromatography | (Thomas et al., 2002) |
| 4 | Total Organic Carbon (TOC) | mg/kg | TOC analyzer | (Avramidis and Bekiari, 2021) |
| 5 | Total Nitrogen (TN) | mg/kg | TOC analyzer | (Avramidis and Bekiari, 2021) |
| 6 | Total Phosphate (TP) | mg/kg | Vanadomolybdophosphoric Acid (4500-P C) | APHA (2017) |

**Supplementary Table S3.** Oligonucleotides used for ARGs and MGE detection by qPCR reactions.

| **Target gene** | **Reference** | **Probe name** | **Oligonucleotide sequence 5'-3'** | **Conc. in reaction (nmol $L^{-1}$)** | **Amplicon size (bp)** | **Ann. Tª (ºC)** | **Standard Curve** |
|---|---|---|---|---|---|---|---|
| ***16S rDNA*** | (Muyzer et al., 1993) | q_338F | ACTCCTACGGGAGGCAGCAG | 200 | 198 | 60 | $R^2$= 0.996 Slope = -3.34 Efficiency (%) = 99.40 |
| | | q_518R | ATTACCGCGGCTGCTGG | | | | |
| ***qnrS*** | (Marti and Balcázar, 2013a) | q_qnrSrtF11 | GACGTGCTAACTTGCGTGAT | 200 | 117 | 60 | $R^2$= 0.998 Slope = -3.237 Efficiency (%) = 103.664 |
| | | q_qnrSrtR11 | TGGCATTGTTGGAAACTTG | | | | |
| ***uidA*** | (Frahm and Obst, 2003; Silkie et al., 2008) | q_uidA-F | CGGAAGCAACGCGTAAACTC | 200 | 90 | 62 | $R^2$ = 0.999 Slope = -3.51 Efficiency (%) = 92.64 |
| | | q_uidA-R | TGAGCGTCGCAGAACATTACA | | | | |
| ***aph(3")-Ib*** | (Liang et al., 2021) | q_aph(3")-Ib-F | ACTGGCAGGAGGAACAGGAGGGTG | 200 | 240 | 58 | $R^2$ = 1.000 Slope = -3.398 Efficiency (%) = 96.912 |
| | | q_aph(3")-Ib-R | CGTCCGGTAAGAAGTCGGGATTGA | | | | |
| ***sul2*** | (Pei et al., 2006) | q_sul2-F | TCCGGTGGAGGCCGGTATCTGG | 200 | 192 | 62 | $R^2$ = 0.999 Slope = -3.51 Efficiency (%) = 92.64 |
| | | q_sul2-R | CGGGAATGCCATCTGCCTTGAG | | | | |

| | | | | | | | |
|---|---|---|---|---|---|---|---|
| ***$bla_{NDM}$*** | Resistomap AY152 | q_blaNDM-F | GGCCACACCAGTGACAATATCA | 200 | 66 | 59 | $R^2$ = 0.997 Slope = -3.26 Efficiency (%) = 102.72 |
| | | q_blaNDM-R | CAGGCAGCCACCAAAAGC | | | | |
| ***$bla_{CTX-M}$*** | (Marti and Balcázar, 2013b) | q_CTXM-F | CTATGGCACCACCAACGATA | 200 | 104 | 57 | $R^2$ = 0.999 Slope = -3.51 Efficiency (%) = 90.79 |
| | | q_CTXM-R | ACGGCTTTCTGCCTTAGGTT | | | | |
| ***intI1*** | (Barraud et al., 2010) | q_intI-F | GATCGGTCGAATGCGTGT | 200 | 196 | 61 | $R^2$ = 0.999 Slope= -3.344 Efficiency (%) = 99.093 |
| | | q_intI-R | GCCTTGATGTTACCCGAGAG | | | | |
| ***ermF*** | Resistomap AY535 | q_ermF -F | TCTGATGCCCGAAATGTTCAAG | 300 | 170 | 57 | $R^2$ = 0.998 Slope = -3.227 Efficiency (%) = 104.113 |
| | | q_ermF-R | TGAAGGACAATTGAACCTCCCA | | | | |
| ***tetW*** | (Walsh et al., 2011) | q_tetW -F | CGGCAGCGCAAAGAGAAC | 200 | 59 | 57 | $R^2$ = 0.986 Slope = -3.28 Efficiency (%) = 101.94 |
| | | q_tetW -R | CGGGTCAGTATCCGCAAGTT | | | | |

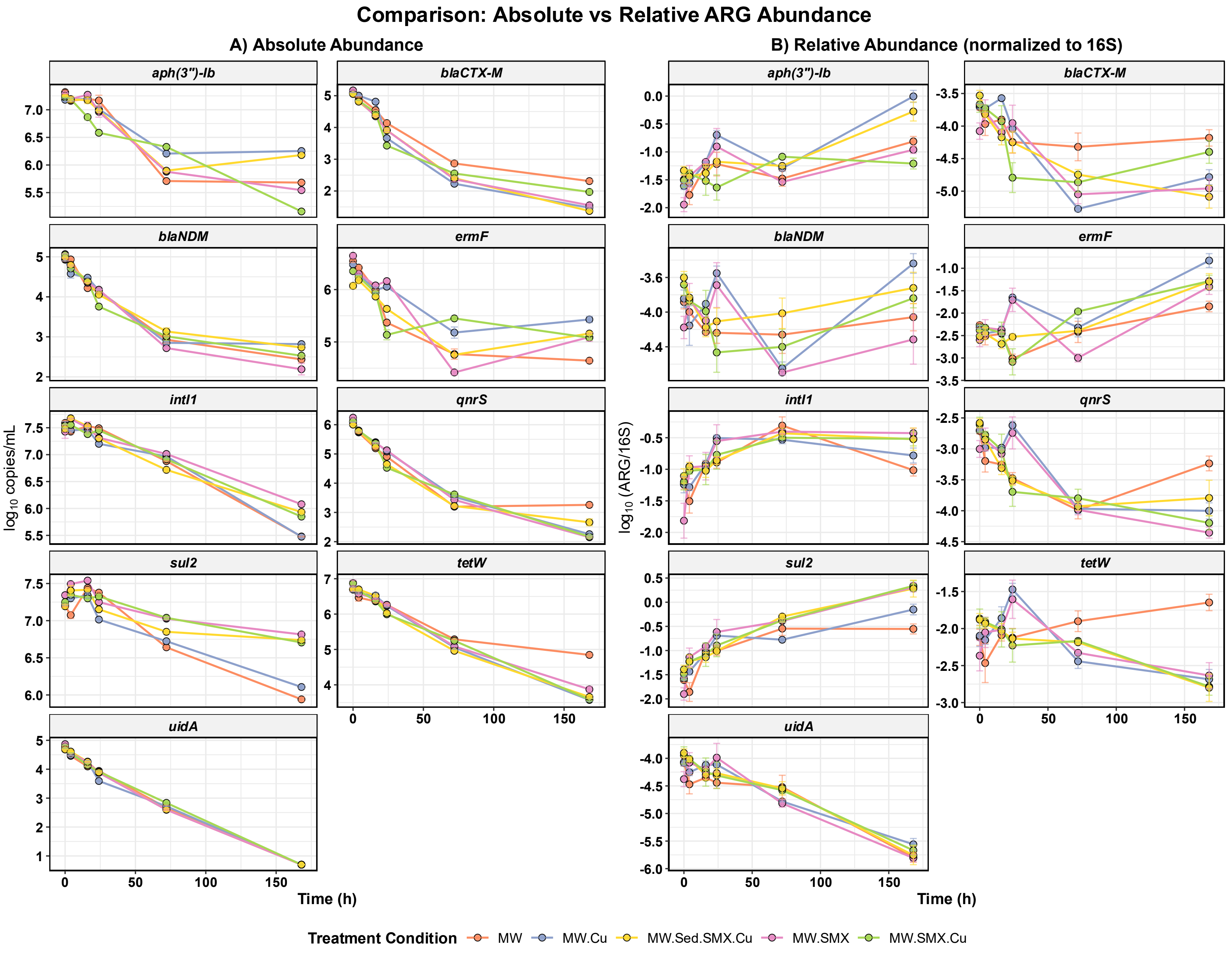
Comparison: Absolute vs Relative ARG Abundance
A) Absolute Abundance
B) Relative Abundance (normalized to 16S)
aph(3")-Ib
blaCTX-M
blaNDM
ermF
intI1
qnrS
sul2
tetW
uidA
log10 copies/mL
log10 (ARG/16S)
Time (h)
Treatment Condition
MW
MW.Cu
MW.Sed.SMX.Cu
MW.SMX
MW.SMX.Cu

**Supplementary Figure S3.** Comparison of **(A)** log absolute abundance ($log_{10}$ copies/mL) with **(B)** relative abundance (normalized to 16S rDNA gene copies). Absolute abundance shows that most genes drop by many orders of magnitude while all drop by at least one order of magnitude. The different extents of the drops mean that some genes increase in relative abundance, stay at similar levels or decline.

**Supplementary Table S4A.** Physicochemical WQPs of mesocosm treatments across sampling time points (mean ± SD, n = 2).

| Treatment | Time (h) | pH | Temp (° C) | TUR (NTU) | DO (mg/L) | COD (mg/L) | TOC (mg/L) | Ammonium (mg/L) | Nitrate (mg/L) | Nitrite (mg/L) | TP (mg/L) |
|---|---|---|---|---|---|---|---|---|---|---|---|
| **MW** | 0 | 7.835 ± 0.007 | 23.9 ± 0 | 126 ± 0 | 0 ± 0 | 78.933 ± 7.543 | 43.59 ± 0.651 | 36.728 ± 0.608 | 3.915 ± 0.018 | 0.032 ± 0.001 | 20.817 ± 0.735 |
| | 4 | 8.08 ± 0 | 25.35 ± 0.071 | 78.5 ± 0.707 | 0 ± 0 | 80 ± 6.034 | 41.38 ± 0.297 | 36.829 ± 1.037 | 1.046 ± 0.007 | 0.038 ± 0.001 | 22.898 ± 0.736 |
| | 16 | 8.31 ± 0.014 | 23.1 ± 0 | 51 ± 1.414 | 0.55 ± 0.071 | 54.4 ± 3.017 | 38.465 ± 0.445 | 34.35 ± 0.894 | 9.39 ± 0.043 | 0.078 ± 0.002 | 29.404 ± 1.105 |
| | 24 | 8.36 ± 0 | 24.2 ± 0 | 43.37 ± 0.453 | 2.35 ± 0.212 | 45.867 ± 0 | 37.915 ± 0.092 | 36.754 ± 0.501 | 0 ± 0 | 0.076 ± 0.001 | 15.873 ± 0.368 |
| | 72 | 8.54 ± 0 | 23.3 ± 0 | 21.645 ± 0.191 | 4.55 ± 0.778 | 24.534 ± 12.068 | 32.155 ± 0.474 | 38.373 ± 0.644 | 0 ± 0 | 0.12 ± 0.002 | 35.909 ± 2.208 |
| | 168 | 8.555 ± 0.007 | 24.3 ± 0 | 19.485 ± 0.247 | 4.45 ± 0.071 | 23.466 ± 28.661 | 40.065 ± 3.274 | 28.785 ± 0.177 | 0 ± 0 | 26.497 ± 0.26 | 27.843 ± 1.104 |
| **MW.Cu** | 0 | 7.825 ± 0.007 | 24.2 ± 0 | 123 ± 0 | 0 ± 0 | 84.267 ± 6.034 | 45.62 ± 0.042 | 38.499 ± 0.608 | 4.135 ± 0.049 | 0.035 ± 0.001 | 30.445 ± 0.375 |
| | 4 | 8.13 ± 0 | 25.05 ± 0.071 | 78 ± 1.414 | 0 ± 0 | 74.666 ± 1.508 | 44.215 ± 5.621 | 38.449 ± 0.25 | 40.61 ± 0.594 | 0.056 ± 0.004 | 31.49 ± 1.103 |
| | 16 | 8.41 ± 0 | 23.2 ± 0 | 48.97 ± 0.99 | 0.45 ± 0.071 | 51.2 ± 1.509 | 68.325 ± 0.078 | 38.575 ± 0.219 | 6.54 ± 0.028 | 0.04 ± 0.002 | 15.87 ± 0.368 |

| | | | | | | | | | | | |
|---|---|---|---|---|---|---|---|---|---|---|---|
| | 24 | 8.43 ± 0 | 23.8 ± 0 | 40.18 ± 0.014 | 2.55 ± 0.212 | 46.933 ± 1.508 | 36.545 ± 2.015 | 38.145 ± 0.969 | 0 ± 0 | 0.04 ± 0.002 | 31.49 ± 0.368 |
| | 72 | 8.56 ± 0 | 23.1 ± 0 | 20.1 ± 0.127 | 4.25 ± 0.071 | 20.267 ± 0 | 30.61 ± 0.495 | 37.13 ± 0.467 | 0 ± 0 | 0.114 ± 0.002 | 28.88 ± 0.368 |
| | 168 | 8.595 ± 0.007 | 24.1 ± 0 | 15.085 ± 0.191 | 4.9 ± 0 | 27.733 ± 7.543 | 36.46 ± 0.764 | 29.945 ± 0.177 | 0 ± 0 | 32.829 ± 0.348 | 31.23 ± 0.735 |
| **MW.SMX** | 0 | 7.82 ± 0.014 | 23.8 ± 0 | 123 ± 0 | 0 ± 0 | 96 ± 1.509 | 76.325 ± 0.375 | 34.755 ± 0.757 | 23.46 ± 0.834 | 0.045 ± 0.004 | 20.56 ± 0.368 |
| | 4 | 8.12 ± 0 | 25.3 ± 0 | 77.5 ± 0.707 | 0 ± 0 | 80 ± 0 | 71.08 ± 0 | 36.78 ± 0.75 | 6.99 ± 0.028 | 0.044 ± 0.001 | 28.88 ± 0.368 |
| | 16 | 8.42 ± 0 | 23.3 ± 0 | 51 ± 0 | 0.65 ± 0.071 | 61.867 ± 4.525 | 72.44 ± 1.556 | 35.055 ± 0.035 | 14.93 ± 0 | 0.074 ± 0.001 | 17.17 ± 0.735 |
| | 24 | 8.445 ± 0.007 | 24.05 ± 0.071 | 41.105 ± 0.658 | 2.9 ± 0 | 44.8 ± 1.509 | 66.565 ± 0.148 | 33.315 ± 0.431 | 0 ± 0 | 0.077 ± 0.003 | 22.898 ± 0.736 |
| | 72 | 8.575 ± 0.007 | 23.25 ± 0.071 | 28.765 ± 0.148 | 3.9 ± 0.141 | 35.2 ± 21.118 | 55.34 ± 0.41 | 38.095 ± 0.318 | 0 ± 0 | 0.064 ± 0.001 | 27.843 ± 0.368 |
| | 168 | 8.575 ± 0.007 | 24.2 ± 0 | 16.845 ± 0.007 | 4.7 ± 0 | 8.533 ± 4.525 | 52.37 ± 1.81 | 35.69 ± 0.283 | 0 ± 0 | 11.803 ± 0.261 | 32.53 ± 0.368 |
| **MW.SMX.Cu** | 0 | 7.805 ± 0.007 | 24 ± 0 | 126.5 ± 7.778 | 0 ± 0 | 91.733 ± 4.525 | 77.32 ± 2.659 | 35.36 ± 1.683 | 0 ± 0 | 0.05 ± 0.001 | 24.72 ± 1.103 |

| | | | | | | | | | | | |
|---|---|---|---|---|---|---|---|---|---|---|---|
| | 4 | 8.09 ± 0 | 25 ± 0 | 77 ± 0 | 0 ± 0 | 68.267 ± 1.508 | 70.99 ± 3.536 | 32.33 ± 0.679 | 0 ± 0 | 0.09 ± 0.004 | 32.27 ± 1.471 |
| | 16 | 8.385 ± 0.007 | 23.25 ± 0.071 | 49.19 ± 0.099 | 0.75 ± 0.071 | 56.534 ± 6.034 | 69.27 ± 2.376 | 34.935 ± 0.785 | 0 ± 0 | 0.072 ± 0.002 | 17.43 ± 0.368 |
| | 24 | 8.38 ± 0 | 23.7 ± 0 | 43.265 ± 0.191 | 3.35 ± 0.212 | 53.333 ± 4.525 | 67.97 ± 0.339 | 28.86 ± 1.146 | 0 ± 0 | 0.022 ± 0.001 | 29.92 ± 0.368 |
| | 72 | 8.51 ± 0 | 23.05 ± 0.071 | 24.43 ± 0.735 | 4.5 ± 0.141 | 37.334 ± 6.034 | 54.465 ± 0.544 | 39.895 ± 0.573 | 0 ± 0 | 0.135 ± 0.001 | 29.4 ± 0.368 |
| | 168 | 8.42 ± 0 | 24.1 ± 0 | 11.905 ± 0.148 | 4.55 ± 0.071 | 21.333 ± 4.525 | 53.3 ± 2.461 | 31.77 ± 0.608 | 0 ± 0 | 27.911 ± 0.086 | 34.61 ± 1.103 |
| **MW.Sed.SMX.Cu** | 0 | 7.805 ± 0.007 | 23.85 ± 0.071 | 120 ± 0 | 0 ± 0 | 83.2 ± 1.509 | 75.05 ± 0 | 38.75 ± 0.396 | 0 ± 0 | 0.058 ± 0.001 | 39.29 ± 0.368 |
| | 4 | 8.065 ± 0.007 | 24.9 ± 0 | 77 ± 0 | 0 ± 0 | 72.533 ± 4.525 | 70.755 ± 0.346 | 34.225 ± 2.001 | 0 ± 0 | 0.032 ± 0.001 | 34.09 ± 1.838 |
| | 16 | 8.35 ± 0 | 23.45 ± 0.071 | 51.5 ± 0.707 | 0.8 ± 0 | 57.6 ± 1.509 | 37.995 ± 1.025 | 31.035 ± 0.785 | 0 ± 0 | 0.073 ± 0.003 | 21.34 ± 0.735 |
| | 24 | 8.4 ± 0 | 23.45 ± 0.071 | 44.635 ± 0.049 | 2.75 ± 0.212 | 73.6 ± 3.017 | 72.295 ± 0.134 | 26.255 ± 0.389 | 0 ± 0 | 0.009 ± 0 | 33.31 ± 0.735 |
| | 72 | 8.585 ± 0.007 | 22.9 ± 0 | 26.75 ± 0.325 | 4.9 ± 0.283 | 20.267 ± 12.068 | 56.475 ± 0.615 | 35.565 ± 1.464 | 0 ± 0 | 0.068 ± 0.001 | 37.73 ± 0.368 |

| | | | | | | | | | | | |
|---|---|---|---|---|---|---|---|---|---|---|---|
| | 168 | 8.59 ± 0 | 24.3 ± 0 | 12.66 ± 0.17 | 4.8 ± 0 | 25.6 ± 1.509 | 51.215 ± 0.757 | 35.465 ± 0.318 | 0 ± 0 | 17.921 ± 0.304 | 33.31 ± 1.471 |

**Supplementary Table S4B.** Physicochemical properties of sediment samples.

| Treatment | Time (h) | pH | Ammonium (mg/L) | Nitrate (mg/L) | TP (mg/L) | TOC (mg/L) | TN (mg/L) |
|---|---|---|---|---|---|---|---|
| **MW.Sed.SMX.Cu** | 0 | 8.41 ± 0.014 | 3.765 ± 0.148 | 0.25 ± 0.071 | 25.41 ± 0.368 | 727.45 ± 1.202 | 156.1 ± 0.141 |
| | 24 | 8.49 ± 0.014 | 2.51 ± 0.014 | 0.495 ± 0.007 | 6.405 ± 0.021 | 454.795 ± 0.488 | 84.335 ± 0.332 |
| | 72 | 8.525 ± 0.007 | 1.835 ± 0.007 | 0.43 ± 0.014 | 5.74 ± 0.028 | 412.815 ± 0.262 | 80.075 ± 0.262 |
| | 168 | 8.585 ± 0.007 | 0.875 ± 0.021 | 0.425 ± 0.007 | 4.18 ± 0.028 | 365.75 ± 0.354 | 75.465 ± 0.035 |

**Supplementary Table S5A.** Comparison of first-order decay model performance for the 0–72 h and 0–168 h fitting windows across all target genes.

| Gene | Time (h) | $R^2$ | Adj $R^2$ | RMSE | AIC |
|---|---|---|---|---|---|
| ***16S rDNA*** | 0-168 h | 0.922 | 0.906 | 0.556 | 63.9 |
| | 0-72 h | 0.888 | 0.859 | 0.436 | 43.4 |
| ***aph(3")-Ib*** | 0-168 h | 0.718 | 0.659 | 0.832 | 88.1 |
| | 0-72 h | 0.828 | 0.783 | 0.535 | 53.7 |
| ***blaCTX-M*** | 0-168 h | 0.859 | 0.83 | 1.093 | 104.5 |
| | 0-72 h | 0.949 | 0.936 | 0.497 | 50 |
| ***blaNDM*** | 0-168 h | 0.828 | 0.792 | 0.876 | 91.2 |
| | 0-72 h | 0.959 | 0.949 | 0.339 | 30.9 |
| ***ermF*** | 0-168 h | 0.568 | 0.478 | 0.941 | 95.5 |
| | 0-72 h | 0.817 | 0.769 | 0.585 | 58.1 |
| ***intl1*** | 0-168 h | 0.961 | 0.953 | 0.294 | 25.7 |
| | 0-72 h | 0.874 | 0.841 | 0.21 | 6.9 |
| ***qnrS*** | 0-168 h | 0.858 | 0.828 | 1.156 | 107.8 |
| | 0-72 h | 0.963 | 0.953 | 0.443 | 44.3 |
| ***sul2*** | 0-168 h | 0.809 | 0.769 | 0.379 | 40.9 |
| | 0-72 h | 0.673 | 0.587 | 0.295 | 23.8 |
| ***tetW*** | 0-168 h | 0.89 | 0.868 | 0.823 | 87.5 |
| | 0-72 h | 0.929 | 0.911 | 0.434 | 43.2 |
| ***uidA*** | 0-168 h | 0.992 | 0.991 | 0.298 | 26.5 |
| | 0-72 h | 0.977 | 0.971 | 0.258 | 17.1 |

**Supplementary Table S5B.** Tests of the common-slope assumption for ANCOVA first-order decay models based on (Time × Treatment) interaction analyses**.**

| Gene | F-statistic | df (Num, Den) | Interaction p-value | Slopes equal (p > 0.05)? |
|---|---|---|---|---|
| ***16S rDNA*** | 0.41 | (4, 15) | 0.79 | Yes (Common slope validated) |
| ***uidA*** | 0.51 | (4, 15) | 0.72 | Yes (Common slope validated) |
| ***sul2*** | 0.42 | (4, 15) | 0.79 | Yes (Common slope validated) |
| ***intl1*** | 1.11 | (4, 15) | 0.38 | Yes (Common slope validated) |
| ***tetW*** | 5.23 | (4, 15) | 0.007 | No |
| ***blaNDM*** | 0.62 | (4, 15) | 0.65 | Yes (Common slope validated) |
| ***ermF*** | 2.26 | (4, 15) | 0.11 | Yes (Common slope validated) |
| ***qnrS*** | 1.84 | (4, 15) | 0.17 | Yes (Common slope validated) |
| ***aph(3”)-Ib*** | 4.90 | (4, 15) | 0.009 | No |
| ***blaCTX-M*** | 0.74 | (4, 15) | 0.57 | Yes (Common slope validated) |

**Supplementary Table S5C.** Per treatment decay rate estimates ($h^{-1}$) with SE. For plots of these data see **Supplementary Figure S4**.

| Gene | MW | MW.Cu | MW.SMX | MW.SMX.Cu | MW.Sed.SMX.Cu |
|---|---|---|---|---|---|
| ***16S rDNA*** | −0.0546 ± 0.0043 | −0.0413 ± 0.0132 | −0.0504 ± 0.0143 | −0.0410 ± 0.0016 | −0.0491 ± 0.0053 |
| ***uidA*** | −0.0632 ± 0.0038 | −0.0623 ± 0.0090 | −0.0696 ± 0.0046 | −0.0600 ± 0.0049 | −0.0672 ± 0.0025 |
| ***sul2*** | −0.0183 ± 0.0101 † | −0.0184 ± 0.0049 | −0.0136 ± 0.0052 † | −0.0086 ± 0.0029 † | −0.0155 ± 0.0058 † |
| ***intI1*** | −0.0216 ± 0.0046 | −0.0181 ± 0.0037 | −0.0178 ± 0.0046 | −0.0205 ± 0.0026 | −0.0285 ± 0.0047 |
| ***tetW*‡** | −0.0852 ± 0.0101 | −0.0530 ± 0.0042 | −0.0545 ± 0.0035 | −0.0499 ± 0.0062 | −0.0580 ± 0.0050 |
| ***blaNDM*** | −0.0640 ± 0.0067 | −0.0640 ± 0.0044 | −0.0721 ± 0.0037 | −0.0610 ± 0.0114 | −0.0570 ± 0.0060 |
| ***ermF*** | −0.0556 ± 0.0115 | −0.0385 ± 0.0040 | −0.0678 ± 0.0077 | −0.0275 ± 0.0177 † | −0.0450 ± 0.0034 |
| ***qnrS*** | −0.1047 ± 0.0034 | −0.0787 ± 0.0038 | −0.0848 ± 0.0057 | −0.0770 ± 0.0143 | −0.0877 ± 0.0081 |
| ***aph(3")-Ib*‡** | −0.0731 ± 0.0132 | −0.0330 ± 0.0046 | −0.0465 ± 0.0061 | −0.0283 ± 0.0071 | −0.0445 ± 0.0050 |
| ***blaCTX-M*** | −0.0708 ± 0.0042 | −0.0946 ± 0.0136 | −0.0864 ± 0.0061 | −0.0794 ± 0.0159 | −0.0833 ± 0.0047 |

Each value is the SE from a simple linear regression analysis within a single treatment (n=5 time points; df=3). This table's primary purpose was to demonstrate treatment equivalence, rather than to report new kinetic parameters.

† $CI_{95}$ included zero: decay not significant at the individual treatment level (n=5, df=3).

‡ ANCOVA Time × Condition interaction significant ($p < 0.05$; **Supplementary Table S5B**), indicating genuine treatment-dependent decay rates; treatment-specific estimates for these two genes are reported in the main text (**Table 1**) instead of a pooled value. † 95% CI includes zero.

**Supplementary text 1:**

Copper addition alone (MW.Cu) did not consistently shift gene decay rates relative to MW control. For *sul2* ($k$ = -0.0184 ± 0.0049 vs. MW: $k$ = −0.0183 ± 0.0101 $h^{-1}$), *blaNDM* (0.0640 ± 0.0044 vs. −0.0640 ± 0.0067 $h^{-1}$), and *intI1* (−0.0181 ± 0.0037 vs. −0.0216 ± 0.0046 $h^{-1}$), the rate in MW.Cu was indistinguishable from MW. In contrast, *tetW* and *aph(3")-Ib* showed a statistically confirmed treatment-dependent decay rate (ANCOVA Time × Condition interaction, p = 0.0076 and p = 0.0099, respectively; **Supplementary Table S5B**), with decay under MW.Cu (*tetW*: −0.053 ± 0.004 $h^{-1}$; *aph(3")-Ib*: −0.033 ± 0.005 $h^{-1}$) markedly slower than under MW alone (*tetW*: −0.085 ± 0.010 $h^{-1}$; *aph(3")-Ib*: −0.073 ± 0.013 $h^{-1}$). However, the MW control estimates for the target genes carried the largest SEs among all the treatments and broadly overlapped the pooled decay rate for all treatments.

Sulfamethoxazole addition (MW.SMX) likewise showed no consistent effect on ARG decay rates across most genes. The sole exception was *sul2*, where the slope in MW.SMX ($k$ = −0.0136 ± 0.0052 $h^{-1}$) was less steep than the MW control (−0.0183 ± 0.0101 $h^{-1}$), and its 95% CI marginally included zero ($CI_{95}$ = [−0.030, +0.003]). Nevertheless, the substantial overlap in SE intervals between MW.SMX and MW, coupled with the wide uncertainty in the MW control estimate, means that the lower rate estimate is not statistically robust. For the remaining genes, the decay rates under the MW.SMX condition remained consistent with both the MW control and the pooled estimate (**Supplementary Figure S4**). No gene exhibited a clear, systematic shift toward faster or slower decay in the presence of sulfamethoxazole.

The combined copper and sulfamethoxazole addition (MW.SMX.Cu) did not result in any additive or synergistic effects distinguishable from estimation uncertainty. Among all gene-treatment combinations, *sul2* had the lowest decay rate under this condition ($k$ = −0.0086 ± 0.0029 $h^{-1}$; $CI_{95}$ = [−0.018, +0.0006])). Since the upper CI approached zero, the net removal of *sul2* was negligible in this treatment. Similarly, *ermF* yielded a rate of −0.028 ± 0.018 $h^{-1}$ with its 95% CI spanning into positive values ($CI_{95}$ = [−0.084, +0.029]). For the remaining genes, decay rates in the MW.SMX.Cu treatment remained within the pooled 95% CI band. Given the limited sample size per treatment, these isolated patterns for *sul2* and *ermF* should be interpreted cautiously.

Finally, the inclusion of sediment together with both stressors (MW.Sed.SMX.Cu) produced no discernible difference in decay kinetics compared with other treatment conditions. In this condition, the 95% CI for all ten gene-specific rates excluded zero, confirming measurable declines in every case. Moreover, the ±1 SE intervals for these rates consistently overlapped with the pooled rate (**Supplementary Figure S4**). The decay rate for *intI1* was slightly steeper in this treatment ($-0.0285 \pm 0.0047$ $h^{-1}$) than the MW control ($-0.0216 \pm 0.0046$ $h^{-1}$). Although the magnitude of this difference is small and the overlapping SE intervals prevent any firm conclusions.

These treatment-level patterns were consistent with the ANCOVA test of slope homogeneity (**Supplementary Table S6**), which identified *tetW* and *aph(3")-Ib* as the only genes exhibiting statistically significant treatment-dependent decay kinetics; all other genes support the pooled common-slope model reported in **Table 1**.

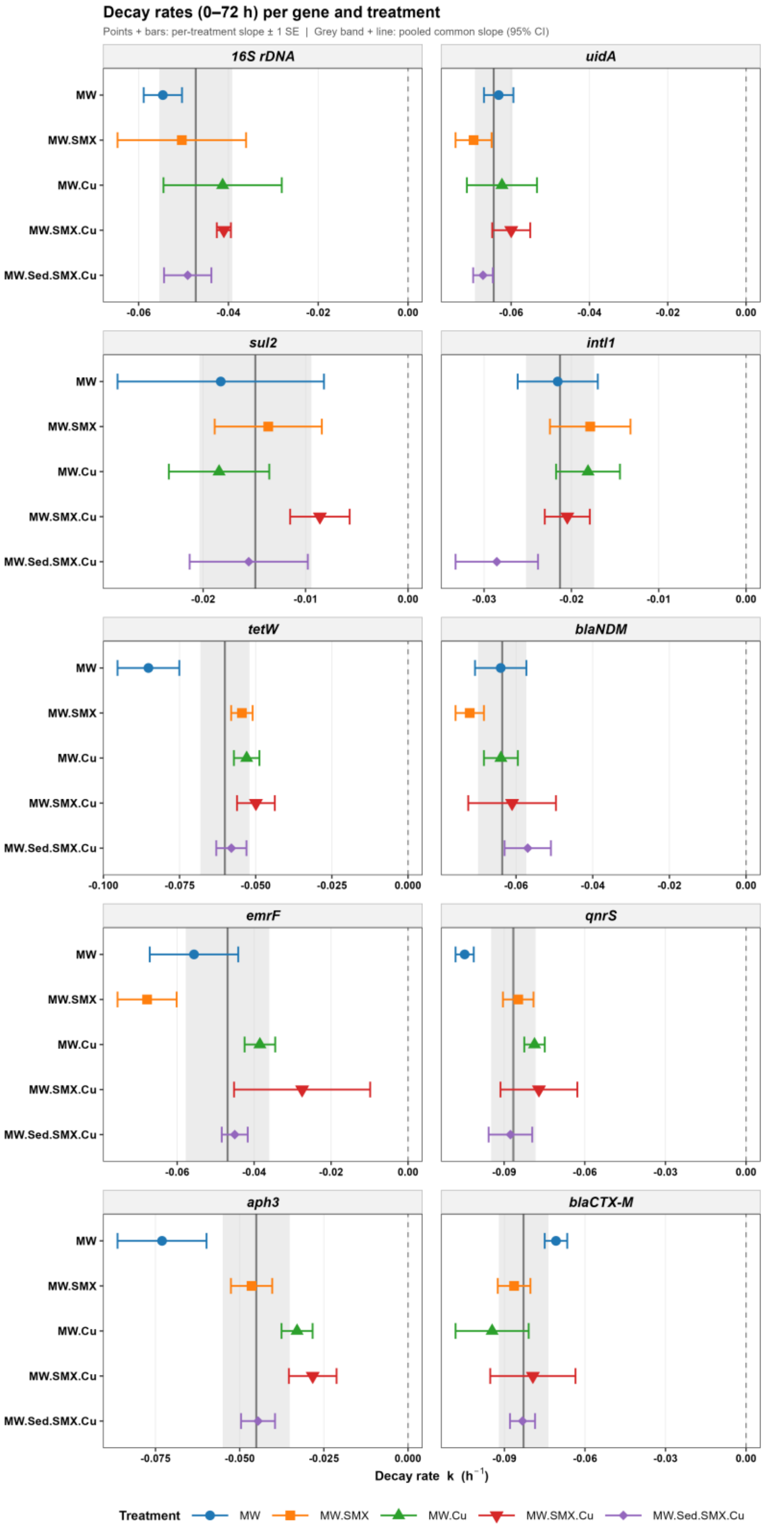


**Supplementary Figure S4.** First-order decay rates (0–72 h) of bacterial markers, mobile genetic elements, and ARGs across mesocosm treatments showing treatment-specific variation in gene dynamics. The gray line shows the pooled decay rate and the band its 95% CI. For the data plotted here, see **Supplementary Table S5(C)**.

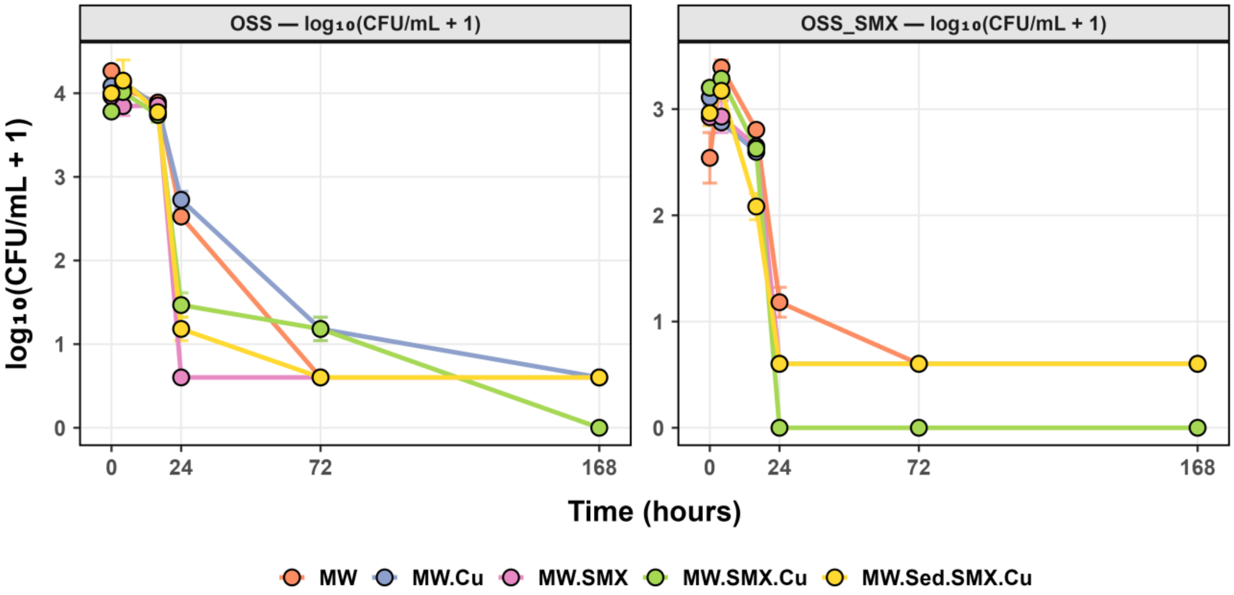


**Supplementary Figure S5.** Temporal changes in bacterial colony forming units on OECD Synthetic Sewage (OSS) agar and sulfamethoxazole-supplemented agar (OSS_SMX; 6.4 mg/L) across mesocosm treatments during 168 h time.

**Supplementary Table S6.** PCA loadings of ARGs on PC1 and PC2.

| **Gene** | **PC1 loading** | **PC2 loading** |
|---|---|---|
| *sul2* | -0.32 | 0.83 |
| *tetW* | -0.39 | 0.123 |
| *blaNDM* | -0.39 | -0.12 |
| *ermF* | -0.35 | -0.50 |
| *qnrS* | -0.40 | -0.04 |
| *aph(3")-Ib* | -0.37 | -0.16 |
| *blaCTX-M* | -0.39 | -0.01 |

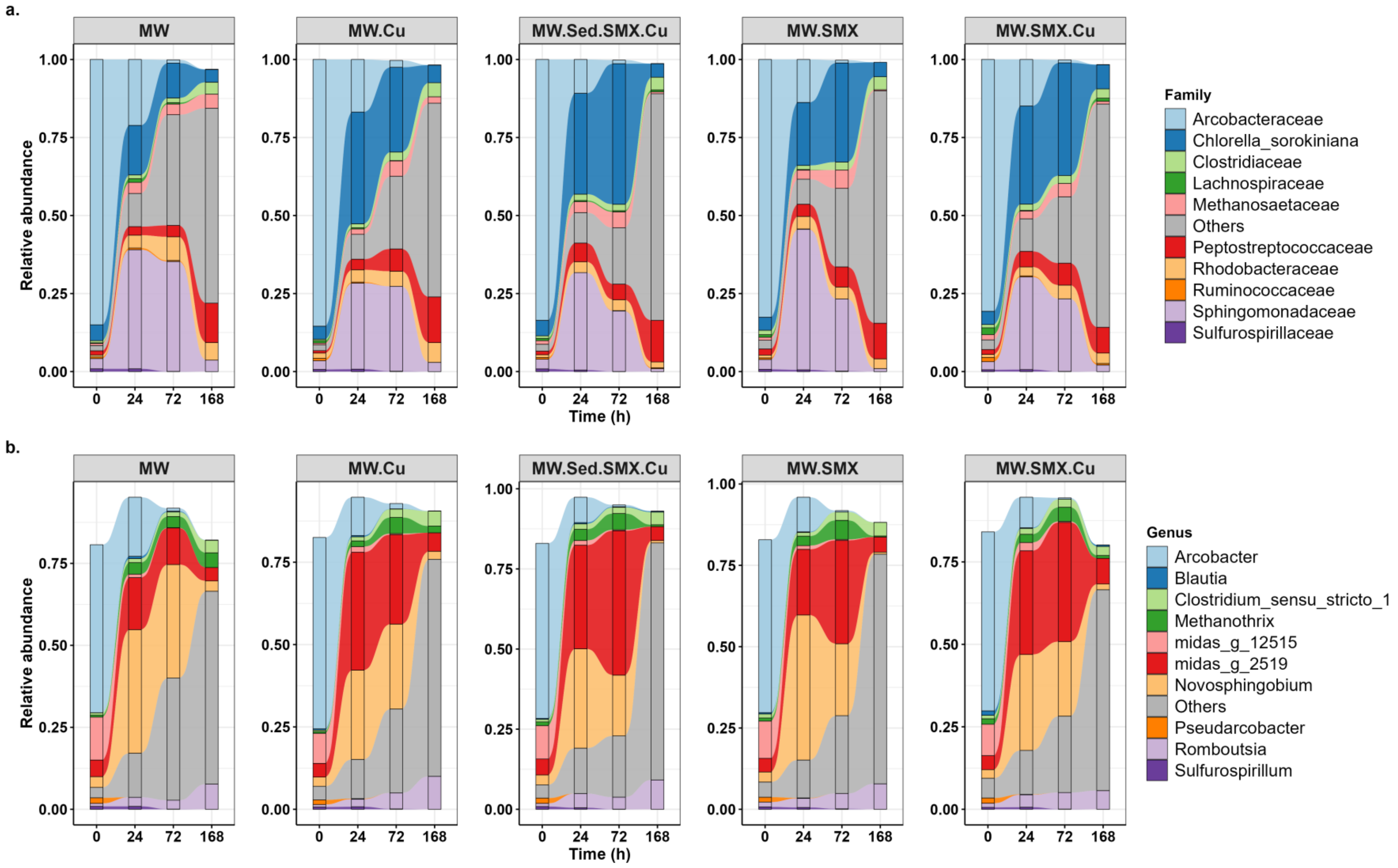


**Supplementary Figure S6.** Temporal changes in relative microbial community composition across mesocosm treatments at (a) family and (b) genus levels over the 168 h period. See **Supplementary Figure S7** for changes in absolute abundance.

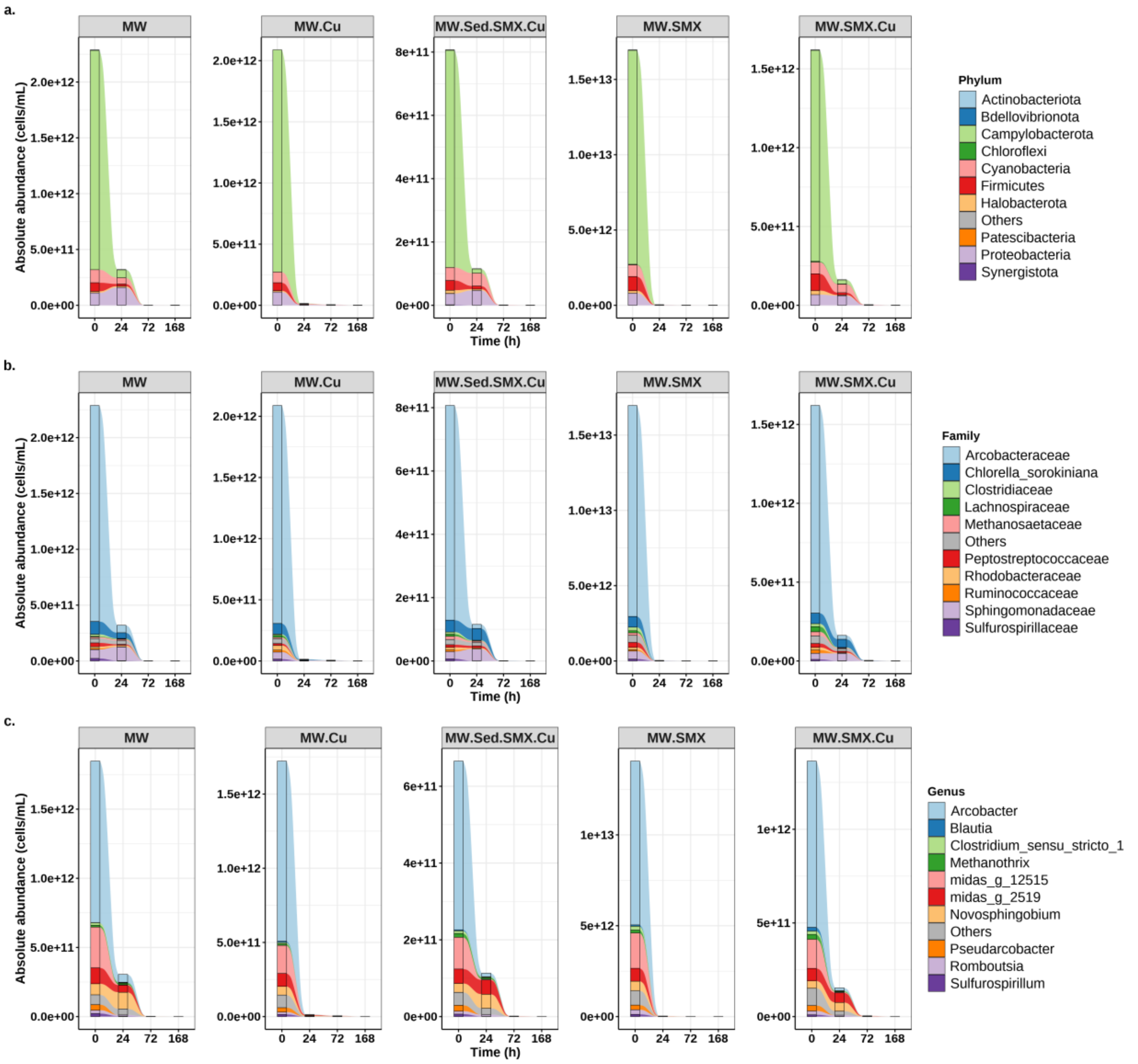


**Supplementary Figure S7.** Temporal changes in absolute microbial abundance under quantitative microbiome profiling (QMP) across mesocosm treatments at (a) phylum, (b) family, and (c) genus levels. While some relative abundances increase (**Figure 4, Supplementary Figure S6**), the absolute abundance of taxa generally drops drastically with few microbes left at the end.

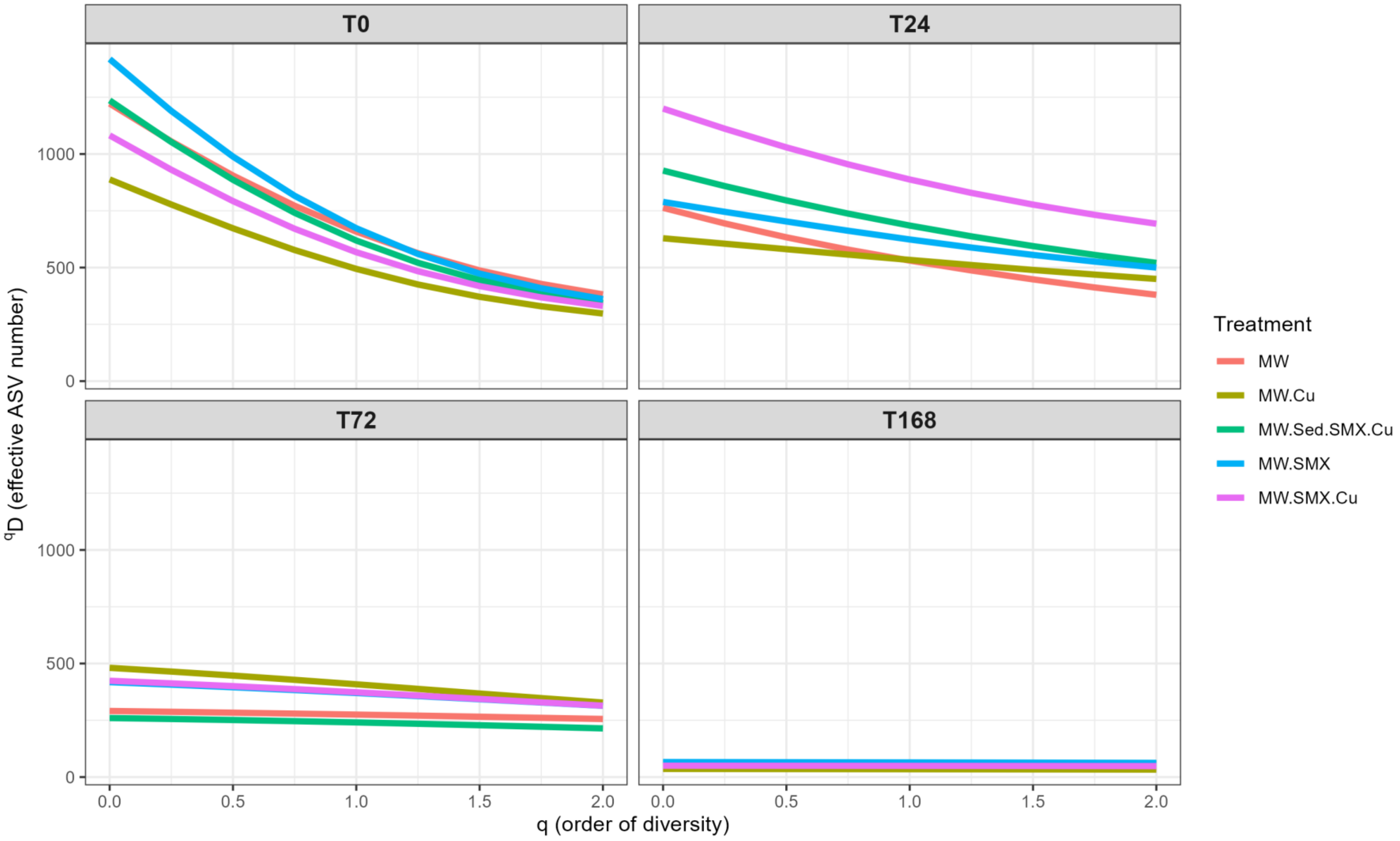


**Supplementary Figure S8.** Changes of Hill diversity profiles based on absolute abundances (QMP) over time showing progressive decline in microbial alpha diversity across all mesocosm treatments while evenness increases. Order of diversity q=0 is richness, q=1 is exponential Shannon index and q=2 is inverse Simpson index. See **Supplementary Table S7** for the numbers.

**Supplementary Table S7.** Temporal changes in alpha diversity (Hill numbers $^{q}D$) across mesocosms based on absolute abundances (QMP). For plots of these data see **Supplementary Figure S8**.

| **Treatment** | **Time (h)** | **$^{q}D$ value** | | |
|---|---|---|---|---|
| | | **q = 0** | **q = 1** | **q= 2** |
| **MW** | 0 | 1222 | 657.99 | 381.86 |
| | 24 | 763 | 530.73 | 379.89 |
| | 72 | 291 | 275.06 | 255.8 |
| | 168 | 64 | 62.16 | 59.76 |
| **MW.Cu** | 0 | 888 | 494.12 | 297.56 |
| | 24 | 629 | 533.71 | 449.93 |
| | 72 | 481 | 408.49 | 328.1 |
| | 168 | 36 | 34.62 | 32.96 |
| **MW.SMX** | 0 | 1418 | 673.14 | 361.28 |
| | 24 | 789 | 623.69 | 500.06 |
| | 72 | 418 | 370.14 | 313.81 |
| | 168 | 66 | 64.4 | 62.23 |
| **MW.SMX.Cu** | 0 | 1082 | 567.59 | 330.89 |
| | 24 | 1200 | 887.77 | 693 |
| | 72 | 424 | 373.39 | 314.38 |
| | 168 | 50 | 49.3 | 48.29 |
| **MW.Sed.SMX.Cu** | 0 | 1236 | 619.05 | 344.02 |
| | 24 | 927 | 684.98 | 520.63 |
| | 72 | 260 | 240.69 | 214.37 |
| | 168 | 63 | 62.29 | 61.23 |

**Supplementary Table S8.** Summary of db-RDA models for ARG and microbial community (MC) composition. The model with time only was always significant while the model with treatment only never was.

| Response | Model | $R^2$ | Adj. $R^2$ | F | p-value | Significance |
|---|---|---|---|---|---|---|
| **ARG** | Full (Time + Treatment) | 0.66 | 0.54 | 5.63 | 0.001 | ** |
| | Time only | 0.59 | 0.57 | 25.23 | 0.001 | ** |
| | Treatment only | 0.06 | -0.17 | 0.73 | 0.64 | ns |
| | Treatment \| Time (unique) | 0.06 | -0.02 | 0.73 | 0.354 | ns |
| | Time \| Treatment (unique) | 0.59 | 0.728 | 25.23 | 0.001 | ** |
| **MC (Phylum)** | Full (Time + Treatment) | 0.81 | 0.74 | 12.27 | 0.001 | ** |
| | Time only | 0.78 | 0.77 | 59.16 | 0.001 | ** |
| | Treatment only | 0.02 | -0.23 | 0.54 | 0.78 | ns |
| | Treatment \| Time (unique) | 0.02 | -0.02 | 0.548 | 0.07 | ns |
| | Time \| Treatment (unique) | 0.78 | 0.97 | 59.16 | 0.001 | ** |
| **MC (Family)** | Full (Time + Treatment) | 0.64 | 0.52 | 5.163 | 0.001 | ** |
| | Time only | 0.60 | 0.580 | 23.99 | 0.001 | ** |
| | Treatment only | 0.04 | -0.20 | 0.45 | 0.933 | ns |
| | Treatment \| Time (unique) | 0.04 | -0.05 | 0.45 | 0.418 | ns |
| | Time \| Treatment (unique) | 0.60 | 0.73 | 23.99 | 0.001 | ** |
| **MC (Genus)** | Full (Time + Treatment) | 0.63 | 0.50 | 4.84 | 0.001 | ** |
| | Time only | 0.57 | 0.55 | 22.07 | 0.001 | ** |
| | Treatment only | 0.05 | -0.19 | 0.53 | 0.914 | ns |
| | Treatment \| Time (unique) | 0.05 | -0.05 | 0.53 | 0.122 | ns |
| | Time \| Treatment (unique) | 0.57 | 0.69 | 22.07 | 0.001 | ** |

'Unique' refers to the proportion of total variance explained solely and independently by a given factor (ARGs/Community, Treatment, or Time) after partialling out (removing) all other model covariates.

**Supplementary Table S9.** Scores and vector loadings for phylum, family and genus level associated with the constrained ordination axes from the db-RDA analysis of microbial community composition across the mesocosms based on the model MC: Full (Time + Treatment). Plotted in **Supplementary Figure S9 and S10** for family and genus level.

| **a. Phylum** | | |
|---|---|---|
| **Taxon** | **CAP1** | **CAP2** |
| *Cyanobacteriota* | -0.453 | 0.194 |
| *Pseudomonadota* | 0.217 | -0.055 |
| *Bacillota* | 0.073 | 0.103 |
| *Campylobacterota* | -3.126 | -0.060 |
| *Halobacteriota* | -0.480 | -0.010 |
| *Actinobacteriota* | 1.202 | -0.081 |
| *Chloroflexota* | 1.145 | 0.227 |
| *Euryarchaeota* | 0.516 | 0.101 |
| *Desulfobacterota* | 0.811 | -0.236 |
| *Patescibacteriota* | 1.045 | -0.167 |
| *Bdellovibrionota* | -0.583 | -0.209 |
| *Bacteroidota* | -0.366 | 0.191 |
| **b. Family** | | |
| *Arcobacteraceae* | -2.199 | 0.013 |
| *Rickettsiaceae* | 1.093 | 0.023 |
| *Microtrichaceae* | 1.216 | 0.031 |
| *Ilumatobacteraceae* | 1.115 | -0.007 |
| *Hyphomicrobiaceae* | 1.099 | 0.052 |
| *Micavibrionaceae* | 1.433 | 0.086 |
| *Sulfurospirillaceae* | -1.302 | 0.003 |
| **c. Genus** | | |
| *Arcobacter* | -1.839 | -0.079 |
| Ca. Megaira | 1.111 | -0.026 |
| *midas_g_120* | 1.168 | 0.031 |
| *Hyphomicrobium* | 1.105 | 0.079 |
| *midas_g_22103* | 1.412 | 0.055 |
| *midas_g_12515* | -1.480 | 0.162 |

| | | |
|---|---|---|
| *Sulfurospirillum* | -1.126 | 0.014 |

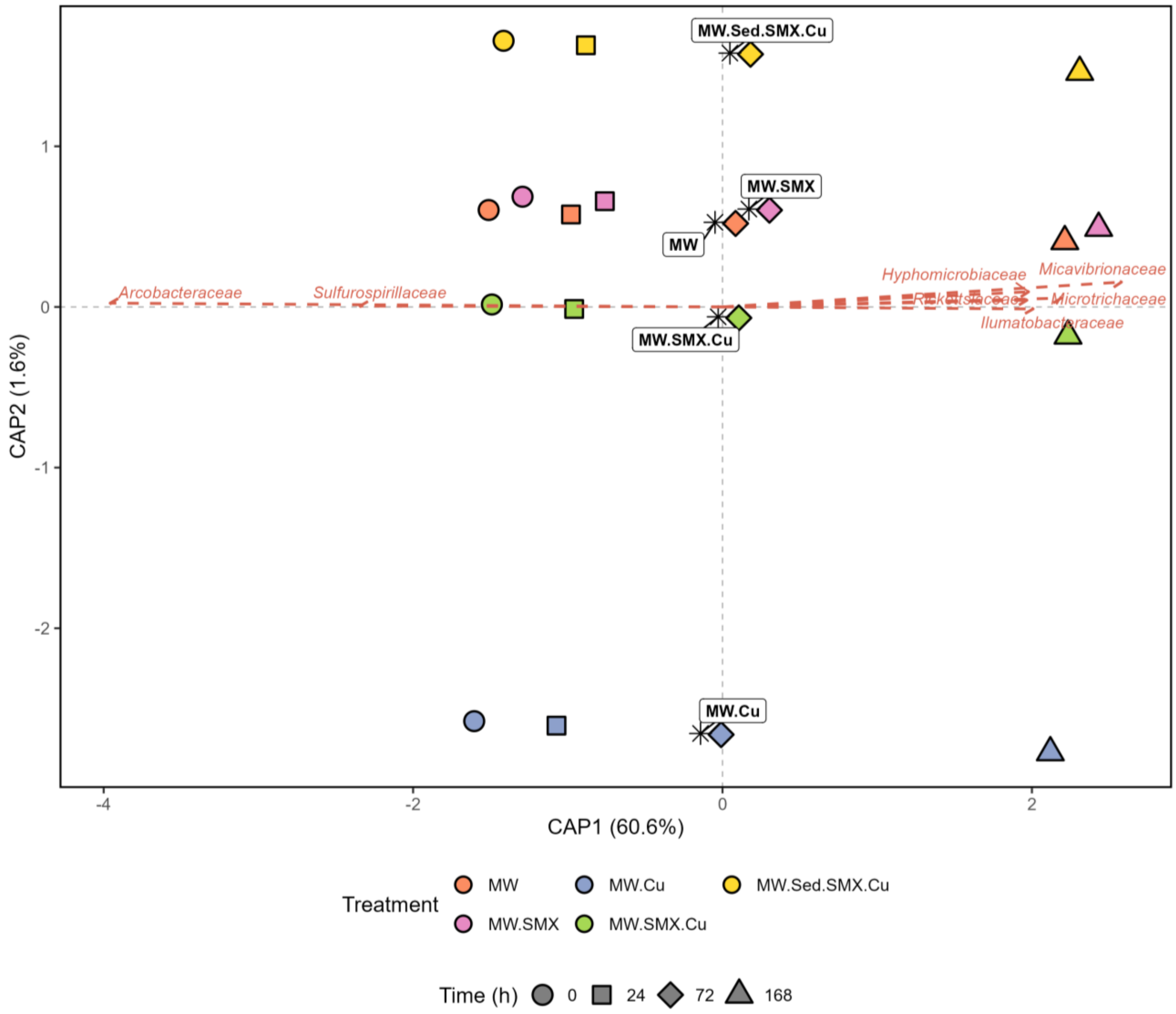


**Supplementary Figure S9.** db-RDA showing time and treatment-associated shifts in family-level bacterial community composition across mesocosms (Community (Family) = Treatment + Time). All mesocosm communities changed along CAP1 over time but not along CAP2, which hardly explained any of the variation. Treatment only differentiated along with this CAP2. Each * indicates the central average position of all time points within a given treatment group (labeled with text boxes like MW, MW.Cu, MW.Sed.SMX.Cu, etc.), showing how treatments cluster relative to one another in the ordination space.

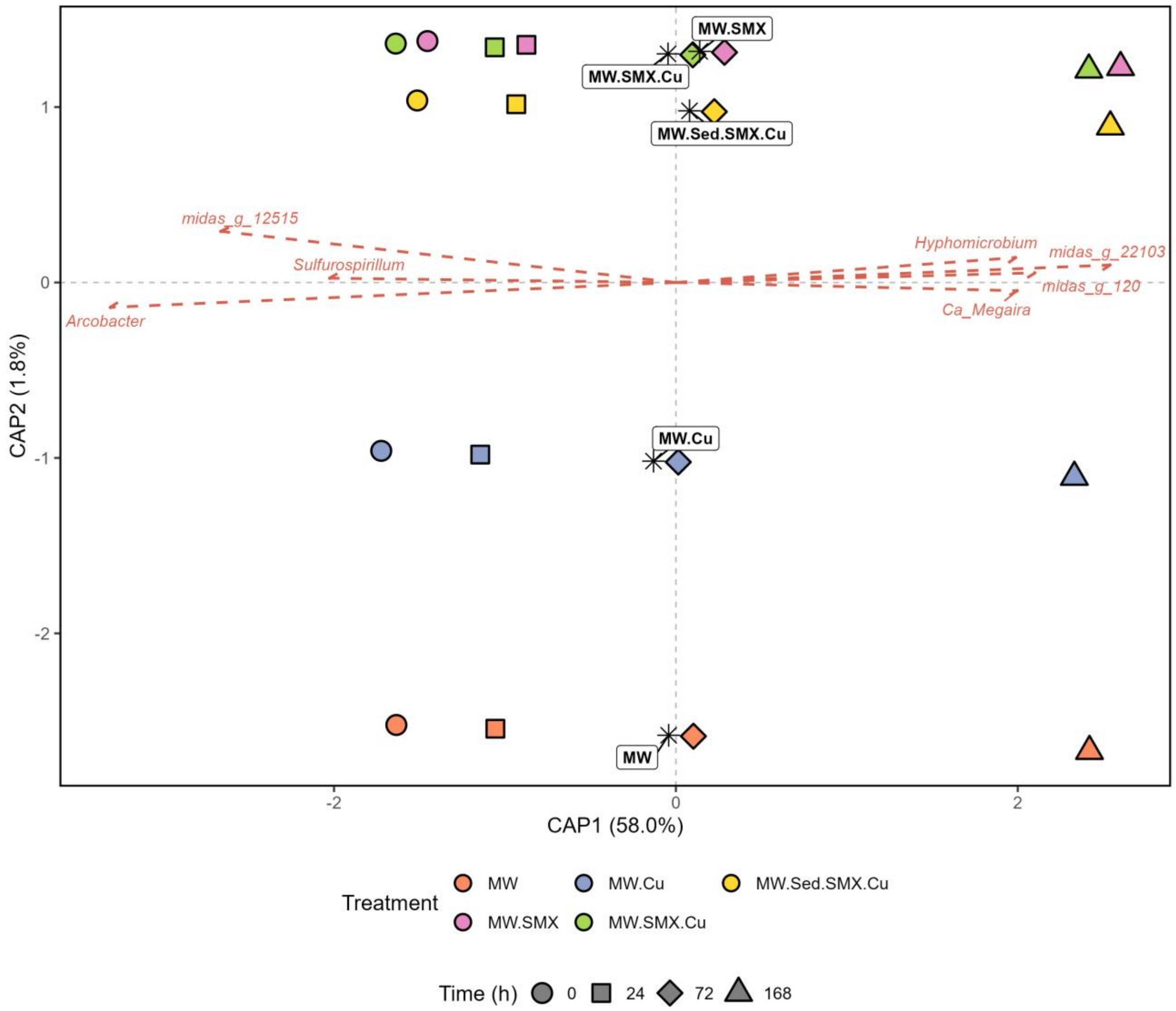


**Supplementary Figure S10.** db-RDA showing time and treatment-associated shifts in genus-level microbial community composition across mesocosm (Community (Genus) = Treatment + Time). All mesocosm communities changed along CAP1 over time but not along CAP2, which hardly explained any of the variation. Treatment only differentiated along with this CAP2. Each * indicates the central average position of all time points within a given treatment group (labeled with text boxes like MW, MW.Cu, MW.Sed.SMX.Cu, etc.), showing how treatments cluster relative to one another in the ordination space.

**Supplementary Table S10.** db-RDA of ARG and MC composition constrained by the four WQPs that changed over time (pH, DO, TUR, COD). Plotted in the **Supplementary Figure S11 and S12.**

| Response | Model | $R^2$ | Adj. $R^2$ | Term | F | p-value |
|---|---|---|---|---|---|---|
| **ARG** | *Full (ARG = WQPs)* | 0.639 | 0.596 | All axes | 15.015 | 0.001 |
| | | | | CAP1 | 28.660 | 0.001 |
| | | | | CAP2 | 1.369 | 0.248 |
| | *Partial (ARG = WQPs \| Time (unique))* | 0.147 | 0.121 | CAP1 | 7.988 | 0.394 |
| | | | | CAP2 | 1.241 | 0.349 |
| **MC (Phylum)** | *Full (Phylum = WQPs)* | 0.727 | 0.695 | All axes | 22.626 | 0.001 |
| | | | | CAP1 | 34.500 | 0.001 |
| | | | | CAP2 | 10.753 | 0.001 |
| | *Partial (Phylum = WQPs \| Time (unique))* | 0.114 | 0.106 | CAP1 | 14.718 | 0.231 |
| | | | | CAP2 | 3.202 | 0.112 |
| **MC (Family)** | *Full (Family = WQPs)* | 0.548 | 0.495 | All axes | 10.312 | 0.001 |
| | | | | CAP1 | 14.667 | 0.001 |
| | | | | CAP2 | 5.957 | 0.005 |
| | *Partial (Family = WQPs \| Time (unique))* | 0.201 | 0.186 | CAP1 | 13.489 | 0.224 |
| | | | | CAP2 | 2.859 | 0.197 |
| **MC (Genus)** | *Full (Genus = WQPs)* | 0.513 | 0.456 | All axes | 8.960 | 0.001 |
| | | | | CAP1 | 12.625 | 0.001 |
| | | | | CAP2 | 5.295 | 0.01 |
| | *Partial (Genus = WQPs \| Time (unique))* | 0.202 | 0.184 | CAP1 | 11.889 | 0.237 |
| | | | | CAP2 | 2.706 | 0.163 |

'Unique' refers to the proportion of total variance explained solely and independently by a given factor (ARGs/Community, Treatment, or Time) after partialling out (removing) all other model covariates.

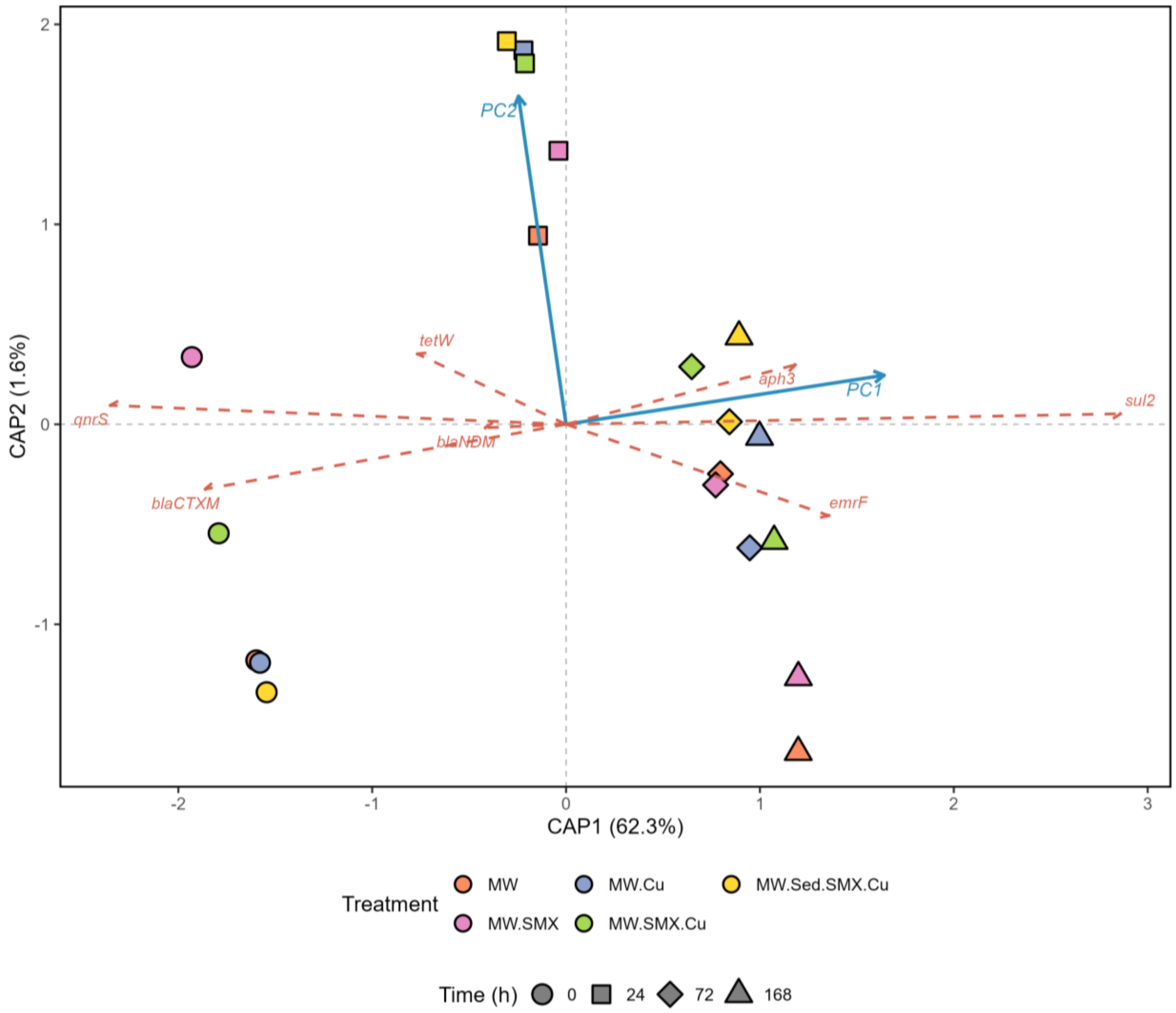


**Supplementary Figure S11.** db-RDA showing the relationship between ARG composition, temporal progression and the four main WQPs (pH, DO, NTU, COD) across mesocosms (ARG = PC1 + PC2, where Aitchison distance (Euclidean distance calculated on Centered Log-Ratio (CLR) transformed ARG relative abundance) and PC1 and PC2 are the top principal components derived from a PCA on the measured WQPs.

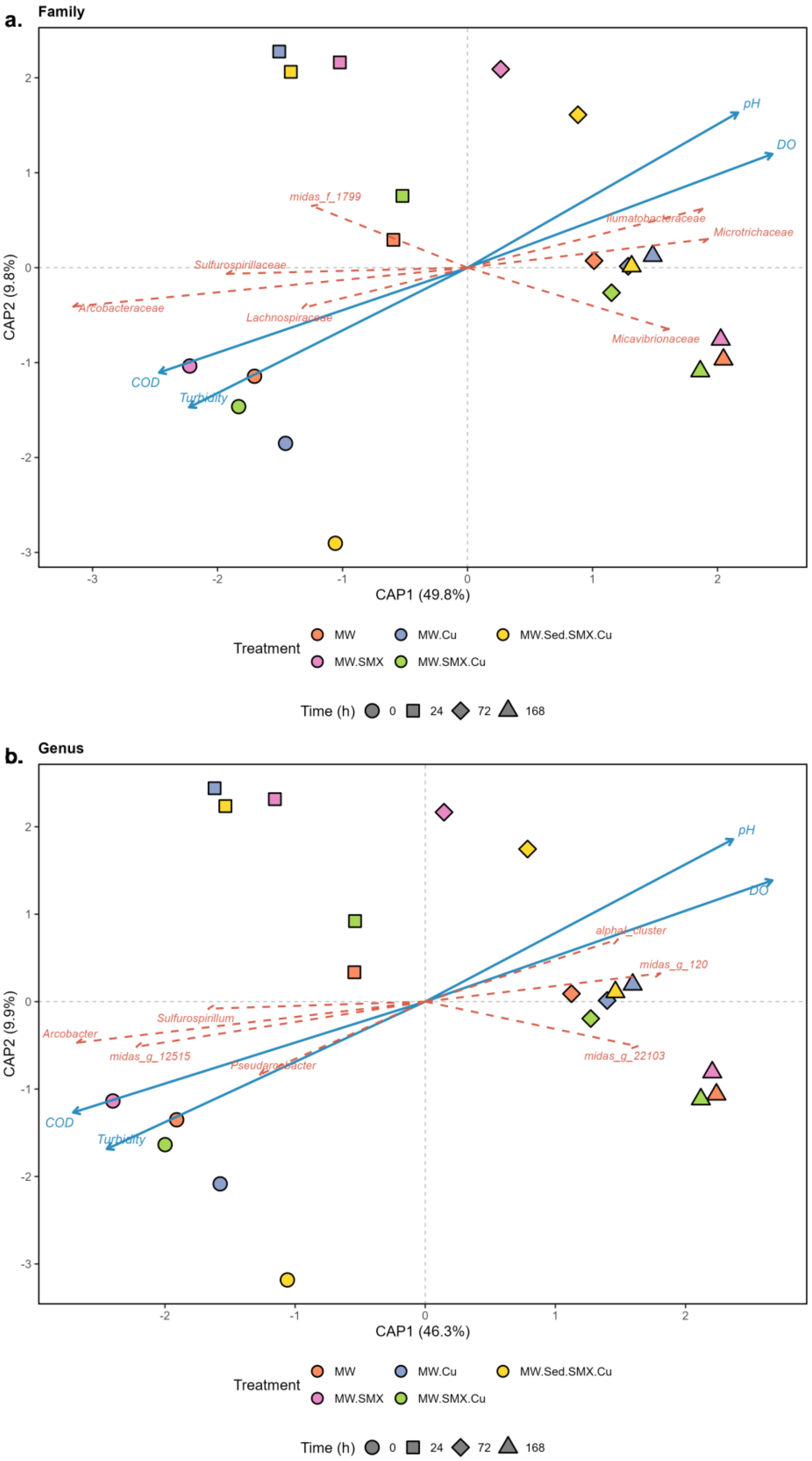


**Supplementary Figure S12.** db-RDA showing the relationship between microbial community composition and the four WQPs that changed over time (pH, DO, NTU, COD) at the (a) family,

(b) genus levels across mesocosms. See **Supplementary Table S11** for the numerical results.

**Supplementary Table S11.** Scores for phylum, family and genus level associated with the constrained ordination axes from the db-RDA analysis of microbial community composition across the mesocosms based on the model MC: Full (MC ~ WQPs) for the four WQPs that changed (pH, DO, TUR, COD).

| Taxon | CAP1 | CAP2 |
|---|---|---|
| **a. Phylum** | | |
| *Cyanobacteriota* | -0.288 | 0.513 |
| *Pseudomonadota* | 0.248 | 0.516 |
| *Bacillota* | -0.088 | 0.022 |
| *Campylobacterota* | -3.053 | -0.306 |
| *Halobacteriota* | -0.279 | 0.243 |
| *Actinobacteriota* | 1.170 | -0.046 |
| *Chloroflexota* | 0.963 | -0.086 |
| *Euryarchaeota* | 0.434 | -0.232 |
| *Desulfobacterota* | 0.558 | -0.351 |
| *Patescibacteriota* | 0.851 | -0.436 |
| *Bdellovibrionota* | -0.249 | 0.208 |
| *Bacteroidota* | -0.267 | -0.045 |
| **b. Family** | | |
| *Arcobacteraceae* | -2.070 | 0.246 |
| *Microtrichaceae* | 1.322 | 0.060 |
| *Ilumatobacteraceae* | 1.299 | -0.272 |
| *Micavibrionaceae* | 0.969 | 0.295 |
| *Lachnospiraceae* | -0.930 | 0.083 |
| *Sulfurospirillaceae* | -1.246 | 0.007 |
| *midas_f_1799* | -0.802 | -0.558 |
| **c. Genus** | | |
| *Arcobacter* | -1.769 | 0.214 |
| *alphaI_cluster* | 1.013 | -0.410 |
| *midas_g_120* | 1.250 | 0.020 |
| *midas_g_22103* | 0.976 | 0.239 |
| *Pseudarcobacter* | -0.880 | 0.535 |
| *midas_g_12515* | -1.482 | 0.284 |
| *Sulfurospirillum* | -1.084 | -0.012 |

**Supplementary Table S12A.** Summary of community–resistome coupling analyses across taxonomic levels based on db-RDA (full and partial), Mantel correlation, and Procrustes superimposition analyses.

| Taxa level | db-RDA (Full) | | | db-RDA (partial) | | | Mantel | | Procrustes | | |
|---|---|---|---|---|---|---|---|---|---|---|---|
| | $R^2$ | Adj. $R^2$ | p | $R^2$ | Adj. $R^2$ | p | r | p | r | $M^2$ | p |
| **Phylum** | 0.813 | 0.704 | 0.001 | 0.145 | 0.154 | 0.006 | 0.771 | 0.001 | 0.841 | 0.292 | 0.001 |
| **Family** | 0.788 | 0.664 | 0.003 | 0.12 | 0.114 | 0.027 | 0.724 | 0.001 | 0.789 | 0.377 | 0.001 |
| **Genus** | 0.787 | 0.663 | 0.001 | 0.119 | 0.113 | 0.019 | 0.704 | 0.001 | 0.779 | 0.394 | 0.001 |

**Supplementary Table S12B.** Partial db-RDA and additive variance partitioning decomposing resistome variation into contributions from bacterial community composition, incubation time, and treatment, at three taxonomic levels.

| Tax level | Component | $R^2$ | Adj. $R^2$ | p-value |
|---|---|---|---|---|
| **Phylum** | Community \| Treatment only (Time retained) | 0.724 | not reportable* | 0.001 |
| | Community \| Time + Treatment (unique) | 0.145 | 0.154 | 0.009 |
| | Time \| Community + Treatment (unique) | 0.019 | 0.005 | 0.324 |
| | Treatment \| Community + Time (unique) | 0.063 | 0.001 | 1 |
| | Confounded overlap (Community x Time x Treatment) | N/A | 0.543 | N/A |
| **Family** | Community \| Treatment only (Time retained) | 0.707 | not reportable* | 0.001 |
| | Community \| Time + Treatment (unique) | 0.12 | 0.114 | 0.028 |
| | Time \| Community + Treatment (unique) | 0.012 | -0.009 | 0.684 |
| | Treatment \| Community + Time (unique) | 0.076 | 0.006 | 1 |
| | Confounded overlap (Community x Time x Treatment) | N/A | 0.553 | N/A |
| **Genus** | Community \| Treatment only (Time retained) | 0.71 | not reportable* | 0.003 |
| | Community \| Time + Treatment (unique) | 0.119 | 0.113 | 0.03 |
| | Time \| Community + Treatment (unique) | 0.007 | -0.015 | 0.824 |
| | Treatment \| Community + Time (unique) | 0.074 | 0.003 | 1 |
| | Confounded overlap (Community x Time x Treatment) | N/A | 0.562 | N/A |

**Supplementary Table S13.** Permutation-based db-RDA model statistics for community–resistome coupling analyses.

| **Taxon** | **Model Term** | **F** | **p-value** |
|---|---|---|---|
| **a. Phylum** | CommPC1 | 40.148 | 0.001 |
| | CommPC2 | 6.788 | 0.028 |
| | Time | 1.181 | 0.202 |
| | Treatment | 1.011 | 0.002 |
| **b. Family** | CommPC1 | 31.514 | 0.002 |
| | CommPC2 | 8.053 | 0.016 |
| | Time | 0.673 | 0.269 |
| | Treatment | 1.068 | 0.018 |
| **c. Genus** | CommPC1 | 29.789 | 0.001 |
| | CommPC2 | 9.847 | 0.005 |
| | Time | 0.537 | 0.437 |
| | Treatment | 1.037 | 0.016 |

**Supplementary Table S14.** Major microbial taxa at each level associated with ARG restructuring based on db-RDA environmental vector fitting analysis.

| Taxa | CAP1 | CAP2 | $R^2$ | p-value | Direction |
|---|---|---|---|---|---|
| **a. Phylum** | | | | | |
| *Actinobacteriota* | 0.874 | 0.002 | 0.763 | 0.003 | Positive |
| *Chloroflexota* | 0.839 | -0.058 | 0.707 | 0.003 | Positive |
| *Euryarchaeota* | 0.728 | -0.253 | 0.594 | 0.007 | Positive |
| *Patescibacteriota* | 0.700 | -0.232 | 0.544 | 0.014 | Positive |
| *Desulfobacterota* | 0.584 | -0.387 | 0.491 | 0.019 | Positive |
| *Bacteroidota* | -0.332 | 0.342 | 0.227 | 0.116 | Negative |
| *Halobacteriota* | -0.489 | 0.431 | 0.425 | 0.019 | Negative |
| *Planctomycetota* | -0.657 | -0.618 | 0.812 | 0.003 | Negative |
| *Acidobacteriota* | -0.792 | -0.141 | 0.646 | 0.003 | Negative |
| *Campylobacterota* | -0.944 | -0.026 | 0.891 | 0.003 | Negative |
| **b. Family** | | | | | |
| *Caldilineaceae* | 0.9073 | 0.0166 | 0.823 | 0.008875 | Positive |
| *Ilumatobacteraceae* | 0.9018 | 0.2325 | 0.867 | 0.008875 | Positive |
| *Beijerinckiaceae* | 0.8555 | 0.1849 | 0.766 | 0.008875 | Positive |
| *Microtrichaceae* | 0.8101 | 0.0087 | 0.656 | 0.008875 | Positive |
| *Phormidiaceae* | 0.8025 | -0.0069 | 0.644 | 0.008875 | Positive |
| *midas_f_50127* | -0.8163 | 0.1666 | 0.694 | 0.008875 | Negative |
| *Family_XI* | -0.8606 | -0.0964 | 0.750 | 0.008875 | Negative |
| *Sulfurimonadaceae* | -0.8617 | -0.2657 | 0.813 | 0.008875 | Negative |
| *Sulfurospirillaceae* | -0.8963 | 0.0513 | 0.806 | 0.008875 | Negative |
| *Arcobacteraceae* | -0.9376 | 0.0063 | 0.879 | 0.008875 | Negative |
| **c. Genus** | | | | | |
| *alphaI_cluster* | 0.949 | 0.103 | 0.911 | 0.017 | Positive |
| *midas_g_1965* | 0.874 | 0.044 | 0.766 | 0.017 | Positive |
| *Romboutsia* | 0.865 | 0.112 | 0.760 | 0.017 | Positive |
| *midas_g_34* | 0.855 | -0.039 | 0.732 | 0.017 | Positive |
| *Planktothrix_NIVA-CYA_15* | 0.850 | -0.010 | 0.722 | 0.017 | Positive |
| *Pseudarcobacter* | -0.869 | -0.281 | 0.833 | 0.017 | Negative |
| *Subdoligranulum* | -0.870 | -0.201 | 0.798 | 0.017 | Negative |
| *Sulfurospirillum* | -0.891 | 0.039 | 0.795 | 0.017 | Negative |
| *midas_g_12515* | -0.922 | -0.041 | 0.852 | 0.017 | Negative |
| *Arcobacter* | -0.942 | -0.030 | 0.888 | 0.017 | Negative |

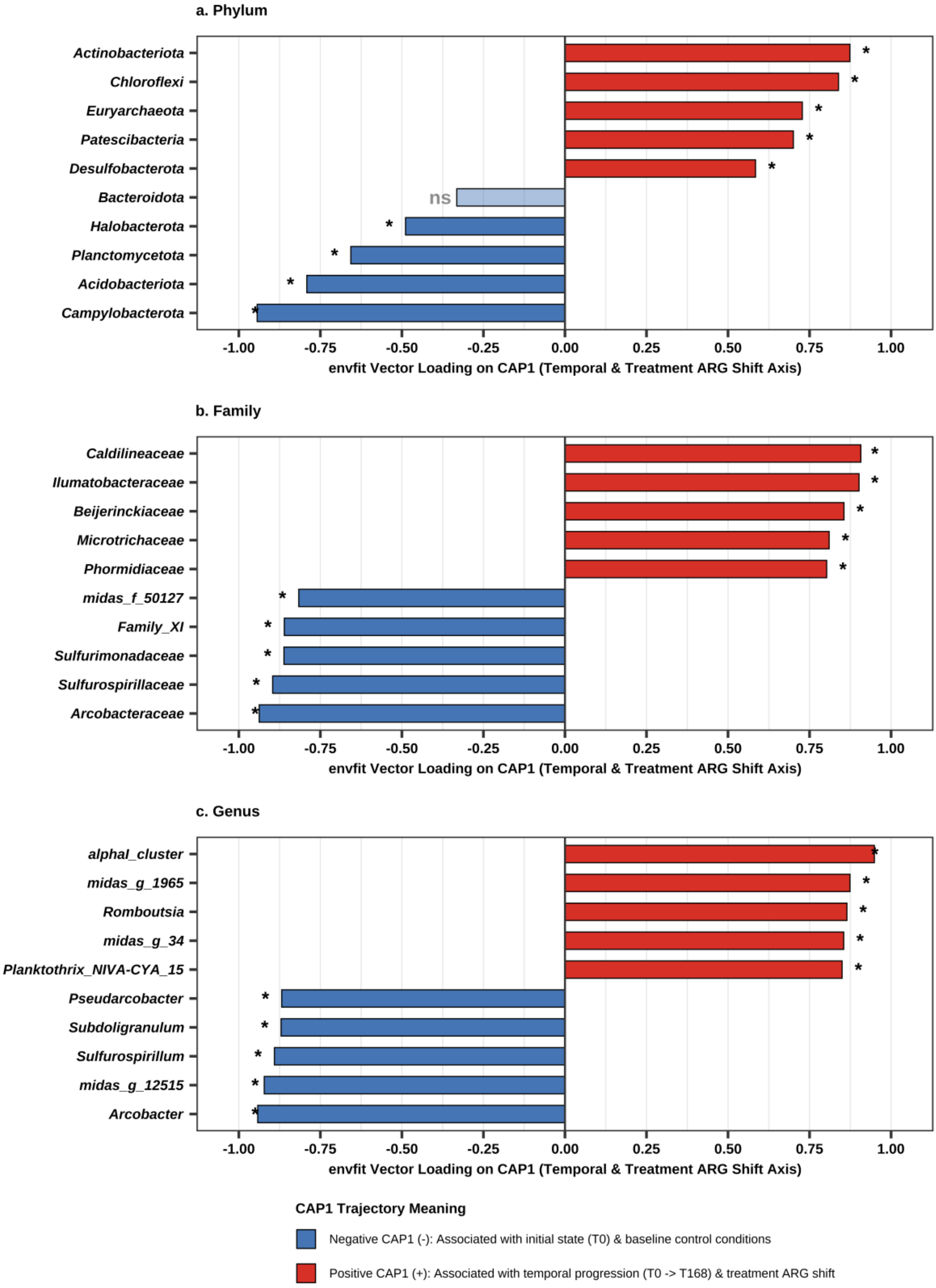


**Supplementary Figure S13.** Major microbial taxa associated with ARG restructuring gradients identified by db-RDA vector fitting analyses at the phylum, family, and genus levels. Positive and negative bar directions indicate associations along the dominant constrained ordination axis (CAP1), and asterisks denote significant taxa after Benjamini–Hochberg correction (adjusted $p < 0.05$). In distance-based RDA (ARG= Common PCs + Time + Treatment), CAP1 represents the primary constrained axis of ARG profile variation. Positive CAP1 loading indicates taxa whose CLR relative abundance increases over time during mesocosm succession and/or in response to treatment disturbance (co-occurring with the shifted ARG profile). Negative CAP1 loading indicates taxa associated with the initial baseline state ($T_0$) conditions.

**Supplementary text 2 for additional db-RDA, Procrustes and Mantel test analysis:**

For ARG composition, the full WQP model explained a substantial fraction of variation ($R^2$ = 0.639; adjusted $R^2$ = 0.596, **Figure 7; Supplementary Table S10 and Supplementary Figure S11**), with the primary constrained axis identified as highly significant (F = 28.66, p = 0.001). However, after conditioning for temporal effects, explanatory power declined markedly (Adj. $R^2$ = 0.121), and neither constrained axis remained statistically significant (PC1: F = 7.99, p = 0.355; PC2: F = 1.24, p = 0.307).

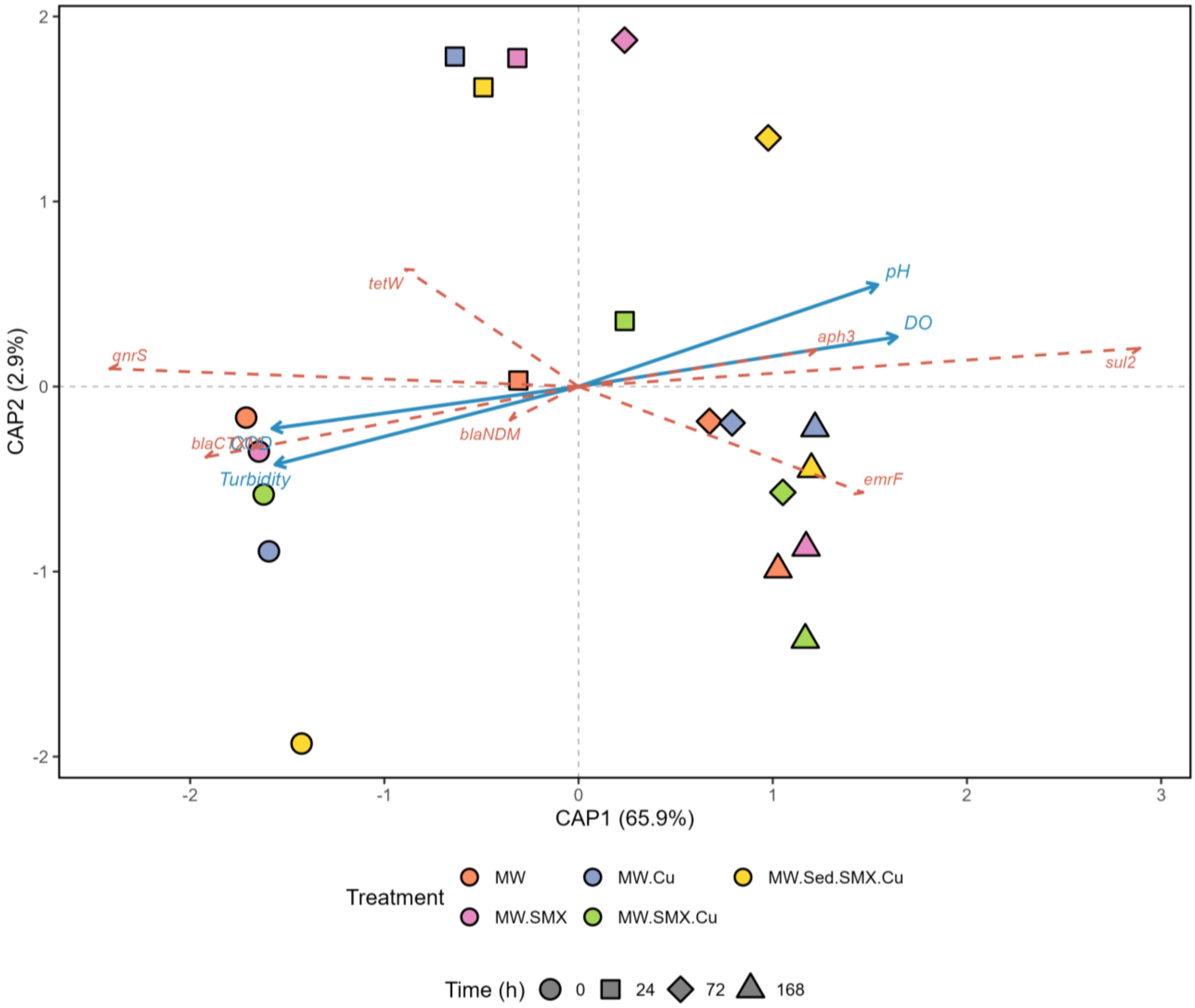


**Supplementary Figure S14.** db-RDA ordination of ARG composition constrained by WQPs showing strong coupling between resistome restructuring and physicochemical changes during mesocosm incubation.

A comparable pattern was observed for phylum-level community composition (**Figure 7, Supplementary Table S10**). The full WQP model explained a large proportion of community variation ($R^2 = 0.791$; adjusted $R^2 = 0.739$), with the first constrained axis identified as highly significant ($F = 53.03$, $p = 0.001$). After conditioning on time, the independent explanatory contribution of WQPs decreased substantially (Adj. $R^2 = 0.233$), although the first constrained axis remained statistically significant ($F = 13.35$, $p = 0.017$).

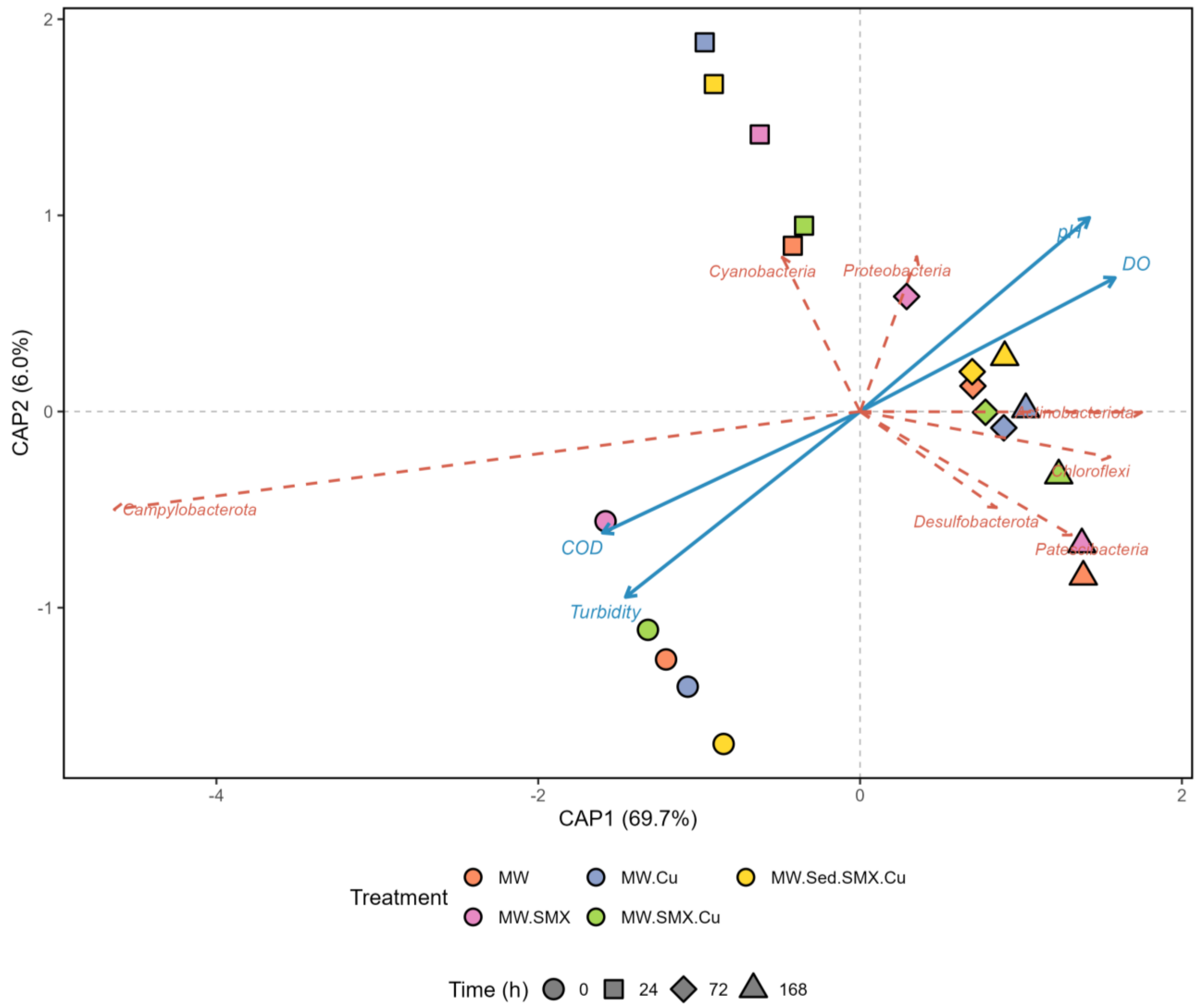


**Supplementary Figure S15.** db-RDA ordination of phylum-level bacterial community composition constrained by water quality parameters showing strong association between microbial succession and physicochemical changes across mesocosm treatments.

These findings indicate that physicochemical gradients were closely associated with both resistome and bacterial community restructuring (**Supplementary Figure S14 and Supplementary Figure S15**).

Procrustes superimposition analysis (**Supplementary Table S12 and Supplementary Figure S16**) further showed strong concordance between microbial community and resistome ordination patterns. The strongest concordance was observed at the phylum level (Procrustes r = 0.841, p = 0.001), followed by family-level (r = 0.789, p = 0.001) and genus-level ordinations (r = 0.779, p = 0.001). These high concordance values indicate that changes in microbial community composition were closely linked to resistome restructuring.

Mantel analysis (**Supplementary Table S12 and Supplementary Figure S17**) further supported the association between microbial community succession and resistome restructuring. Significant positive correlations were observed between bacterial community dissimilarity and ARG compositional dissimilarity at all taxonomic levels examined, suggesting but not proving genetic linkage (**Supplementary Table S12**). The strongest correlation was detected at the phylum level (Mantel r = 0.772, p = 0.001), followed by the family (r = 0.724, p = 0.001) and genus levels (r = 0.704, p = 0.001).

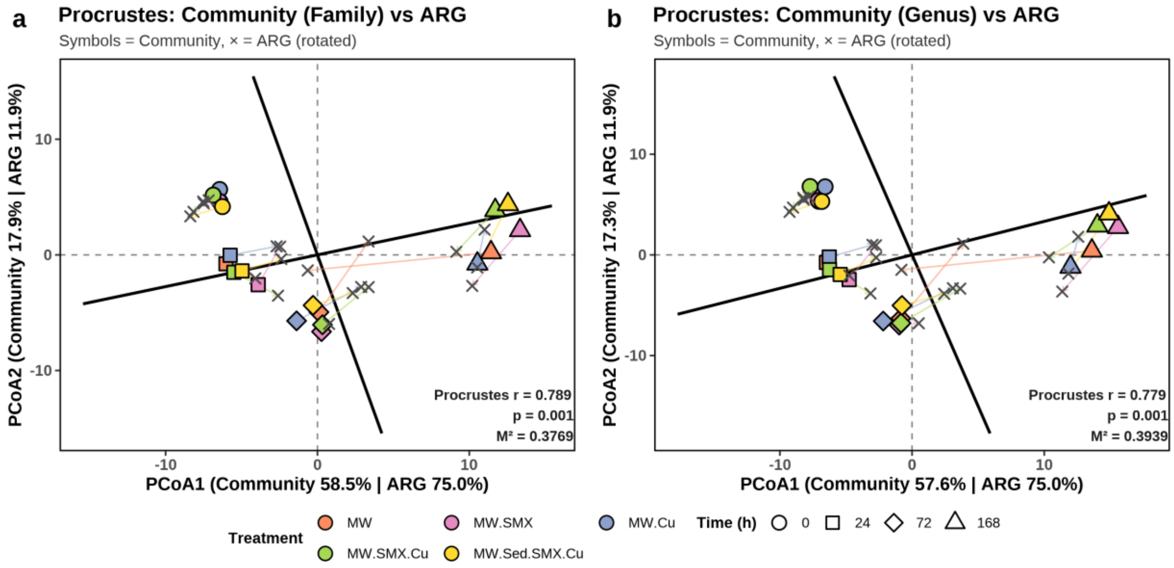


**Supplementary Figure S16.** Procrustes superimposition analyses showing concordance between microbial community (MC) composition and ARG profiles across mesocosm treatments. (a) Family-level MC vs. ARG profiles. (b) Genus-level MC vs. ARG profiles. MC PCoA coordinates are represented by filled shapes (color = treatment, shape = time in hours). Rotated ARG ordination coordinates are marked by grey crosses. Line segments connect each microbial community sample to its corresponding ARG profile; shorter segment lengths denote stronger structural concordance ($M^2$ = Procrustes sum-of-squared residuals, where lower $M^2$ and higher Procrustes r indicate stronger alignment significance, 999 permutations).

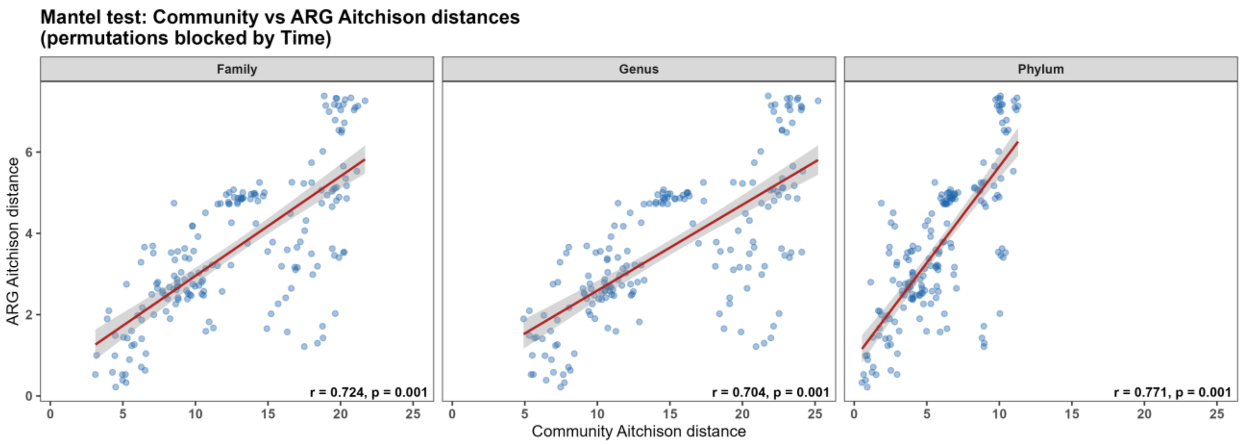


**Supplementary Figure S17.** Mantel correlations between microbial community and ARG Aitchison distance matrices across phylum, family, and genus levels.

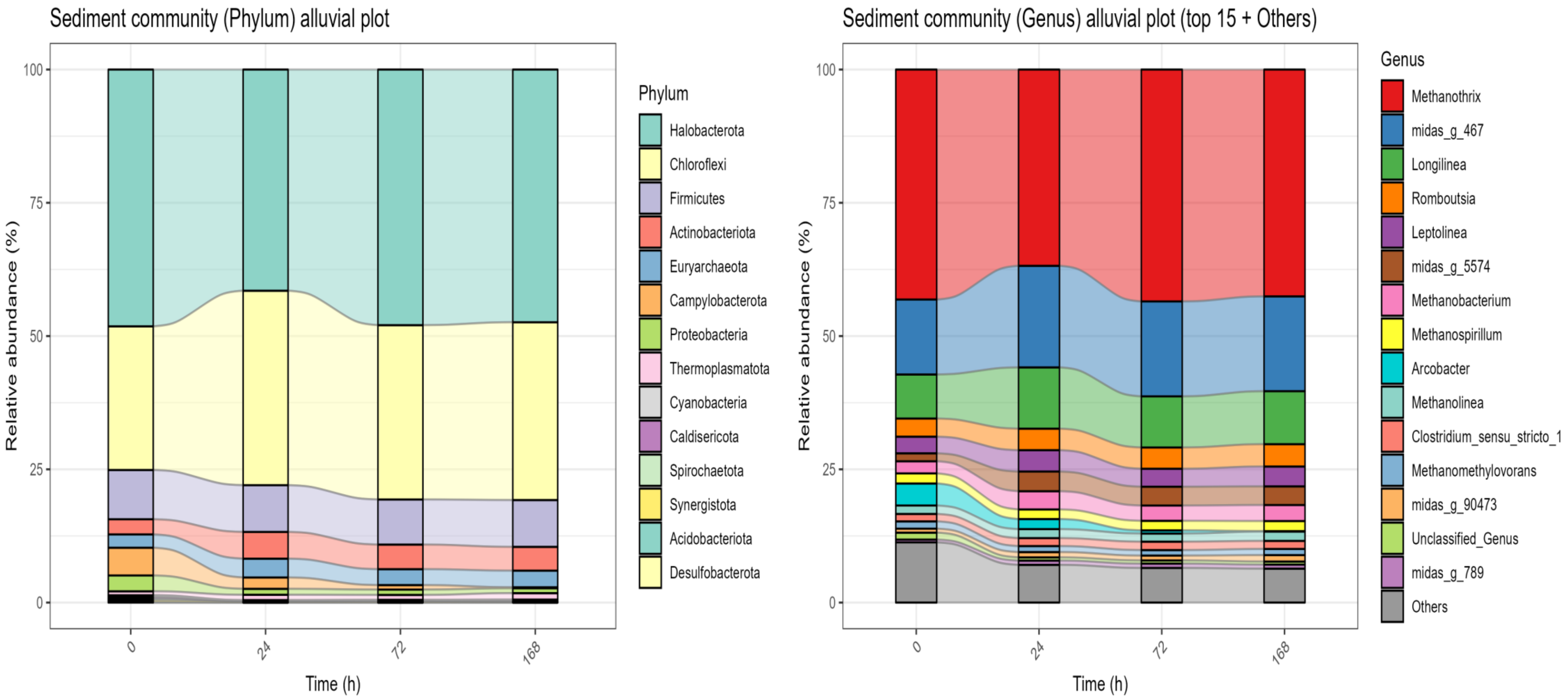


**Supplementary Figure S18.** . Bar plot of microbial community relative abundance across treatments and time points in the sediment samples on (a) Phylum level, (b) Genus's level.

### References


Avramidis, P., Bekiari, V., 2021. Application of a catalytic oxidation method for the simultaneous determination of total organic carbon and total nitrogen in marine sediments and soils. PLoS One 16, e0252308. https://doi.org/10.1371/JOURNAL.PONE.0252308

AWWA APHA, 2017. Standard methods for the examination of water and wastewater. cir.nii.ac.jp.

Barraud, O., Baclet, M.C., Denis, F., Ploy, M.C., 2010. Quantitative multiplex real-time PCR for detecting class 1, 2 and 3 integrons. Journal of Antimicrobial Chemotherapy 65, 1642–1645. https://doi.org/10.1093/JAC/DKQ167

Frahm, E., Obst, U., 2003. Application of the fluorogenic probe technique (TaqMan PCR) to the detection of Enterococcus spp. and Escherichia coli in water samples. J. Microbiol. Methods 52, 123–131. https://doi.org/10.1016/S0167-7012(02)00150-1

Liang, C., Wei, D., Zhang, S., Ren, Q., Shi, J., Liu, L., 2021. Removal of antibiotic resistance genes from swine wastewater by membrane filtration treatment. Ecotoxicol. Environ. Saf. 210, 111885. https://doi.org/10.1016/J.ECOENV.2020.111885

Marti, E., Balcázar, J.L., 2013a. Real-time PCR assays for quantification of qnr genes in environmental water samples and chicken feces. Appl. Environ. Microbiol. 79, 1743–1745. https://doi.org/10.1128/AEM.03409-12;JOURNAL:JOURNAL:AM;WGROUP:STRING:PUBLICATION

Marti, E., Balcázar, J.L., 2013b. Real-time PCR assays for quantification of qnr genes in environmental water samples and chicken feces. Appl. Environ. Microbiol. 79, 1743–1745. https://doi.org/10.1128/AEM.03409-12;JOURNAL:JOURNAL:AM;WGROUP:STRING:PUBLICATION

Muyzer, G., De Waal, E.C., Uitterlinden, A.G., 1993. Profiling of complex microbial populations by denaturing gradient gel electrophoresis analysis of polymerase chain reaction-amplified genes coding for 16S rRNA. Appl. Environ. Microbiol. 59, 695–700. https://doi.org/10.1128/AEM.59.3.695-700.1993;ISSUE:ISSUE:DOI

Pei, R., Kim, S.C., Carlson, K.H., Pruden, A., 2006. Effect of River Landscape on the sediment concentrations of antibiotics and corresponding antibiotic resistance genes (ARG). Water Res. 40, 2427–2435. https://doi.org/10.1016/J.WATRES.2006.04.017

Silkie, S.S., Tolcher, M.P., Nelson, K.L., 2008. Reagent decontamination to eliminate false-positives in Escherichia coli qPCR. J. Microbiol. Methods 72, 275–282. https://doi.org/10.1016/J.MIMET.2007.12.011

Thomas, D.H., Rey, M., Jackson, P.E., 2002. Determination of inorganic cations and ammonium in environmental waters by ion chromatography with a high-capacity cation-exchange column. J. Chromatogr. A 956, 181–186. https://doi.org/10.1016/S0021-9673(02)00141-3

Walsh, F., Ingenfeld, A., Zampicolli, M., Hilber-Bodmer, M., Frey, J.E., Duffy, B., 2011. Real-time PCR methods for quantitative monitoring of streptomycin and tetracycline resistance genes in agricultural ecosystems. J. Microbiol. Methods 86, 150–155. https://doi.org/10.1016/J.MIMET.2011.04.011
